# Proteome-wide quantification of protein turnover in frog and fly embryos reveals divergent strategies of maternal inheritance

**DOI:** 10.64898/2026.09.28.754234

**Authors:** Edward R. Cruz, Argit Marishta, Gloria Bao, Alexander N.T. Johnson, Felix C. Keber, Vyas Pujari, Daniil Ivanov, Michael Neinast, Marc W. Kirschner, Joshua D. Rabinowitz, Eric Wieschaus, Martin Wühr

## Abstract

Every embryo inherits a maternal proteome that it must remodel with zygotic proteins to build its many cell types. The fate of the maternal proteome remains contested because indirect measurements cannot resolve it. Here, we combine ^18^O-water labeling with multiplexed proteomics to quantify protein turnover proteome-wide in frog and fly embryos. Through hatching, the frog preserves the bulk of its maternal proteome, confining rapid degradation to a small regulatory module. The fly cannot meet its synthesis demand from yolk alone and instead degrades nearly all maternal proteins, including housekeeping proteins long assumed stable, recycling them into new protein. Yet the turnover hierarchy is conserved, with disordered and regulatory proteins degrading fastest, while the fly rescales the whole proteome∼eightfold faster. These results recast the developmental proteome as both informational inheritance and metabolic reserve, establish ^18^O-water labeling as a turnover method for non-feeding organisms, and provide a resource of embryonic half-lives.

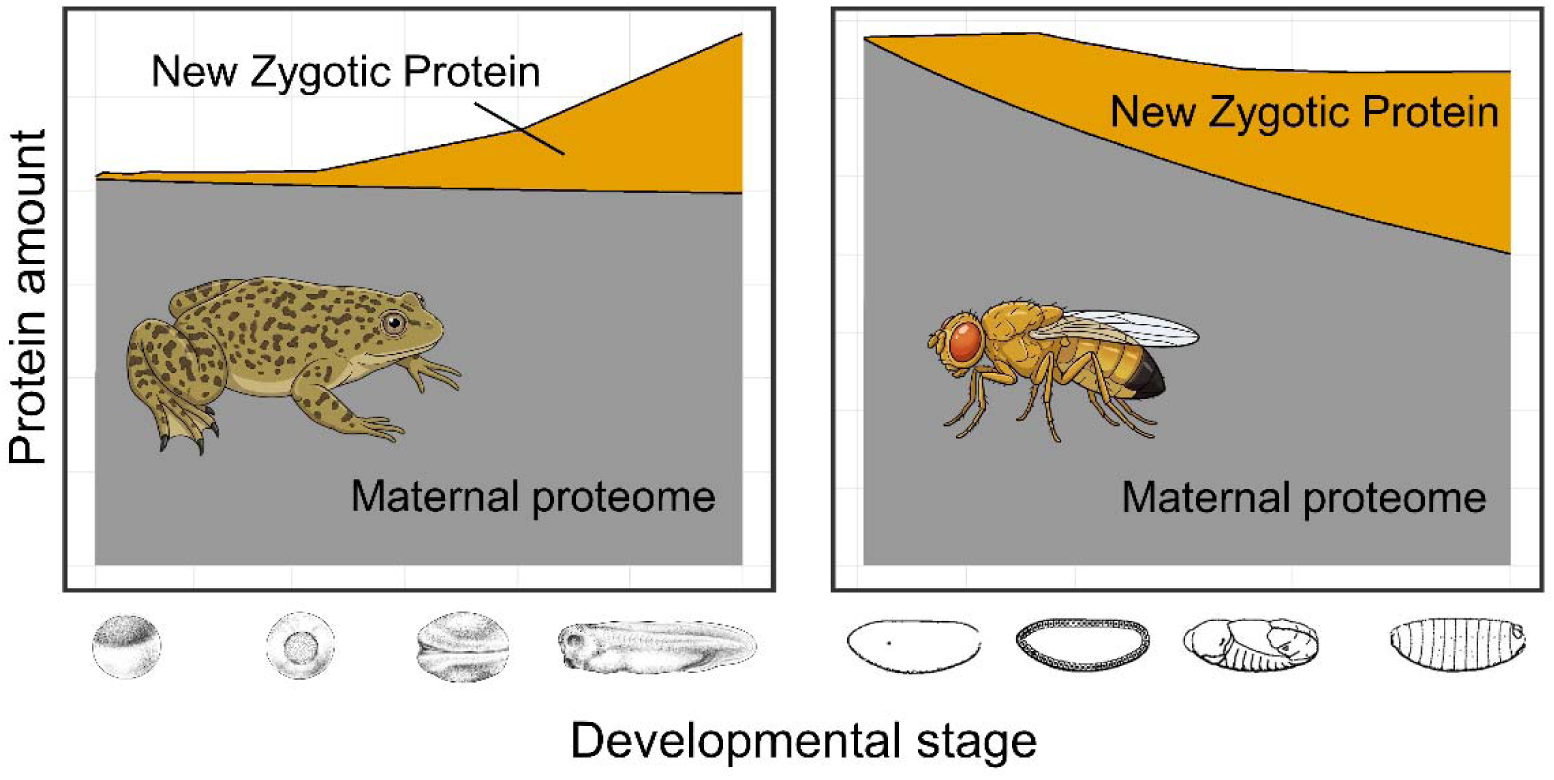

## Introduction

Every embryo begins as a single cell stocked by its mother, with the father contributing little beyond a genome and a centrosome^1,2^. In externally developing embryos such as frogs and flies, which take up no nutrients before feeding, this maternal endowment is the entire supply of building blocks for early development. How it is remodeled into the embryonic body is a fundamental question in developmental biology.

That question has been addressed overwhelmingly through mRNA. The maternal-to-zygotic transition (MZT) is itself defined by transcript clearance, and the pathways that execute it are well characterized. A substantial but species- and stage-dependent fraction of maternal transcripts is selectively destabilized through temporally separable maternal and zygotic decay activities^3–5^, specified by RNA-binding proteins such as Smaug and Brat in *Drosophila melanogaster*^6–8^, and by microRNAs including the miR-309 cluster in Drosophila^9^ and miR-427 in *Xenopus laevis*^10^.

Transcripts, however, are a proxy for the proteins that carry out development, and the fidelity of that proxy is set by protein half-life. Degrading a maternal mRNA halts new synthesis, but it removes the encoded protein only if that protein turns over on a timescale comparable to development. A protein whose half-life exceeds the developmental window persists largely unchanged long after its transcript is gone, so its clearance has little proteomic consequence within the embryo. Protein half-lives are therefore the parameter required to translate transcript measurements into statements about the molecules that do the work.

Consistent with this, mRNA and protein correspond poorly in embryos. In Xenopus and in the chordate *Ciona robusta*, transcript levels are weak predictors of protein abundance, with protein-level changes damped or delayed relative to transcript dynamics^11,12^. Translational control accounts for part of the discordance. Poly(A)-tail length is strongly coupled to translational efficiency before gastrulation, and egg ribosomes in frog and fish are held dormant by conserved factors released during early embryogenesis^13,14^. The other term, degradation, remains unquantified, so the discordance cannot be decomposed.

The obstacle is that protein abundance, the standard observable in developmental proteomics, cannot reveal turnover. Foundational studies have defined how total protein levels change across embryogenesis^11,15,16^, yet abundance alone cannot distinguish a protein that is stable from one held constant by balanced synthesis and degradation. While the clearance of specific maternal proteins is known to be required for orderly developmental transitions^17–20^, far less is established for the proteome as a whole. The available inferences even point in opposite directions. In Drosophila, translationally upregulated mRNAs often fail to yield corresponding protein increases, read as synthesis balanced by degradation^21,22^. In Xenopus, protein abundance changes little from fertilization to hatching, read as a broadly stable maternal proteome^11,23^. In both species, neither conclusion rests on a direct measurement.

Measuring turnover requires labeling either the pre-existing or the newly synthesized protein pool^24,25^, and coupling metabolic labeling to mass spectrometry has extended this proteome-wide across cell lines, primary cells, microbes, mouse oocytes, and adult tissues^26–32^. In stem-cell-derived progenitors, the same approach has shown that human proteins turn over more slowly than their mouse counterparts, linking protein stability to the pace of development^33–35^. The primary challenge is that none of these approaches work in a non-feeding embryo. Dietary labeling is unavailable, injected amino acids must displace large endogenous precursor pools^36^, D_2_O reports only *de novo* synthesis of non-essential amino acids and perturbs cleavage and patterning at useful enrichments^37–39^, and bioorthogonal labels, such as L-Azidohomoalanine (AHA), report new synthesis rather than loss of the pre-existing pool^40,41^.

Here, we introduce ^18^O-water labeling to measure protein turnover in intact embryos by mass spectrometry. We show that ^18^O is rapidly incorporated into the free amino acid pool and subsequently into newly synthesized proteins, while pre-existing proteins remain unlabeled. Tracking decay of the unlabeled signal therefore reports directly on loss of the pre-existing pool, allowing us to quantify turnover for ∼9,600 proteins in Xenopus at the two-cell and gastrulation stages and ∼5,900 proteins in Drosophila at gastrulation. We find that the two embryos pursue divergent regulatory strategies. Xenopus preserves the bulk of its maternal proteome through hatching, whereas Drosophila degrades nearly all of it to varying extents, including many canonical structural and metabolic proteins long assumed to be stable.

Despite this divergence, the functional architecture of turnover is conserved between the species, globally rescaled to Drosophila’s faster tempo.

## Results

### 18O-water labeling proteomics enables quantification of protein turnover in embryos

To measure protein turnover directly in intact embryos via mass spectrometry, we reasoned that we could label the free amino acid pool by bathing embryos in 0.1X MMR prepared with H_2_^18^O. In theory, as pre-existing proteins are hydrolyzed, ^18^O from water is incorporated into the carboxyl group of each liberated amino acid, progressively labeling the free amino acid pool (Figure 1a). This exchange was exploited half a century ago to estimate bulk protein turnover in sporulating bacteria from the ^18^O content of hydrolyzed amino acids^42^, but has not been combined with peptide-resolved mass spectrometry. Newly synthesized proteins then draw on this pool and accumulate ^18^O, while pre-existing proteins remain unlabeled (Figure 1b).

**Figure 1.**
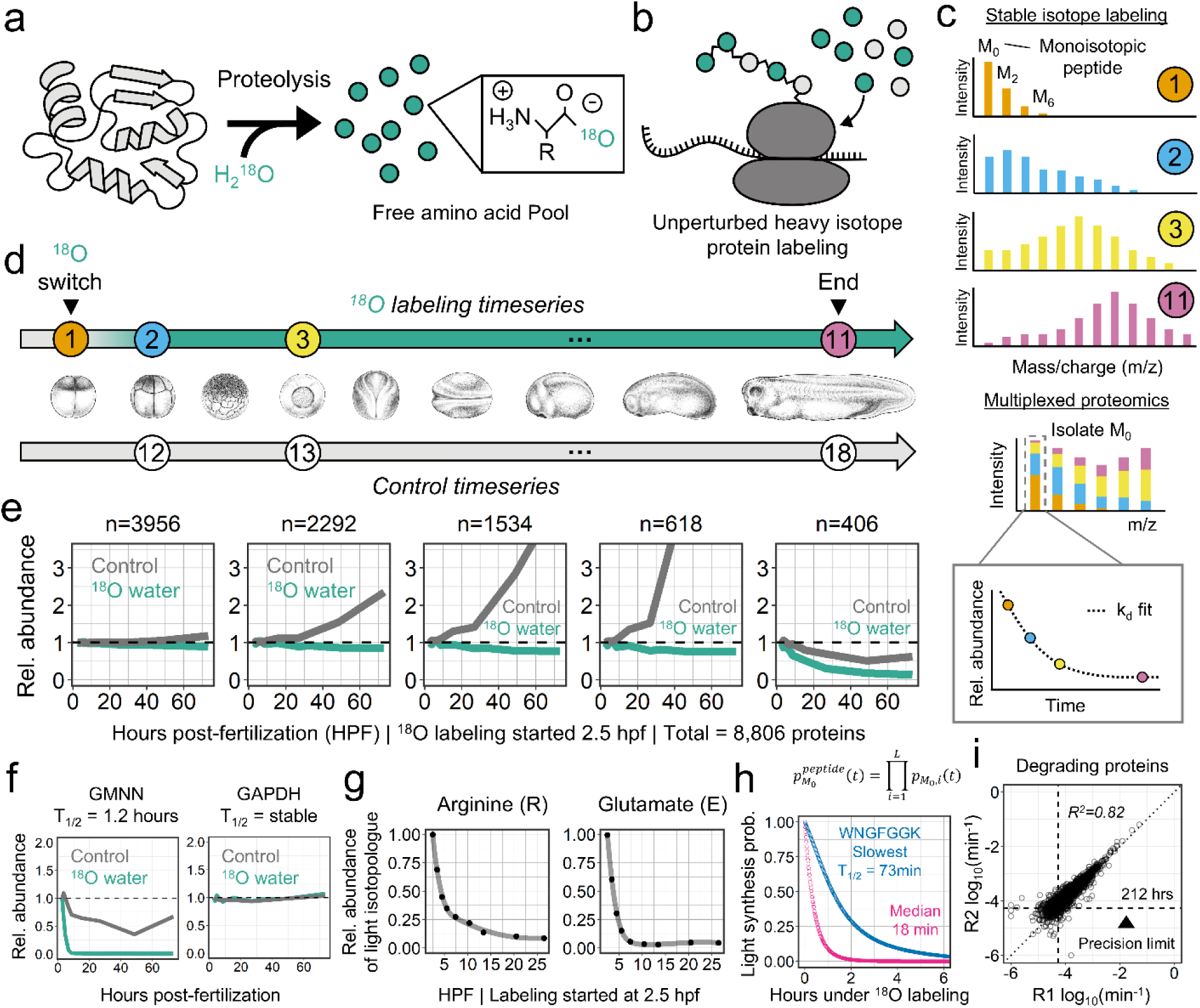
Quantifying protein turnover in intact embryos by ^18^O-water labeling. **(a, b)** Rationale for ^18^O metabolic labeling. Pre-existing proteins are hydrolyzed in H_2_^18^O, transferring an ^18^O atom from water to the carboxyl group of each liberated amino acid. The progressively labeled free amino acid pool is then used by ribosomes to synthesize new proteins, which therefore accumulate ^18^O while pre-existing proteins remain unlabeled. The unlabeled (monoisotopic, M_0_) peptide signal reports on the pre-existing pool. **(c)** ^18^O quantification strategy. A schematic of how peptide isotopic envelopes shift over time as ^18^O is progressively incorporated into newly synthesized proteins and their derived tryptic peptides (top). The monoisotopic peak (M_0_), corresponding to the unlabeled pre-existing peptide population, is isolated and quantified across the time course using multiplexed acquisition (bottom). Decay of the M_0_ signal reflects loss of the pre-existing protein pool and is used to estimate degradation rates (k_d_). **(d)** Experimental design. *Xenopus laevis* embryos were transferred to H_2_^18^O after the first cleavage to initiate labeling and sampled at 11 time points across early development through ∼73 hours post-fertilization (hatching) to generate a labeling time series (top)^46^. A parallel control series of 7 time points acquired in normal water across the same window was used to quantify changes in total protein abundance (bottom). All 18 samples were combined for multiplexed quantification using TMTpro. **(e)** Protein *k*-means clustering of the combined abundance and ^18^O-labeling data. Five resolved clusters are shown with three major trends: proteins with no measurable change in either channel, proteins that accumulate in total abundance while their pre-existing pool decays, and a set declining in both. Control channels (grey) report total protein abundance. ^18^O channels (green) report decay of the pre-existing protein pool. **(f)** Representative protein trajectories. Control channels (grey) report total protein abundance; ^18^O channels (green) track decay of the pre-existing protein pool. Examples illustrate rapid degradation (GMNN, left) and no measurable degradation (GAPDH, right). **(g)** Free amino acid labeling dynamics measured by orthogonal metabolomics following the ^18^O switch. Representative trajectories illustrate the time-dependent decay of the unlabeled fraction for individual amino acids. **(h)** Modeling the precursor labeling lag. From the measured free amino acid labeling curves (Figure S2), we calculated the time-dependent fraction of newly synthesized peptides that are fully unlabeled (light) for each detected sequence. The median tryptic peptide halves within 18 minutes of label exposure; the slowest-labeled peptide observed halves within 73 minutes. **(i)** Reproducibility of degradation rate estimates across biological replicates. Only proteins classified as degrading (Figure S3) are shown, and inter-replicate agreement among them is high (R2 = 0.82). Dashed line marks the precision threshold of 212 hours, three times the ∼71-hour labeling window and approximately the maximum half-life we can reliably resolve. Beyond it, minimal decay prevents precise rate estimation.

As labeling proceeds, ^18^O incorporation shifts each peptide’s isotopic envelope toward higher m/z away from the monoisotopic (M_0_) peak (Figure 1c, top). Reconstructing this shifting envelope at every time point is impractical and yields poor fits, so we instead combine the timepoints into a single multiplexed TMTpro set^43^, isolate the M_0_ signal across all channels, and quantify it by real-time-search MS3 (RTS-MS3)^44,45^. (Figure 1c, bottom). Because all channels are acquired together, each peptide has no missing values, and the decay of the unlabeled M_0_ peak across the multiplexed channels directly reports turnover of the pre-existing pool^32^.

We first applied this to Xenopus, transferring viable embryos into H_2_^18^O shortly after the first cleavage (2.5 hours post-fertilization, 16°C). For technical reasons, labeling cannot begin at fertilization, so the pre-existing pool we follow is the proteome present at the 2-cell stage. Because very few proteins change in abundance between fertilization and first cleavage^23^, for simplicity we refer to this pre-existing pool as the maternal proteome throughout. We sampled 11 time points over the following ∼71 hours (hatching). A parallel control series of 7 time points in normal water tracked total protein abundance (Figure 1d). Wide isolation normalization, that estimates overall channel abundance, was applied to correct for sample-handling variability (Figure S1, Table S1). A *k*-means clustering of the 8,806 quantified proteins (Table S2) visualizes the proteome’s kinetic behaviors, from proteins with no measurable change in either channel, through proteins that accumulate in total abundance while their pre-existing pool decays, to a small set declining in both (Figure 1e). Most proteins fall in the dominant cluster, flat in both channels, where abundance alone would call them stable, and the labeling channel confirms genuinely low turnover rather than the balanced synthesis and degradation it would otherwise mask. At the kinetic extremes, the cell-cycle regulator Geminin (GMNN), rapidly destroyed in early cleavages, and the housekeeping enzyme GAPDH behave as expected, confirming that the method resolves turnover across its full dynamic range (Figure 1f).

To turn M_0_ decay into a degradation rate, we need to know how quickly newly synthesized protein becomes labeled. Until the free amino acid pool is sufficiently labeled, the existing amino acids feeding new synthesis are still partly unlabeled, so new protein itself can add to the M_0_ signal, where it reads as pre-existing protein that has not decayed. We therefore measured the pool directly by metabolomics over the same window (Figure 1g; Figure S2). The maximum number of 18O atoms each amino acid incorporated revealed the route by which it acquired label. The two backbone carboxyl oxygens are labeled in every amino acid, while glutamate and aspartate reach four through their side chains. Side-chain labeling varies by amino acid, and threonine and serine demonstrate a clear distinction. Both carry a side-chain hydroxyl, yet only serine labels it. Serine is made *de novo*, while threonine is essential and has no route to label its side chain^47^ (Figure S2). This dependence on metabolic origin is consistent with ^18^O entering through biosynthesis as well as proteolysis, which validates the measurement and shows the labeling is more complicated than our initial hypothesis and the carboxyl-only scheme shown in Figure 1a. Importantly, the additional metabolomics information allows us to impute each peptide’s light-synthesis probability empirically from the measured curves rather than from a flux model of the precursor pools^32^.

The light synthesis probability is the chance that a peptide is fully unlabeled if it was synthesized (Figure 1h, Supplementary Methods). For the median tryptic peptide, this reaches 50% within 18 minutes of the switch, and even the slowest peptide-labeling halves within 73 minutes (Figure 1h). This window is short relative to the degradation timescales of most proteins we measure, and the labeling lag sets a rough floor on the shortest half-lives we can confidently measure. At later times, newly made proteins are essentially never unlabeled and the M_0_ signal reflects degradation alone.

We modeled the turnover of each protein with a single mass-balance framework^32^. In our model, the rate of change of M_0_ reflects synthesis of new light peptide from the precursor pool minus first-order degradation:

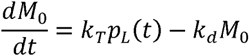

Where *k_t_*is the M_0_ peptide synthesis rate, *k_d_* is the degradation rate, and *p_L(t)_* is the empirically derived estimation for the fraction of newly synthesized peptide that remains fully unlabeled at time *t* (Supplementary Methods). Jointly fitting the control and ^18^O labeling time courses determines k_d_ and k_T_ for each protein, whether it is stable, increasing, decreasing, or any possible combination of these behaviors. Using this approach, we fit a protein half-life of 1.2 hours for GMNN and no measurable degradation for GAPDH (Figure 1f).

Degrading proteins were identified by a Bayesian Information Criterion (BIC) against a null model (ΔBIC > 2; Figure S3a). Degradation rates were highly reproducible between replicates for degrading proteins (R^2^ = 0.82), and all proteins with a fitted half-life longer than three times the ∼71-hour labeling window (∼212 hours) were collapsed to the precision threshold (Figure 1i). Across replicates, 29% of the proteome showed significant degradation, just under half of these (14%) turning over within the experimental window itself, broadly consistent with the largely stable Xenopus proteome inferred indirectly from earlier protein abundance studies^11^.

### Rapid turnover of regulatory proteins against a globally preserved frog proteome

With the kinetic-fitting pipeline established, we examined the structure of turnover across the Xenopus proteome. Fitted degradation rates spanned three orders of magnitude, but the proteome sat overwhelmingly at the slow end, with a long tail of rapidly degrading proteins (Figure 2a). Even within the degrading subpopulation, the median half-life was 83 hours (∼3.5 days), beyond our 71-hour labeling window, and far longer than the median half-lives reported for dividing or non-dividing mammalian cells^28,29^ (Figure 2b). Most of the early Xenopus proteins therefore carry no measurable degradation across development, and even the proteins that do degrade turn over slowly.

**Figure 2.**
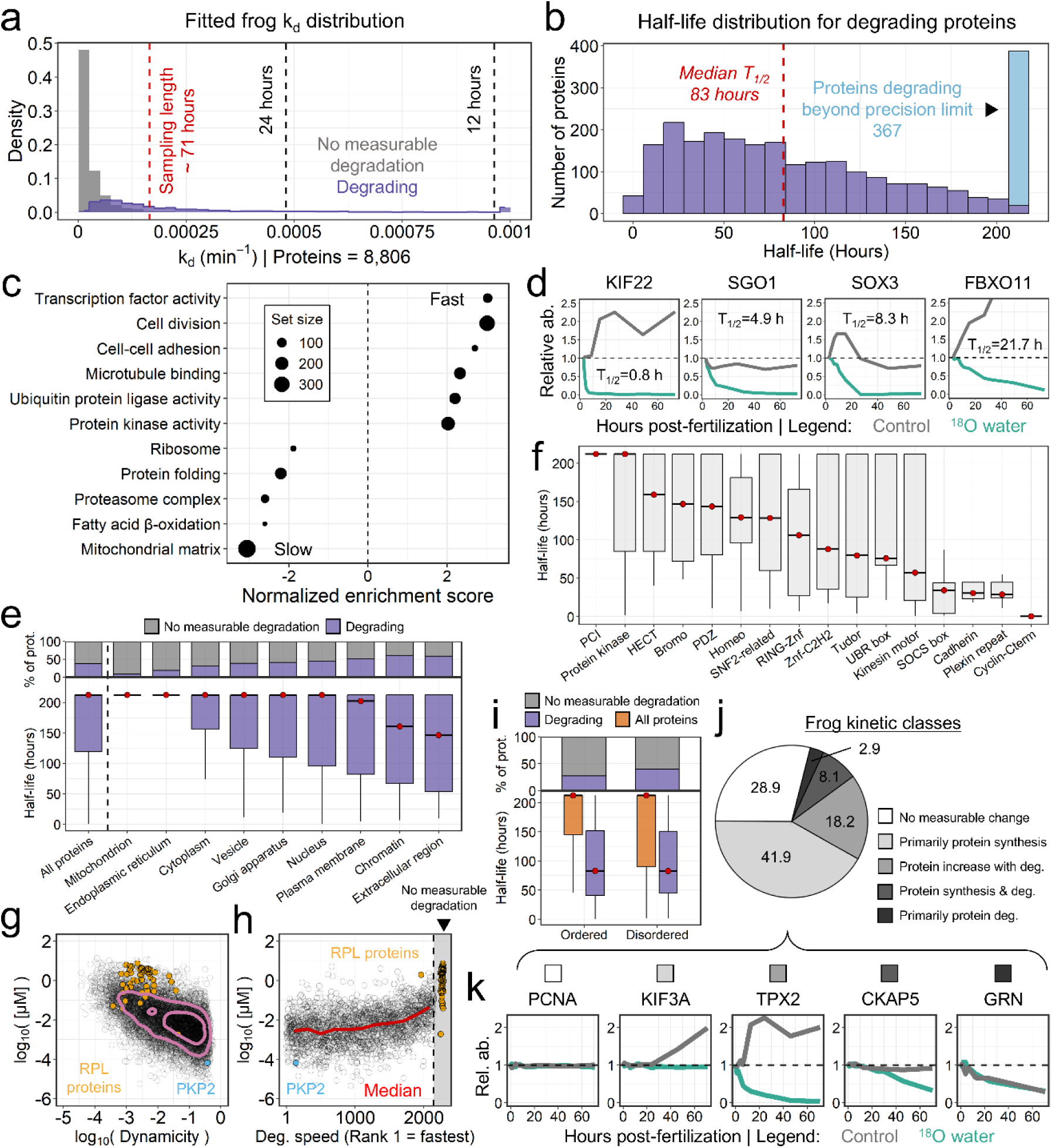
Rapid turnover of regulatory proteins against a globally preserved proteome in the frog embryo. **(a)** Distribution of fitted degradation rate constants (k_d_) for the Xenopus 2-cell time course. Degradation rates were obtained by fitting a single mass-balance model jointly to the ^18^O and control time courses, determining synthesis (k_T_) and degradation (k_d_) for each protein (Supplementary Methods). Proteins exceeding a ΔBIC of 2 against a null model were classified as degrading (purple). Reference lines mark the labeling window (∼71 hours, red) and approximate half-lives of 24 and 12 hours. **(b)** Distribution of Xenopus protein half-lives. For the subset of the proteome undergoing quantifiable degradation, turnover is generally slow, with a median half-life of 83 hours (∼3.5 days). Proteins with half-lives beyond the precision limit (limit = 212 hours, 367 proteins total) are collected in the final bin (blue). **(c)** Functional enrichment of proteins with degradation rates. Positive scores indicate enrichment among fast-degrading proteins. All displayed terms are statistically significant (*p. adj* < 4×10^-3^). Fast-degrading proteins are highly enriched for regulatory functions, including cell division and transcription factor activity, whereas stable proteins are enriched for core metabolic enzymes, such as proteins of the mitochondrial matrix. **(d)** Representative protein examples from the fast-degrading proteins in functional enrichment. Control channels (grey) report total protein abundance; ^18^O channels (green) track decay of the pre-existing protein pool. Examples shown are KIF22, SGO1, SOX3, and FBXO11. **(e)** Protein half-lives grouped by cellular compartment. Top, the proportion of proteins in each compartment classified as degrading (purple) or showing no measurable degradation (grey). Bottom, boxplots of half-life for all proteins in each compartment. Red points mark medians. Mitochondrial and endoplasmic reticulum proteins sit almost entirely at the maximum resolvable half-life (212 hours) with minimal degradation. Chromatin and the extracellular region degrade fastest. **(f)** Relationship between protein domains and corresponding protein half-lives. Boxplots of protein half-lives grouped by functional domain with red points marking medians. Proteins containing classic regulatory domains, such as Cyclin and SOCS box, are among the fastest- degrading populations. **(g-h)** Absolute protein abundance versus dynamicity and turnover rank. **(g)** Dynamicity is the cosine distance of a protein’s trajectory from a flat line, with larger values indicating greater deviation from no change. Abundance is inversely correlated with dynamicity. **(h)** Proteins are rank ordered by degradation speed with 1 as the fastest. Red line marks the running median. Abundance relates weakly to turnover speed, mainly through the slow turnover of the most abundant proteins. Ribosomal proteins (RPL, orange) and PKP2 (light blue) are highlighted in both panels. Non-degrading proteins, including RPL, have no turnover rank and are plotted past the break at the far right. **(i)** Structural disorder and degradation. Top, proportion of ordered and disordered proteins classified as degrading (purple) or showing no measurable degradation (grey). Disordered proteins are more often degrading (40% vs. 27%; χ^2^ test, *p* < 2.2×10^-16^). Bottom, half-life boxplots for all proteins (dark orange) and the degrading subset (purple) with red points marking medians. Among degrading proteins, disorder does not shift the rate (Welch *t*-test on log10(k_d_), *p* = 0.91). Therefore, in Xenopus, protein disorder predicts whether a protein degrades, not how fast it is. **(j)** Global turnover composition. Each protein is assigned to one of five kinetic classes by combining its degradation status from the ^18^O channel (degrading or no measurable degradation) with its abundance trajectory from the control channel (increasing, unchanged, or decreasing). Fold-change cutoffs for increasing and decreasing in the control channel were set by the 1st and 99th percentiles of the non-degrading protein group (Figure S1). The resulting classes show that most of the proteome exhibits no measurable degradation or is primarily accumulating. **(k)** Example trajectories of the five major classes defined in **(j)**. SOX3, a maternally supplied transcription factor that regulates ectodermal and neural gene expression^56,57^, turned over more slowly still, with an estimated half-life of 8.3 hours. The inherited protein is functionally engaged throughout this period, pre-binding ectodermal cis-regulatory modules from cleavage stages onward, before zygotic transcription has begun^57^. At the measured rate, the maternal pool falls to roughly 15% of its initial level by the onset of gastrulation, approximately when zygotic *sox3* transcription begins and its expression becomes spatially restricted to the dorsal ectoderm^58,59^. SOX3 therefore illustrates how protein half-life determines whether transcript measurements report on protein. Because its inherited pool is largely exhausted within the window, sustaining the protein requires zygotic transcription, and transcript and protein dynamics are coupled.

### Although turnover was globally slow, the degrading proteins showed clear functional structure

Gene Ontology (GO) analysis ranked by degradation rate placed cell division, transcription factor activity, and ubiquitin ligases among the fastest, while core metabolic enzymes such as those of the mitochondrial matrix were among the slowest (Figure 2c). This tiering fits their roles: transient regulators turn over rapidly, whereas the metabolic and structural machinery on which the embryo depends is preserved.

Representative trajectories from the faster degrading proteins are shown in Figure 2d. KIF22 (often called Xkid in Xenopus) was one of the most rapidly depleted proteins in our dataset, with an estimated half-life of 49 minutes, approximately one cleavage cycle at 16°C^48^. This is consistent with work showing that degradation of this chromokinesin, which aligns chromosomes at metaphase, is required for anaphase chromosome movement at each division^49,50^. SGO1 was also lost from the inherited pool, but far more slowly than KIF22, with an estimated half-life of 4.9 hours. SGO1 is therefore not quantitatively destroyed in each division and only a small fraction of the inherited pool is lost per cycle. Both proteins are APC/C substrates in Xenopus extracts^49,51^, but the requirement for their destruction differs. KIF22 must be degraded for anaphase chromosome movement to proceed^49,50^, whereas non-degradable SGO1 does not perturb sister-chromatid separation or mitotic exit^52,53^. As a centromeric cohesion protector bound to cohesin^54,55^, only the chromatin-associated SGO1 pool would be exposed to cell-cycle-coupled proteolysis, while most of the maternal pool persists.

Together, these examples illustrate that rapid turnover is not confined to the expected cell-cycle regulators but extends to developmental transcription factors and to components of the degradation machinery itself (Figure 2d). They also span a 27-fold range of half-lives, and in several cases the measured rate corresponds to the timescale over which the protein acts, from clearance within a single division for KIF22 to persistence until zygotic transcription assumes control for SOX3.

Turnover was also organized by location and by domain. Across subcellular compartments, chromatin and extracellular proteins degraded faster than the proteome-wide baseline, while mitochondrial and endoplasmic-reticulum proteins were more stable (Figure 2e). Within annotated domains, classic regulatory domains such as the Cyclin and SOCS-box domains ranked among the fastest, while the PCI domain shared by the proteasome lid, the COP9 signalosome, and eIF3 sat among the slowest (Figure 2f).

Cell-cell adhesion emerged as a recurrent feature of the degrading proteome, enriched among faster-degrading proteins in the GO analysis and among cadherin-associated domain annotations. These proteins had a median half-life of approximately 31 hours and spanned three structurally distinct adhesion systems: the *pcdh1.L/S* homeologs, the classical cadherin *cdh3.L/S*, and the desmosomal cadherins *dsc3.L/S* and *dsg2.S*. Typically, they are more often treated as stable elements of tissue architecture rather than as turnover substrates. However, across the same window in which their total abundance is stable or rising, their inherited pools decay, indicating that maternal adhesion machinery is progressively replaced by newly synthesized protein. The best-supported case is *cdh3.L/S*, the early Xenopus C-cadherin and the major classical cadherin present from egg through blastula stages, whose maternally inherited pool is required for adhesion at the blastula stage^60^. The *pcdh1* homeologs encode axial protocadherin, which mediates prenotochord cell sorting and is expressed in the notochord from the end of gastrulation^61^, so the inherited pool is largely depleted before the protein’s characterized site of action. Desmosomal cadherins are represented as well^62^, though on fewer peptides. The adhesion signal therefore appears at both the process and domain levels and identifies cell-surface adhesion machinery as among the most actively replaced components of the inherited proteome during the window in which embryonic cells remodel their contacts and undergo morphogenetic movements^63^.

*Xenopus laevis* is an allotetraploid, with most genes retained as paired homeologs, short (S) and long (L), derived from two progenitor genomes that merged roughly 17 million years ago^64^. Degradation rates of paired homeologs were typically very similar, and markedly more so than for randomly paired proteins (Figure S4). Therefore, turnover is largely determined by features intrinsic to a protein’s sequence and structure rather than by regulatory differences that have arisen between the copies, which also justifies treating either homeolog as representative in the cross-species comparisons that follow.

Absolute abundance predicted both how much a protein’s level changed and how quickly it was degraded. Abundance was inversely related to dynamicity (Figure 2g), reproducing an effect described previously in Xenopus^11^ where the most dynamic proteins are typically the least abundant. Ribosomal proteins illustrate the point, being both highly abundant and among the least dynamic. Degradation rate tracked abundance in the same direction, though weaker (Figure 2h). Ribosomal proteins again anchor one extreme, being abundant, undynamic, and not measurably degrading, while PKP2, a desmosomal plaque protein that scaffolds junction assembly^65^, anchors the other, being low in abundance, highly dynamic, and rapidly turned over.

Structural disorder acted through a different axis. Disordered proteins were more likely to be degraded than well-folded ones, but among the proteins that did degrade, disorder had no effect on the rate (Figure 2i). In cultured cells and bacteria, disordered segments accelerate turnover^32,66,67^. Disorder could in principle act on either axis, biasing which proteins enter the degrading pool or setting how fast they turn over once they do. In a largely preserved proteome, only the first, propensity, appears free to vary, and disorder in Xenopus therefore predicts whether a protein is degraded, not how quickly.

Because the ^18^O experiment integrates degradation across the entire developmental window, some of the measured turnover could reflect proteins made and destroyed each cell cycle rather than developmental remodeling. To test this, we used standard multiplexed proteomics to profile a single synchronized cell cycle in electrically activated eggs^23^, sampling every 9 minutes over two hours (Figure S5; Table S3). Of 8,052 proteins, only 47 (∼0.6%) were dynamic at all, and just 6 favored a cell-cycle-coupled model. All six oscillating proteins were known cell-cycle regulators^68–70^, and the 34 proteins that decreased were factors shed or secreted at activation^23,71–73^. Cell-cycle-coupled turnover is therefore a negligible fraction of the proteome, and the bulk of the ^18^O signal reflects developmental turnover rather than repeated oscillation.

Finally, we placed every protein into one of five kinetic classes by combining its degradation status from the labeled channels with its abundance trajectory from the control channels (Figure 2j). Most of the proteome falls into two classes that carry no measurable degradation. 29% show no measurable change in either channel (Figure 2k, e.g. PCNA), so any spatial differences that emerge between cells most likely reflect asymmetric partitioning at cleavage rather than synthesis or turnover. A further 42% rise in abundance through synthesis while showing no measurable degradation (e.g. KIF3A). These two classes account for 71% of the proteome and suggest its broad preservation through these stages. The remaining 29% degrade measurably, though net loss is rarer. 18% degrade even as their total abundance rises, so synthesis outpaces turnover (e.g. TPX2), and 8% hold steady through balanced synthesis and degradation (e.g. CKAP5). Only the final 3% decline in both channels, the sole class in true net loss (e.g. GRN). The rising and balanced classes are detectable at all only because the labeling channel resolves the pre-existing pool directly, and with standard abundance-based measurements, they would look like pure synthesis or stability.

### Proteome turnover is largely unchanged from early cleavage to gastrulation

Our measurements so far describe turnover where metabolic labeling started at the 2-cell stage. A natural question is whether protein half-lives are constant or whether they change when measured at a different developmental stage, as the cell cycle lengthens, zygotic transcription begins, and morphogenesis starts. We therefore performed a second ^18^O time course beginning at the onset of gastrulation, roughly 22 hours later (∼24.5 hours post-fertilization at 16°C; Figure 3a). Because this start follows the MZT, it also reports whether the onset of zygotic expression reshapes turnover. As before, we measured the free amino acid pool directly (Figure S6) and used it to compute peptide light-synthesis probabilities. These halved about twice as fast at gastrulation than at the 2-cell stage (median 9 versus 18 minutes; Figure S7a), consistent with the higher metabolic activity of the older embryo, and we accounted for this faster precursor labeling in the kinetic fitting.

**Figure 3.**
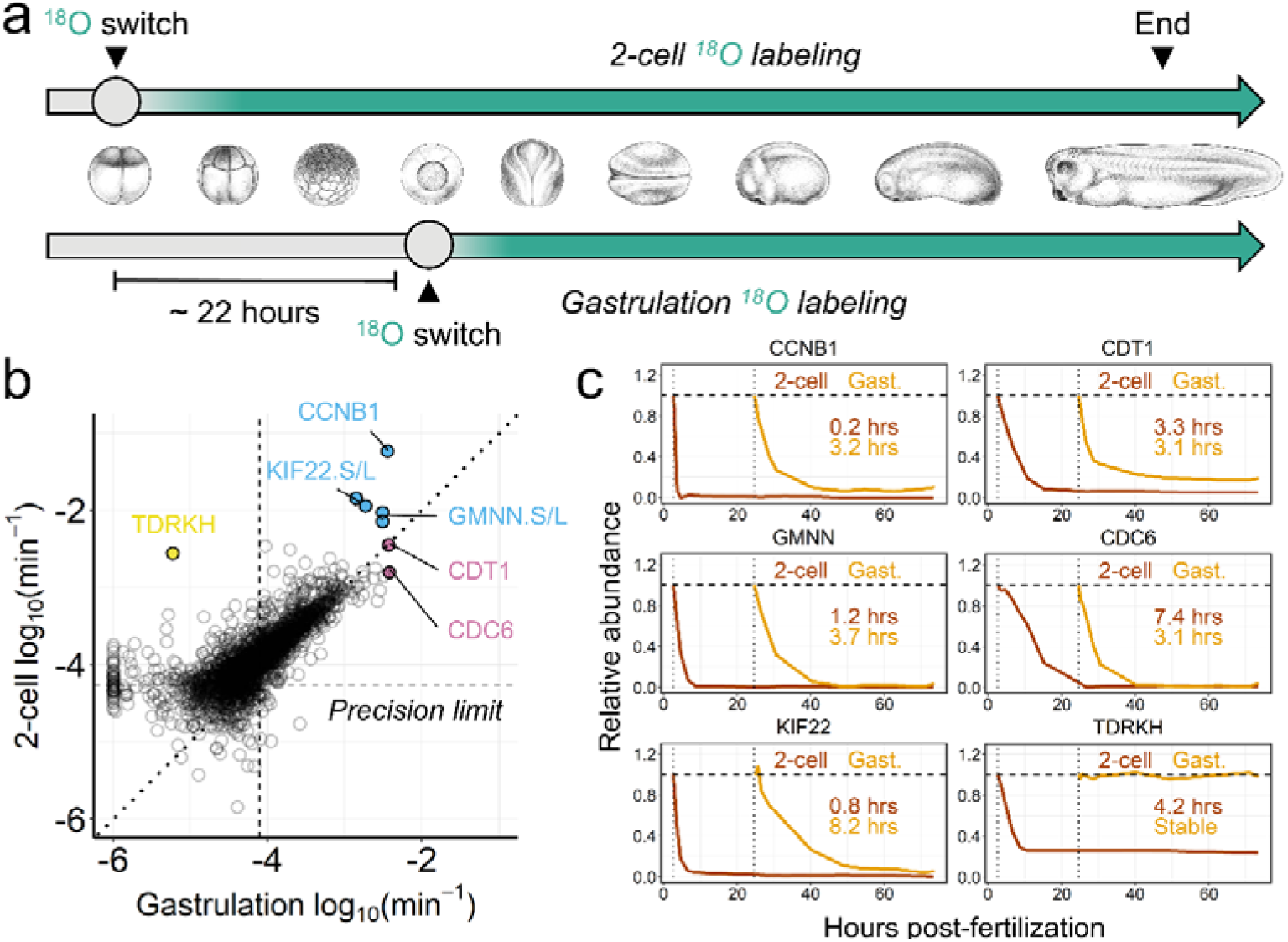
Global proteome turnover is largely conserved from early cleavage to gastrulation in the frog embryo. **(a)** Experimental design comparing early cleavage and gastrulation turnover. Embryos were labeled either from the 2-cell stage (top) or from the onset of gastrulation (∼24.5 hours post-fertilization at 16°C, bottom), offset by ∼22 hours, both ending at hatching. **(b)** Comparison of early and late turnover rates. A scatter plot comparing degradation rates between the 2-cell and gastrulation experiments. While most of the quantifiable proteins maintain equivalent turnover rates (centering along the 1:1 diagonal), specific regulatory proteins deviate. Canonical cell cycle oscillators (e.g., CCNB1, GMNN, KIF22) shift toward slower degradation, whereas CDC6 exhibits accelerated turnover. TDRKH represents a rare outlier that completely stabilizes by gastrulation. **(c)** Representative kinetic shifts. Experimental trajectories comparing 2-cell (brown) and gastrulation (orange) turnover. The apparent slowing of CCNB1, GMNN, and KIF22 turnover directly reflects the known lengthening of the cell cycle at gastrulation. In contrast, CDC6 turnover accelerates, reflecting the onset of strict DNA replication licensing control. Interestingly, it accelerates to match CDT1 (shown as reference, roughly unchanged across stages), its known binding partner. TDRKH stops degrading by gastrulation, after turning over during early cleavage.

We then fit degradation rates as before (Table S4). Within their resolvable ranges, the two experiments were equally reproducible (R^2^ = 0.82 at gastrulation, matching the 2-cell stage; Figure S7b, S7c). Comparing half-lives between the two experiments, most of the quantifiable proteome held very similar rates and fell along the 1:1 diagonal (Figure 3b). The bulk proteome therefore neither systematically accelerates nor slows down between cleavage and gastrulation, and the regime of broad preservation established at the 2-cell stage carries forward.

The kinetic classification appears to show a fraction of the proteome shifting toward stability at gastrulation, but this is an artifact of experimental design, not biology (Figure S7d). Both experiments end at the same developmental point, so the shorter gastrulation window resolves a shorter maximum half-life (147 hours at gastrulation against 212 hours at the 2-cell stage, each three times the respective labeling window). Slowly degrading proteins that were resolvable at the 2-cell stage therefore fall beyond precision at gastrulation and read as stable when they have only dropped below resolution. Mapping each protein’s 2-cell half-life against its gastrulation class reinforces this interpretation. Proteins still quantifiably degrading came almost entirely from the shortest initial half-lives (median 47 hours at the 2-cell stage), while those reclassified as "beyond precision" or "no measurable degradation" had a median initial half-life of 125 hours, closer to the 147-hour ceiling of the gastrulation window (Figure S7e).

Against this stable background, a small set of regulatory proteins shifted in defined and interpretable ways (Figure 3c). The first were canonical cell-cycle proteins that had longer half-lives: CCNB1 from 0.2 to 3.2 hours, GMNN from 1.2 to 3.7 hours, and KIF22 from 0.8 to 8.2 hours. This slowing is consistent with the known lengthening of the cell cycle as embryonic divisions become less rapid, reducing the apparent per-unit-time turnover of oscillatory cell-cycle proteins^74–77^. CDC6 shifted in the opposite direction, accelerating from 7.4 to 3.1 hours and converging on CDT1, which was unchanged at ∼3.2 hours. This behavior is consistent with tightening of DNA replication licensing control after the MZT, when licensing is integrated with longer and more regulated cell cycles^78–81^. A distinct outlier, TDRKH, a piRNA-pathway-associated factor^82^, degraded with a half-life of 4.2 hours at the 2-cell stage but showed no detectable degradation at gastrulation. Because its initial half-life was far shorter than the gastrulation resolution limit, this stabilization cannot be a consequence of the shorter analysis window. That the turnover regime largely persists across the MZT itself is informative, suggesting that the onset of zygotic transcription reshapes the transcriptome without measurably reshaping protein turnover.

### Protein degradation is proteome-wide in the early fly embryo

The Xenopus embryo preserves the bulk of its maternal proteome, degrading only a small regulatory module. To ask whether this preservation is a general feature of embryogenesis or specific to Xenopus, we applied the same ^18^O-water labeling pipeline to Drosophila. Embryos were dechorionated and transferred to an ^18^O-soaked paper towel at the onset of gastrulation (∼3.1 hours post-fertilization at 22°C), and parallel control and labeling series were collected through 24.1 hours post-fertilization, shortly before hatching (∼25.2 hours) (Figure 4a). At the start of labeling the proteome is still overwhelmingly maternal, so the pre-existing pool we track is the maternal proteome present at the onset of gastrulation. As in Xenopus, we measured the free amino acid pool directly (Figure S8) and used it to compute peptide light-synthesis probabilities (Figure S9). Normalization and kinetic fitting then followed the Xenopus pipeline unchanged (Figure S10; Figure S3e), so differences between the species reflect biology rather than differences in analysis. Notably, each species was raised at its standard temperature, 16°C for Xenopus and 22°C for Drosophila.

**Figure 4.**
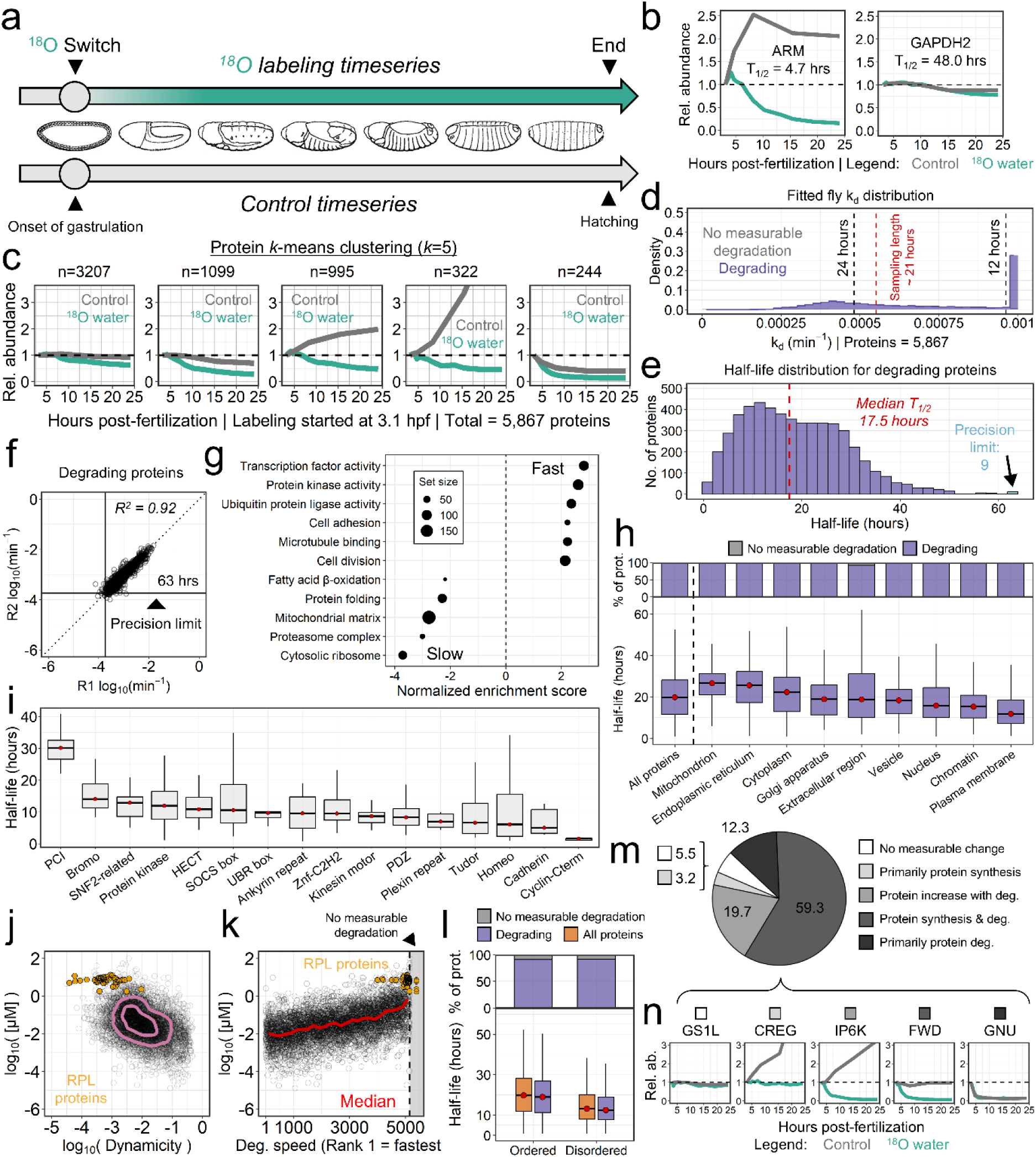
Rapid, global protein degradation dominates the early fly embryo. **(a)** Experimental design. *Drosophila melanogaster* embryos were dechorionated and transferred to an ^18^O-soaked paper towel at the onset of gastrulation (∼3.1 hours post-fertilization at 22°C) to initiate labeling^83^. Parallel control and ^18^O-labeling series were collected through 24.1 hours post-fertilization, shortly before hatching. All samples were combined for multiplexed quantification. **(b)** Representative protein trajectories. Control channels (grey) report total protein abundance; ^18^O channels (green) report decay of the pre-existing pool. Unlike the largely stable Xenopus baseline, both a rapidly degraded regulatory protein (ARM) and a canonical metabolic enzyme (GAPDH2) lose their pre-existing pool. **(c)** Global *k*-means clustering (*k* = 5) of the 5,867 quantified proteins. Decay of the pre-existing pool (green) is present across all clusters, including those whose total abundance (grey) stays flat. **(d)** Distribution of fitted degradation rate constants (k_d_), shifted markedly toward faster degradation relative to Xenopus. Proteins are classified as degrading (purple) or showing no measurable degradation (grey). **(e)** Global half-life distribution. The degrading Drosophila proteome exhibits a remarkably rapid median half-life of 17.5 hours. Proteins with half-lives beyond the precision limit are collected in the final bin. **(f)** Inter-replicate agreement of fitted degradation rates. Agreement is high (R^2^ = 0.92). The dashed line marks the precision limit of 63 hours, three times the ∼21-hour labeling window. Most of the proteome degrades fast enough to be resolved within it. **(g)** Functional enrichment of degradation rate. Positive scores indicate enrichment among fast-degrading proteins. Fast-degrading proteins are enriched for regulatory functions such as cell division and transcription, while slow-degrading proteins are enriched for core metabolic and structural complexes, including ribosomes and proteasomes. The same functional hierarchy observed in Xenopus. **(h)** Protein half-lives grouped by cellular compartment. Top, proportion of proteins in each compartment classified as degrading (purple) or showing no measurable degradation (grey). Bottom, half-life boxplots with red points marking medians. Nuclear and chromatin proteins turn over fastest and mitochondrial proteins slowest, but unlike Xenopus, degradation is present in every compartment. **(i)** Half-lives grouped by protein domain, with red points marking medians. Cyclin and cadherin domains are among the fastest, and even the most stable Xenopus domains, such as the PCI domain, turn over in Drosophila. **(j)** Absolute protein abundance versus dynamicity. Dynamicity is the cosine distance of a protein’s trajectory from a flat line. The inverse relationship between abundance and dynamicity seen in Xenopus is weaker in Drosophila, consistent with a proteome in which nearly everything is dynamic. **(k)** Absolute abundance versus degradation-speed rank (rank 1 = fastest), with the running median in red. Turnover scales inversely with abundance. Ribosomal proteins (RPL, orange) are highly abundant and slow-degrading, falling toward the right-hand end of the ranked axis. Only four RPL proteins have no measurable degradation and are plotted past the break at the far right. **(l)** Structural disorder and degradation. Top, proportion of ordered and disordered proteins classified as degrading (purple) or showing no measurable degradation (grey). Because nearly the whole proteome degrades, disorder does not distinguish whether a protein is degraded (χ^2^test, *p* = 0.50). Bottom, half-life boxplots with red points marking medians. Among degrading proteins, disordered proteins turn over faster than ordered ones (Welch *t*-test on log10(kd), *p* < 2.2×10^-16^). **(m)** Global turnover composition. Each protein is assigned to one of five kinetic classes by combining degradation status (^18^O channel) with abundance trajectory (control channel). ∼91% of the quantified proteome shows measurable degradation. **(n)** Representative trajectories of the five kinetic classes in **(m)**.

The Drosophila proteome behaved in a fundamentally different way from the slow-degrading Xenopus proteome, illustrated by two proteins (Figure 4b). Armadillo, the Drosophila β-catenin homolog, degraded from the inherited pool with a half-life of 4.6 hours while its total abundance rose in the control channels, consistent with its roles in Wingless signaling and adherens-junction assembly^84,85^. This is slower than is often assumed for β-catenin, but the ^18^O rate is a bulk average over the whole embryo, so the more stable junctional pool is folded into the measurement alongside the faster signaling pool.

GAPDH2, a core glycolytic enzyme whose Xenopus counterparts show no measurable degradation at either stage, also degraded, with a half-life of 48.0 hours, among the slowest in this dataset (Table S5). GAPDH degradation during Drosophila embryogenesis has not, to our knowledge, been described, though GAPDH can be a regulated degradation substrate in other contexts^86,87^. RNA-seq shows both are differentially expressed across development^88^, yet only Armadillo increased at the protein level, reinforcing that transcript dynamics are a poor proxy for protein-level behavior. The two span the range from a fast regulatory protein to a slow metabolic one, and Drosophila degrades even the housekeeping enzymes Xenopus preserves.

This species difference was not confined to a few case examples. A *k*-means clustering of the ∼5,900 quantified Drosophila proteins showed decay of the pre-existing pool across every cluster, including clusters whose total abundance stayed flat (Figure 4c), and the distribution of degradation rates was shifted markedly toward faster degradation than in Xenopus (Figure 4d). Among the degrading proteins, the Drosophila median was 17.5 hours (Figure 4e), against ∼83 hours (3.5 days) in Xenopus. Because nearly all Drosophila proteins degrade, the median half-life across all proteins is 18.3 hours, whereas Xenopus has no comparable all-protein median, since its median protein shows no measurable degradation. The acceleration is not an artifact of poorer quantification. The Drosophila measurements were highly reproducible (R^2^ = 0.92), with a precision limit of 63 hours set by the shorter labeling window (Figure 4f), and despite that shorter limit, almost all Drosophila proteins degraded fast enough to be resolved within it, the reverse of Xenopus.

Despite the acceleration, the architecture of turnover was conserved. GO analysis placed fast-degrading Drosophila proteins among regulatory functions such as cell division and transcription, and slow-degrading proteins among core metabolic and structural complexes, including ribosomes and proteasomes (Figure 4g), the same hierarchy seen in Xenopus. Even the large-subunit ribosomal proteins, preserved in Xenopus, degraded in Drosophila with a median half-life of 44.9 hours.

Compartment organization matched as well, with nuclear and chromatin proteins fastest and mitochondrial proteins slowest (Figure 4h), though in Drosophila, unlike Xenopus, degradation was present in every compartment. The extracellular region was an exception, turning over quickly in Xenopus but slower in Drosophila. Because extracellular annotations include species-specific egg coats and embryo-associated matrices that dechorionation and dejellying may affect differently, we do not treat it as directly comparable, and we restricted cross-species comparisons to intracellular classes. Ranking by domain reproduced Xenopus patterns, with cyclin and cadherin domains among the fastest (Figure 4i), but even the most stable Xenopus domains turned over. The PCI domain of the proteasome and COP9 signalosome, which showed no measurable degradation in Xenopus, turned over in 30.2 hours in Drosophila.

The Xenopus protein-level relationships largely carried over, though the inverse relationship between abundance and dynamicity was somewhat weaker in Drosophila (Figure 4j). Turnover again scaled inversely with abundance, with high-abundance proteins degrading slowest (Figure 4k), more strongly than in Xenopus.

Structural disorder revealed the complementary half of the Xenopus result. Because nearly all Drosophila proteins degrade, disorder no longer distinguished whether a protein was degraded (Figure 4l, top). Instead, the effect emerged in the half-lives, where disordered proteins turned over significantly faster than ordered ones (Figure 4l, bottom). Where the preserved Xenopus proteome leaves only propensity free to vary, Drosophila’s near-complete degradation saturates that axis and exposes the rate effect beneath it. Disorder therefore biases proteins toward degradation in both embryos, acting on whichever axis, propensity or rate, is free to move^32,66,67^.

The scale of the difference is clearest in the global composition (Figure 4m, 4n). Of the quantified Drosophila proteome, ∼91% underwent measurable degradation, against 29% in Xenopus. This is likely a conservative floor. Many proteins classified as showing no measurable degradation have trajectories consistent with poor data quality or slow decay rather than true stability (Figure 4n), reflecting the greater experimental difficulty in Drosophila, including imprecise staging and ^18^O-label penetration (Figure S8). Notably, 79% of the quantified Drosophila proteome degrades while its total abundance holds steady or rises, so this turnover is invisible to abundance measurements, and only 12% shows the net decline that abundance alone would detect; the corresponding invisible fraction in Xenopus is 26%.

The Drosophila embryo therefore replaces essentially its entire inherited proteome to varying extents. Yet the organization of that turnover is shared. In both species, regulatory proteins degrade fastest and ribosomal and metabolic proteins slowest, turnover scales inversely with abundance, and disorder biases proteins toward degradation. The two embryos differ not in which proteins are preferentially degraded, but in how much of the proteome degrades at all.

### Frog and fly globally rescale a conserved turnover program to divergent metabolic ends

The Xenopus and Drosophila embryos occupy different turnover regimes. To ask whether this difference reflects a coordinated, proteome-wide rescaling or divergence concentrated in specific pathways, we compared turnover between the species at single-copy orthologs (Figure 5a). The Drosophila experiment begins at gastrulation, but Xenopus turnover is equivalent across the 2-cell and gastrulation stages (Figure 3), so we anchored the comparison on the 2-cell measurements, which have higher measurement precision.

**Figure 5.**
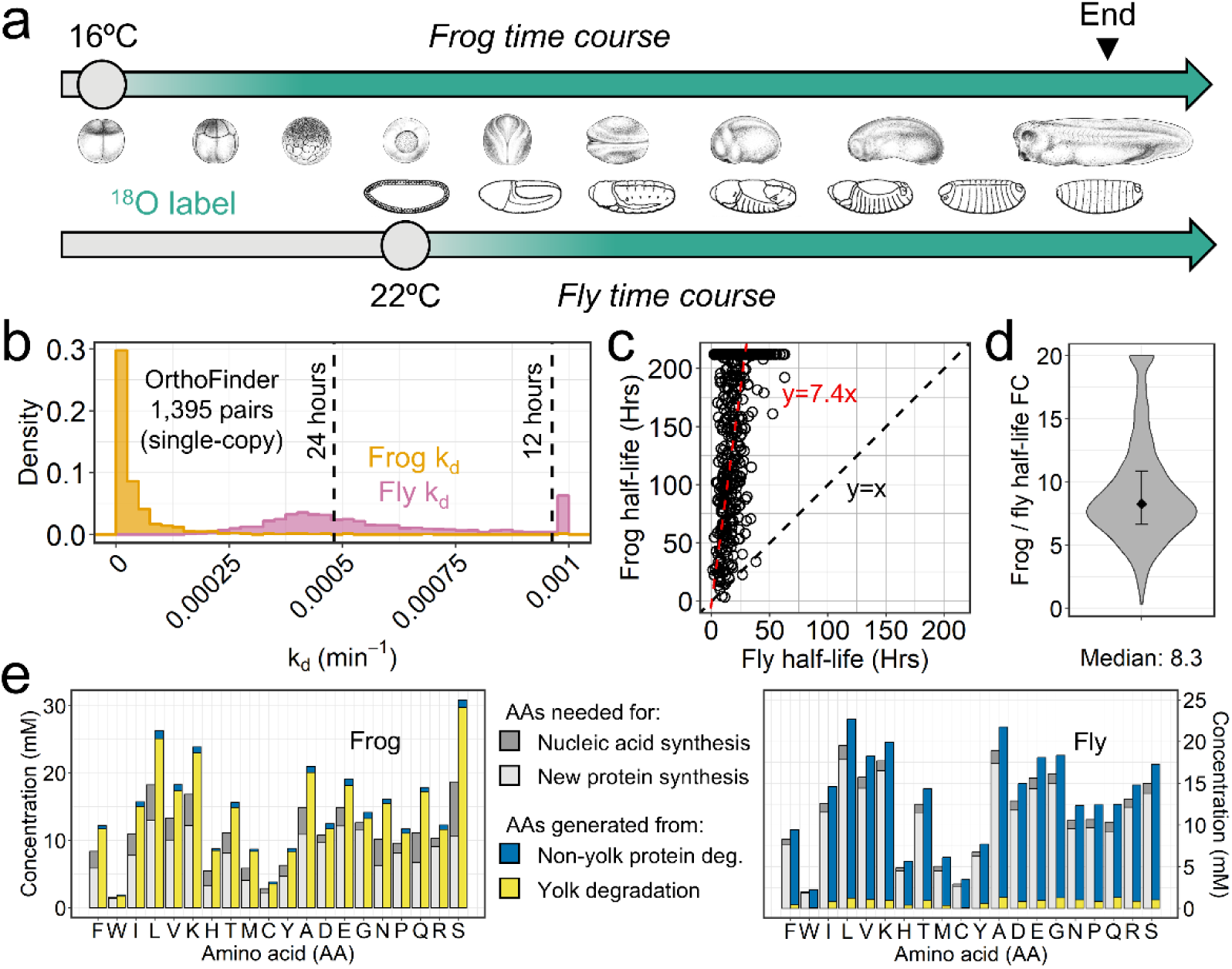
Developmental proteome turnover is globally rescaled between fly and frog. **(a)** Experimental timelines showing the ^18^O labeling windows for Xenopus (16°C) and Drosophila (22°C). Xenopus 2-cell turnover rates were used for the cross-species comparison, as rates at this stage are essentially equivalent to those at gastrulation (Figure 3) while the longer window affords more precise half-life estimates. **(b)** Density distributions of degradation rate constants (k_d_) for the 1,395 single-copy ortholog pairs. Drosophila rates are shifted well above the near-zero Xenopus peak. **(c)** Half-lives of single-copy ortholog pairs plotted against one another. The relationship appears approximately linear across the measured range, with Xenopus half-life ≈ 7.4 × Drosophila half-life (red), well above the line of identity (black dashed). Xenopus proteins whose half-lives exceed the 212-hour precision limit are plotted at that limit and are included in the fit. **(d)** Distribution of per-ortholog fold changes, with a median of 8.3, providing an estimate of the same rescaling independent of the regression in (c). **(e)** Mass-balance models of the free amino acid pool, shown per amino acid for Xenopus (left) and Drosophila (right). Bars show demand, partitioned into new protein synthesis and nucleic acid synthesis, beside supply, partitioned into yolk degradation and degradation of the non-yolk proteome. In Xenopus, yolk supplies nearly all amino acids, and the inherited proteome contributes little. In Drosophila, the proportions are inverted. Yolk contributes a small fraction, and the bulk of the supply comes from degradation of the maternal proteome. In both species supply and demand are closely matched.

We identified single-copy orthologs with OrthoFinder^89^ and restricted the comparison to those well quantified in both experiments. The two degradation-rate distributions are clearly offset, with Drosophila rates shifted well above the near-zero Xenopus peak (Figure 5b). Comparing half-lives ortholog by ortholog (Figure 5c) gives a median fold-change of 8.3 (Figure 5d). This shift is unimodal. Orthologs are not split into species-specific subsets in specific pathways but are instead displaced together proteome-wide. Xenopus proteins whose half-lives exceed our 212-hour precision limit are censored at that bound, so 8.3-fold is a conservative estimate of the true difference.

This shift exceeds what the difference in developmental tempo alone would predict. At a common temperature, Drosophila development from fertilization to hatching is roughly twice as fast as Xenopus^48^, and under our conditions Xenopus took ∼73 hours against Drosophila’s ∼25 hours. Ortholog half-lives differ by considerably more. Expressed per unit of developmental time, Xenopus proteins therefore persist roughly 2-to 3-fold longer than their Drosophila counterparts. Thus, Drosophila does not merely run the same program on a faster clock, it degrades more of its proteome and does so faster than its shorter development requires. Any uniform normalization shifts all orthologs equally and so changes the magnitude of this difference without changing its direction. Together with the conserved ordering of turnover within each species, this indicates that the same kinetic architecture operates in both embryos, across more than 500 million years of evolution^90^, at globally different rates.

These regimes carry a metabolic consequence, which we examined by building mass-balance models of the free amino acid pool in each species (Figure 5e, Table S6, Table S7). In Xenopus, yolk supplies most amino acids for new synthesis, and degradation of the non-yolk proteome adds only a small further pool, as expected for a proteome that is preserved rather than recycled. In Drosophila, the balance is reversed. Yolk catabolism begins at cellularization and proceeds through the interval we measure^91^, so yolk-derived amino acids are available throughout. However, yolk constitutes only ∼9% of the deposited protein (Table S7), and its complete degradation would not supply what new synthesis requires. Degradation of the maternal proteome must therefore make up the shortfall.

The two embryos also differ in what they can do with the nitrogen that degradation releases. In Drosophila, amino acid release and reincorporation are closely matched. However, the small surplus must still be disposed of, and disposal is limited by where the embryo develops. Enclosed within a chorion in a semi-solid medium, with no surrounding water to carry waste away, the embryo cannot excrete ammonia. Surplus nitrogen is instead fixed into uric acid, which is insoluble and inert^92^. We measured uric acid across Drosophila embryogenesis and found a roughly 8-fold rise, from 0.8 to 6.6 mM per embryo (Figure S11), with the increase confined to the interval of maximal proteome turnover. The nitrogen sequestered this way represents only a few percent of the nitrogen held in the proteome, consistent with degradation that is near-quantitatively recycled into new protein rather than catabolized. The Drosophila energy budget is met accordingly, by oxidation of maternal glycogen and triacylglycerol rather than protein^93^.

Xenopus is subject to neither limit. Developing in pond water, it can excrete surplus nitrogen directly as ammonia, with no need for an insoluble sink. Its yolk reserve is correspondingly larger and is not exhausted until well after hatching^94^, so within the window we measure, yolk supplies more amino acids than synthesis consumes, and the excess is available for oxidation^36^.

A complementary accounting follows total protein mass across development (Figure 6). Yolk constitutes ∼86% of the protein deposited in the Xenopus egg^94^ but only ∼9% in Drosophila (Table S7). Xenopus therefore draws on a reserve that dwarfs its synthetic demand and leaves its inherited proteome intact, whereas Drosophila, lacking that reserve, consumes the proteome itself (Fig. 6). Widespread maternal degradation in Drosophila is thus not incidental to development but the source of amino acids that synthesis demands and yolk cannot provide.

**Figure 6.**
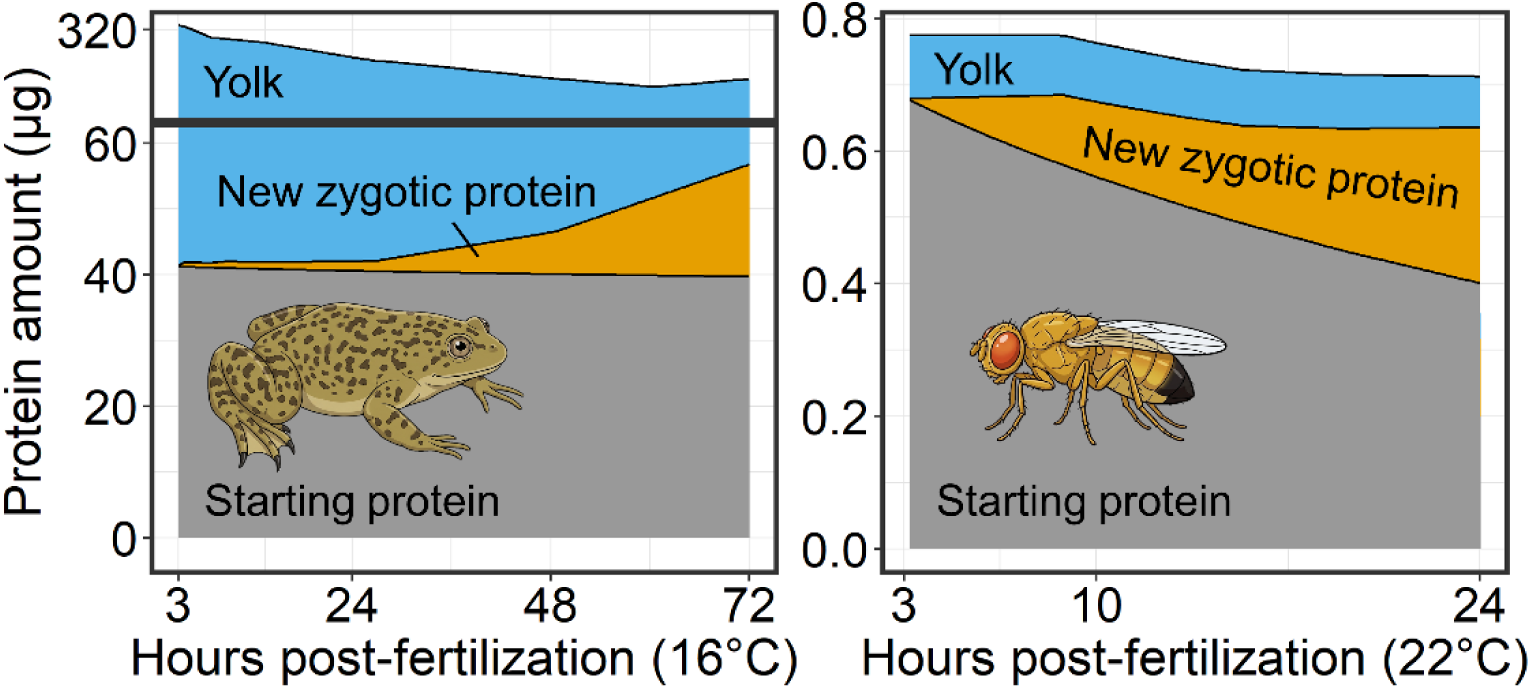
Divergent strategies for building the embryonic proteome in frog and fly. Total protein mass across development, partitioned into yolk (blue), the maternally deposited non-yolk proteome ("starting protein," grey), and newly synthesized zygotic protein (orange). Xenopus (left, 16°C) and Drosophila (right, 22°C); note the axis break in the Xenopus panel. In Xenopus, yolk constitutes ∼86% of the deposited protein mass^94^ and is slowly consumed to supply amino acids for new synthesis, while the non-yolk proteome remains largely intact. In Drosophila, yolk is a much smaller fraction of the starting material (∼9%; Table S7) and its degradation alone cannot supply zygotic synthesis. Instead, the amino acids are drawn from the maternal proteome itself, which is progressively replaced by newly synthesized protein. Roughly two-thirds of Drosophila’s starting protein remains at hatching.

## Discussion

We set out to directly measure the fate of the maternal proteome rather than infer it. ^18^O-water labeling renders the pre-existing protein pool directly visible as a decaying unlabeled signal, allowing proteome-wide turnover to be quantified in intact, non-feeding embryos. Applied to Xenopus and Drosophila, it revealed two embryos that solve the same problem through opposite strategies. Both run development on a finite maternal endowment, yet Xenopus preserves the bulk of its maternal proteome through hatching while Drosophila degrades nearly all its maternal proteins.

These results resolve an apparent contradiction in the literature. Quantitative studies in Drosophila concluded that the maternal proteome is extensively remodeled^21,22^, whereas studies in Xenopus concluded that it is largely stable^11,23^. Because neither could measure turnover directly, each adopted the simplest interpretation consistent with its abundance data, and our direct measurements now show that each was essentially correct. One caveat tempers this reconciliation. The prior Drosophila inferences were drawn largely from the oocyte-to-embryo transition, an earlier window than the gastrulation-to-hatching interval we measured here. Our data therefore extend those observations rather than directly overlay them, but they confirm that pervasive degradation of the maternal proteome is a genuine, species-specific feature of Drosophila development and not of Xenopus.

Direct measurement also sharpens what "stability" means. In Xenopus, nearly a third of all protein species show statistically detectable degradation, yet the median half-life of ∼83 hours (3.5 days) among them exceeds the ∼73 hours from fertilization to hatching, and the most abundant proteins degrade most slowly. By mass, therefore, the maternal proteome is largely preserved even as slow degradation is common across protein species. This coupling between abundance and turnover is not confined to Xenopus. It holds even in Drosophila, where nearly the entire proteome degrades yet the fastest turnover falls on the least abundant proteins and the slowest on the most abundant, so that roughly two-thirds of the maternal protein mass is still present at hatching. This is a distinction that abundance measurements alone cannot draw, and one that explains why earlier work reasonably inferred a stable proteome from the observation that protein levels change little. Degradation rapid enough to replace the maternal pool within development is confined to a small, functionally coherent module of cell-cycle and regulatory proteins.

These half-lives also determine how faithfully mRNA measurements report on the proteome. Because a transcript and its protein are decoupled whenever the protein outlives the developmental window, protein stability sets the fidelity with which transcript dynamics predict protein dynamics. In Xenopus, most maternal proteins persist through hatching, so maternal transcript clearance can proceed with little corresponding loss of protein^11,95^. Thus, mRNA is a poor proxy for the Xenopus proteome. In Drosophila, where nearly every protein turns over on the timescale of development, transcript-level changes likely propagate to protein more directly. The poor mRNA-protein correspondence reported across embryos is therefore not a single phenomenon but a species-specific consequence of how fast each proteome turns over.

Against these divergent strategies, the architecture of turnover is conserved. In both species, the most disordered and regulatory proteins degrade fastest, turnover scales inversely with abundance, and ribosomal and metabolic proteins degrade most slowly. These are the same determinants described in microbes, cell lines, and primary cells^28,29,32,67^. What differs between the two embryos is global rate.

Drosophila proteins degrade roughly eightfold faster than their Xenopus orthologs, and the shift shows no sign of being concentrated in particular pathways. The two embryos thus differ in how fast and in how many of their proteins they turn over, not in which proteins they preferentially degrade. A global, species-specific offset in protein stability has recently been described between mouse and human in stem-cell-derived progenitors, where human proteins are roughly 1.5-fold more stable and the difference tracks developmental tempo, metabolic rate, and proteasome activity^33–35^. Our comparison shows the same principle in intact embryos separated by more than 500 million years, at a far larger magnitude, and ties it to the metabolic role of the maternal proteome.

Beyond its regulatory role, proteome turnover carries metabolic consequences. Externally developing embryos are closed systems that must build themselves from maternally deposited reserves^91,93,96^, drawing down yolk, central metabolites, and nucleotide pools as development proceeds^36,97^. A natural question is then why Drosophila would degrade what Xenopus preserves. A simple accounting of amino-acid supply offers an explanation. The Xenopus egg is dominated by yolk, which more than suffices to supply the amino acids for new synthesis, so the maternal proteome need not be consumed^98^. The Drosophila egg carries proportionally far less yolk, and that yolk alone cannot meet the embryo’s synthesis demand. Even if all the yolk present within our analysis window were fully degraded by hatching, it would fall short of what the embryo’s protein and nucleic acid synthesis require. The shortfall can be made up only by recycling the proteome itself, so Drosophila’s pervasive degradation is not optional but required. That it is recycling rather than catabolism is borne out by the limited nitrogen the embryo sheds as uric acid^92^ and by an energy budget met from maternal glycogen and triacylglycerol rather than protein^93^.

Methodologically, ^18^O-water labeling removes the central obstacle to studying turnover in embryos. To our knowledge, no prior method has measured proteome-wide protein degradation directly in an intact developing metazoan embryo. Existing approaches that rely on labeled diet, injection, or bioorthogonal amino acids are incompatible with non-feeding embryos. Single fluorescent fusion proteins can be followed in zebrafish embryos^99^, and bioorthogonal labeling reports newly synthesized protein in larvae^100^, but neither yields degradation rates across the proteome. Recent advances have made turnover measurement easier and more sensitive in feeding systems, through DIA-based workflows and single-cell pulse-SILAC^101,102^. ^18^O-water labeling is complementary, extending direct turnover measurement to the non-feeding embryos those methods cannot reach. Because water equilibrates into any organism and proteolysis universally transfers oxygen from water to the liberated amino acid, ^18^O-water labeling requires no feeding, injection, or genetic manipulation, and should be applicable to essentially any embryo or other non-feeding system that can be bathed in water. The resulting Xenopus and Drosophila half-life datasets also provide a proteome-wide community reference for which maternal proteins persist, which are replaced, and how quickly.

Finally, our comparison rests on two species, and we are correspondingly cautious about generalization. The conserved determinants of turnover are well precedented and are likely general, but whether the preserve-versus-replace strategies we observe are two recurrent solutions or two samples from a broader range of embryonic strategies cannot be settled with two organisms. Prior proteomic work in other embryos has been limited to protein abundance^11,12,16^, which our turnover measurements now complement, and ^18^O-water labeling makes the necessary comparisons straightforward. Extending it to zebrafish, sea squirt, sea urchin, nematode, and mammalian embryos, which differ in yolk content, would directly test the prediction implicit in our metabolic argument, that the amount of yolk an embryo inherits should determine how much of its maternal proteome it must consume to build itself. More broadly, our results recast the maternal proteome as at once an informational inheritance and a metabolic reserve, and protein turnover as the process that negotiates between the two. Turnover itself has been the least accessible of the quantities that define a proteome, long inferred from snapshots of abundance rather than measured. Water-based labeling now makes it directly quantifiable across organisms and cellular processes, and treating turnover as a systems-level variable in its own right should reveal regulation, in development and beyond, that abundance alone keeps invisible.

## Methods

### *X. laevis* embryo collection and lysis

Mature *X. laevis* females and males were purchased from Xenopus1 and maintained by Laboratory Animal Resources at Princeton University. All animal procedures were approved by the Princeton University Institutional Animal Care and Use Committee (protocol 2070) and were carried out in accordance with institutional and national guidelines. Unfertilized eggs and male testes were collected following standard laboratory procedures previously described^103^. For testes collection, *X. laevis* males were euthanized in 0.1% (w/v) tricaine methanesulfonate (MS-222; Syndel’s Syncaine) and then sacrificed by pithing. The testes were isolated and stored at 4°C in oocyte culture medium that was exchanged daily for up to 1 week. Oocyte culture medium is a Leibovitz’s L-15-based medium containing 13.7 g/L Leibovitz’s L-15 Medium powder (Thermo Fisher Scientific, 41300039), 0.67 mg/mL bovine serum albumin, and 0.83% v/v penicillin-streptomycin (Thermo Fisher Scientific, 15140122). For egg collection, female frogs were injected with 500 U of human chorionic gonadotropin (CG10; Sigma) and kept at 16°C in Marc’s modified Ringer’s solution for 16 hours before collection (1X MMR: 5 mM HEPES [pH 7.8], 0.1 mM EDTA, 100 mM NaCl, 2 mM KCl, 1 mM MgCl2, and 2 mM CaCl2). For *in vitro* fertilization, female eggs collected were cleaned in 1X MMR buffer, and preactivated eggs were removed.

Half of one male testis was used per 500 eggs by crushing in 1X MMR buffer with a sterile pestle and then mixing with the unfertilized eggs. The mixture was incubated at 16°C for 5 minutes, followed by mixing and an additional 5-minute incubation. Fertilization was induced by flooding the eggs with 0.1X MMR. After 1 hour at 16°C, embryo jelly coats were removed by incubating with 2% cysteine in 200 mM KCl, 2 mM MgCl_2_, and 0.2 mM CaCl_2_ [pH 7.8] for 5 minutes, and the embryos were thoroughly washed with 0.1X MMR to remove residual cysteine. Embryos were grown and staged according to Nieuwkoop and Faber nomenclature^104^ at 16°C and then flash-frozen in liquid nitrogen at desired time points.

Embryos were pooled and of undetermined sex, as the stages analyzed precede sexual differentiation; sex was therefore not considered as a variable. For ^18^O-labeling, embryos were transferred following successful first cleavage or at the onset of gastrulation into 0.1X MMR prepared with 97% ^18^O-water (Cambridge Isotope Laboratories).

For the ^18^O experiments, Xenopus embryos were lysed as previously described^105^. Briefly, a lysis buffer consisting of 250 mM sucrose, 1% Nonidet P-40, 10 mM EDTA, 25 mM HEPES (pH 7.2), 10 μM cytochalasin D, and Roche cOmplete protease inhibitor cocktail (1 tablet per 10 mL) was added to the embryos and pipetted up and down 15 times. Samples were then incubated on ice for 10 minutes and vortexed for 10 seconds. Yolk was then removed with a soft spin at 2500 x g for 4 minutes at 4 °C. Next, 100 mM HEPES, pH 7.2, and 2% SDS were then added to denature proteins.

For normalization, Xenopus embryos were prepared without removing the yolk. Samples were lysed in buffer containing 4% SDS and 50 mM HEPES (pH 7.2) and Roche EDTA-free cOmplete protease inhibitor cocktail (1 tablet per 10 mL) and sonicated at 50% amplitude for five cycles of 10 seconds on and 20 seconds off, with samples held on ice between pulses.

### *D. melanogaster* embryo collection and lysis

Canton-S *D. melanogaster* flies were placed in egg-collection chambers fitted with apple-juice agar plates. Plates were collected after approximately 2 hours and transferred to a water-soaked paper towel. Because multiplexed proteomics requires substantial total protein, embryos were collected in bulk rather than individually staged. Bleach was applied to dissolve the chorion, and once the chorion had visibly dissolved, the bleach was removed by blotting the paper towel dry, rinsing with water, and blotting dry again. For ^18^O-labeling, the dechorionated embryos were then transferred to either normal water or 97% ^18^O-water (Cambridge Isotope Laboratories). Eggs accumulated across the approximately 2-hour collection window, so at the start of labeling, embryos spanned roughly 2 to 4 hours post-fertilization. We treated the midpoint, approximately 3 hours post-fertilization, as the labeling start. Embryos were collected in bulk and comprised both sexes; sex was not considered as a variable.

Embryos were sampled at defined times relative to this start and immediately heat-fixed. For heat fixation, embryos were collected onto a 2-cm-diameter metal mesh basket and immersed in a vial containing 5 mL boiling Triton-salt solution (0.4% NaCl, 0.01% Triton X-100). The vial was immediately transferred to an ice bath, and 1 ml of ice-cold water was added. After 5 minutes, the mesh basket was removed with tweezers and the Triton-salt solution was pipetted off, leaving the embryos at the bottom of the vial. Intact embryos were then pipetted out, collected into a microcentrifuge tube, rinsed three times with methanol, and stored at -20°C.

Drosophila samples were lysed in buffer containing 4% SDS and 50 mM HEPES (pH 7.2) and Roche cOmplete protease inhibitor cocktail (1 tablet per 10 mL). Following removal of methanol and air-drying, pellets were resuspended in lysis buffer and sonicated at 50% amplitude for five cycles of 10 seconds on and 20 seconds off, with samples held on ice between pulses. Lysates were then incubated at 70°C for 30 minutes and then clarified by centrifugation at 20,000 x g for 10 minutes.

## Proteomic sample preparation

Proteomic samples were prepared as follows. Following cell lysis, samples were reduced with 5 mM dithiothreitol (DTT) for 20 minutes at 60°C and alkylated with 20 mM N-ethylmaleimide (NEM) for 20 minutes at room temperature. 5 mM DTT was added to quench the excessive alkylating reagents.

Proteins were precipitated following reduction and alkylation using the SP3 method, as previously described^106^. After binding and washing the bead-bound protein, the protein-containing beads were resuspended in 2 M guanidine hydrochloride and 10 mM EPPS (pH 8.5). The resulting bead-protein mixture was digested with 20 ng/μL LysC (Wako) overnight at room temperature. Samples were further diluted 4-fold with 10 mM EPPS (pH 8.5) and digested with an additional 20 ng/μL LysC and 10 ng/μL sequencing-grade trypsin (Promega) at 37°C for 16 hours. After digestion, the supernatant was vacuum-dried, resuspended in 200 mM EPPS (pH 8.0), and clarified at 20,000 x g for 20 minutes to remove residual beads. For TMTpro labeling, TMTpro tags were added at a ratio of 5 μg of TMTpro:1 μg of peptide, mixed, and incubated at room temperature for 2 hours. The reaction was then quenched by addition of 1 μL of 5% hydroxylamine per 10 μL of reaction at room temperature for 30 minutes. The samples were pooled, and the resulting mixture was vacuum-dried. Both Data-Dependent Acquisition (DDA) and Data-Independent Acquisition (DIA) samples were desalted with C18 material (Empore) following digestion clarification and resuspended in 1% formic acid to 1 μg/μL for LC-MS analysis.

Prefractionation was utilized to detect a larger number of peptides in multiplexed samples^107^. Specifically, the dried TMTpro-pooled peptides were resuspended in 10 mM ammonium bicarbonate (pH 8.0) with 5% acetonitrile to a peptide concentration of 1 μg/μL. The dissolved peptides were separated into 96 fractions using medium-pH reverse-phase separation (Zorbax 300Extend C18, 4.6 × 250 mm column) on a 1260 Infinity II LC system (Agilent), as described previously^107^. Each resulting 96-well plate was combined into 24 fractions, and each fraction was desalted and resuspended for LC–MS analysis.

## Xenopus egg activation experiment

Unfertilized Xenopus eggs were collected as described above and activated as previously described^23^. Briefly, eggs were placed in an open-faced gel box on a 3% agar bed in 0.1X MMR and activated by applying an electric field of ∼3 V/cm for 1 second. Eggs were maintained at 16°C and collected every 9 minutes until 124 minutes post-activation. Excess medium was removed and samples were flash-frozen in liquid nitrogen.

Samples were processed as in the standard workflow, with the following addition. To remove phosphorylation dynamics that would otherwise confound total-protein quantification, activated samples were treated with a temperature-labile phosphatase that can be inactivated before multiplexing. For this, we used temperature-labile shrimp alkaline phosphatase (product #78390; Affymetrix). To minimize the volume added to each sample, the enzyme was concentrated to ∼4 U/µL on a 5-kDa Amicon Ultra filter by centrifugation at 4°C. Because the supplied buffer contains Tris, which interferes with TMT labeling, the enzyme was buffer exchanged into 10 mM EPPS (pH 8.0) and stored in 50% glycerol.

After LysC-trypsin digestion, the digests were cooled to room temperature and supplemented with 10 mM MgCl₂, 0.1 mM EDTA, and 5 mM EPPS (pH 8.5). Phosphatase was added to the activated samples at a 3U:1µg ratio of peptide to phosphatase units, directly into the tryptic digest without an intervening peptide cleanup and incubated for 12 hours at room temperature. All samples were then heated at 65 °C for 5 minutes in a water bath to inactivate the phosphatase, dried by vacuum centrifugation, and carried into TMTpro labeling as described above.

## Metabolite extraction and analysis

Flash-frozen Xenopus and Drosophila samples were collected and cryomilled (5 cycles of 20 Hz for 2 minutes with 5 Hz for 30 seconds as cooling intervals; Retsch). 55 µL of cold (-20°C) acetonitrile:methanol:water (2:2:1 v/v/v; all solvents HPLC grade) was added to the samples and immediately vortexed. All remaining steps were carried out at 4°C. After 20 minutes on ice, precipitated macromolecules (protein, DNA, RNA) were pelleted by centrifugation (20,000 x g for 5 minutes), and the supernatant was collected. The pellet was re-extracted with 25 µL of the same solvent mixture by repeating the steps above, and the supernatants were combined to give the final metabolite extract. Metabolites were then measured as previously described^108^.

## UHPLC chromatography for proteomics

All proteomic samples were analyzed on a Vanquish Neo UHPLC system. Solvent A consisted of 2% DMSO and 0.125% formic acid in water, and solvent B consisted of 100% acetonitrile, 2% DMSO, and 0.125% formic acid in water. The UHPLC system was coupled to a modified Orbitrap Ascend mass spectrometer (Tune 4.1). Peptides were separated on an Aurora Series emitter column (25 cm x 75 μm inner diameter, 1.6 μm C18; IonOpticks) and were held at 60°C using an in-house-built column oven.

TMTpro-labeled samples were analyzed with the following 90-minute gradient at a constant flow rate of 350 nL/min after thorough equilibration of the column to 0% B: 0-10% B for 5 minutes, 10-26.4% B for 70 minutes, 26.4-100% B for 10 minutes, and 100% B for 5 minutes.

DDA label-free samples were analyzed with the following 180-minute gradient at a constant flow rate of 350 nL/min after thorough equilibration of the column to 0% B: 0-4.8% B for 5 minutes, 4.8-26.4% B for 160 minutes, 26.4-100% B for 10 minutes, and 100% B for 5 minutes.

DIA label-free samples were analyzed with the following 70-minute gradient at a constant flow rate of 350 nL/min after thorough equilibration of the column to 0% B: 0-4.8% B for 5 minutes, 4.8-26.4% B for 50 minutes, 26.4-100% B for 10 minutes, and 100% B for 5 minutes.

For electrospray ionization, 2.6 kV was applied between 1 and 7 minutes before the end of the LC gradient (TMT: 83 minutes; DDA label-free: 173 minutes; DIA label-free: 63 minutes). To avoid carryover of peptides, 2,2,2-trifluoroethanol was injected in a 30 min wash between each sample^109^. For fractionated samples, this wash was performed between every three fractions from the same original sample.

## RTS-MS3 method

The mass spectrometer was set to analyze positively charged ions in data-dependent MS3 mode, recording centroid data with the RF lens level at 60%. Full scans were taken with the Orbitrap at 120k resolution with an automatic gain control (AGC) target of 4 x 10^5^ charges, maximum ion injection time (IIT) of 50 milliseconds, and scan range of 350-1500 m/z with wide quadrupole isolation enabled. The maximum cycle time between MS1 scans was set to 3 seconds.

Following the survey scan, the following filters were applied for triggering MS2 scans. Monoisotopic peak selection (MIPS) was enabled and set to isolate the monoisotopic peak in the peptide mode. Isolated masses were excluded for 60 seconds after triggering with a mass tolerance window of ±10 ppm while also excluding isotopes and different charge states of the isolated species. A charge-state filter was set for 2-3+ charge states.

The following settings were used for ion-trap MS2 scans. The AGC target was set to 1 x 10^4^ charges, and the maximum IIT was 50 milliseconds. The quadrupole was utilized for isolation with an isolation width of 0.5 Da, and ions were fragmented with collision-induced dissociation (CID) at a normalized collision energy of 35% (10 milliseconds activation time and 0.25 activation Q). The scan rate was 66 kDa/s (rapid resolution) in autoscan range mode.

Exploratory ion trap scans were searched against an appropriate FASTA file with common contaminants as described in DDA database search. Variable methionine oxidation (+15.9949 Da), static TMTpro modifications on lysines/peptide N termini (+304.2071 Da), and static NEM modification of cysteine residues (+125.0477 Da) were allowed. The maximum variable modifications setting was 1, the maximum missed cleavages setting was 1, and the maximum search time was 40 milliseconds. RTS score thresholds were set as follows. Peptide were deemed acceptable if their cross-correlation was greater than 1.4, their delta cross-correlation was greater than or equal to 0.1, and their absolute precursor parts per million deviation was less than or equal to 10.

Following a successful exploratory scan, the following filters were applied. Precursor selection range was set to 400-2000 m/z. Precursor ion exclusion was set to 5 (high) and 70 (low) m/z. Isobaric tag loss exclusion was set to TMTpro, and 5 notches were used to isolate the MS3-precursors with Synchronous Precursor Selection (SPS). MS3 scans were acquired with a 0.4 m/z MS1 isolation window and a 2 m/z MS2 isolation window. The Orbitrap detector was used with a resolution of 45k and an AGC target of 2 x 10^5^, scanning over the range 110-400 m/z. The maximum ion injection time was 91 milliseconds. Ions were fragmented in the higher-energy collision dissociation (HCD) cell at a normalized collision energy of 45%.

## DDA label-free method

The mass spectrometer was set to analyze positively charged ions in data-dependent MS2 mode, recording centroid data with the RF lens level at 60%. Full scans were taken with the Orbitrap at 120k resolution with an automatic gain control (AGC) target of 4 x 10^5^ charges, maximum ion injection time (IIT) of 50 milliseconds, and scan range of 350-1400 m/z with wide quadrupole isolation enabled. The maximum cycle time between MS1 scans was set to 3 seconds.

Following the survey scan, the following filters were applied for triggering MS2 scans. Monoisotopic peak selection (MIPS) was enabled and set to isolate the most abundant peak in the peptide mode. Isolated masses were excluded for 30 seconds after triggering with a mass tolerance window of ±10 ppm while also excluding isotopes and different charge states of the isolated species. A charge-state filter was set for 2-6+ charge states.

The following settings were used for ion-trap MS2 scans. The AGC target was set to 3 x 10^4^ charges, and the maximum IIT was 13 milliseconds. The quadrupole was utilized for isolation with an isolation width of 0.5 Da, and ions were fragmented with HCD at a normalized collision energy of 35%. The scan rate was 125 kDa/s (turbo resolution) with a defined scan range of 200-1400 m/z.

## DIA label-free method

The mass spectrometer was set to analyze positively charged ions in DIA mode, alternating each cycle between a full survey scan and a fixed set of precursor isolation windows. Full scans were taken with the Orbitrap at 120k resolution recording profile data with the RF lens level at 30%, an AGC target of 4 × 10^5^ charges, automatic maximum IIT, and a scan range of 350-1350 m/z with wide quadrupole isolation enabled.

Following each survey scan, DIA MS2 scans were acquired across a precursor range of 350-950 m/z using 75 sequential quadrupole isolation windows of 8 m/z with no window overlap. Precursors were fragmented with HCD at a normalized collision energy of 30%. Fragment spectra were recorded with the Orbitrap at 15k resolution in centroid mode, with an AGC target of 5 × 10^5^ charges and a maximum IIT of 27 milliseconds. Auto scan range mode was used.

## DDA database search

The data were analyzed using GFY software licensed from Harvard University. Raw files were converted to mzXML using ReAdW.exe. MS2 spectra assignment was performed using the SEQUEST algorithm v.28 (rev. 12) by searching the data against the reference proteomes for *X. laevis*^110^ (Xenbase version 10.1) and *D. melanogaster* acquired from Uniprot (UP000000803, September 2022) with common contaminants.

The target-decoy strategy was used to estimate the peptide false discovery rate (FDR)^111^, and a 1% FDR cutoff was used for MS2 spectral assignment. A 1Da precursor ion tolerance with the requirement that both N- and C-terminal peptide ends are consistent with the protease specificities of LysC and trypsin was used for SEQUEST searches. One missed cleavage was allowed, and NEM was set as a static modification of cysteine residues (+125.047679 Da). For TMTpro samples, TMTpro was set as a static modification of lysine residues and the peptide N-terminus (+304.2071 Da). Fragment ion tolerance in the MS2 spectrum was set at 1 Da. Filtering was performed using a linear discriminant analysis with the following features: Sequest parameters XCorr and unique ΔXCorr, peptide length, missed cleavages, adjusted PPM, fraction of ions matched and charge state. Forward peptides within 3 standard deviations of the theoretical m/z of the precursor were used as a positive training set. All reverse peptides were used as the negative training set. Linear discriminant scores were used to sort peptides with at least seven residues and to filter with the desired cutoff. Furthermore, we performed a filtering step on the protein level using the ‘picked’ protein FDR approach^112^. Protein redundancy was removed by assigning peptides to the minimal number of proteins, which can explain all observed peptides, with the above-described filtering criteria.

For reporter ion quantification, reporter ion intensities were extracted, and TMTpro isotopic impurity corrections were done. For 18plex samples, peptides with signal-to-noise ratios of at least 257 for 2+ peptides and 529 for 3+ peptides were retained for further analysis. Peptides with missed cleavages and isolation of non-M_0_ peaks were removed. Signal across peptide-spectrum matches (PSMs) was summed.

## DIA database search

Raw files were analyzed using DIA-NN v.2.5.0. A spectral library was predicted in silico from the FASTA files listed in DDA database search. DIA-NN searched the raw data against this predicted library, generated a refined spectral library from the DIA runs themselves, and re-searched the data against that library in a second pass with match-between-runs enabled. Cross-run retention-time alignment was applied.

*In silico* digestion assumed trypsin/LysC specificity with up to one missed cleavage, and peptides of 7 to 30 residues bearing charges of 1-4+ were considered. The precursor m/z range was restricted to 350-950 and the fragment m/z range to 200-1800. NEM was set as a fixed modification of cysteine (+125.047679 Da, UniMod:108) and methionine oxidation (+15.994915 Da, UniMod:35) as a variable modification, with at most one variable modification per precursor. Peptidoform scoring and PTM-site localization were enabled. Precursor identifications were scored against a target-decoy set using DIA-NN’s neural-network classifier and filtered to 1% FDR, with protein groups reported at a 1% global q-value. Mass accuracy and the scan window were determined automatically by DIA-NN, and quantification was performed in legacy (direct) mode.

## Wide isolation normalization method

The mass spectrometer was set to analyze positively charged ions in data-dependent mode, recording centroid data with the RF lens level at 60%. Full scans were taken with the Orbitrap at 120k resolution with an AGC target of 4 × 10^5^ charges, maximum IIT of 50 milliseconds, and a scan range of 350-1400 m/z with wide quadrupole isolation enabled. The maximum cycle time between MS1 scans was set to 3 seconds.

Following the survey scan, the following filters were applied for triggering MS2 scans. MIPS was enabled in peptide mode with the monoisotopic peak set as the isolation center, and precursor selection was restricted to 400-1200 m/z. Isolated masses were excluded for 60 seconds after a single triggering event with a mass tolerance window of ±10 ppm, excluding isotopes and alternate charge states of the isolated species. A charge-state filter was set for 2-6+ charge states.

The following settings were used for Orbitrap MS2 scans. Rather than narrowly isolating individual precursors, a 200 m/z quadrupole isolation window was applied so that the full isotopic envelope of any peptide-like precursor, together with co-eluting species across the window, was co-isolated and fragmented. Ions were fragmented with HCD at a normalized collision energy of 45%, and fragment spectra were recorded with the Orbitrap at 45k resolution with a first mass of 110 m/z, an AGC target of 2 × 10^5^ charges, and a maximum IIT of 91 milliseconds.

These acquisitions were not subjected to database searching. Because the wide (200 m/z) isolation window co-fragments effectively all peptide-like precursors in the window, the summed TMTpro reporter ion signal-to-noise in each channel provides an identification-free estimate of relative channel loading. This measurement was used solely to normalize loading across the multiplexed ^18^O samples, and no peptide or protein identifications were derived from these runs.

## Protein normalization

To avoid relying on assumptions about which proteins are constant, we ran companion samples with wide-isolation MS1 scans alongside the standard narrow-isolation scans used for degradation measurement (Figure S1a). Wide isolation captures the total reporter-ion signal across each multiplexed channel and provides an orthogonal loading control independent of which specific proteins are present.

We first applied this in Xenopus. Because its embryos contain enormous amounts of yolk protein^105^, we processed embryos without yolk depletion for these normalization experiments. The total reporter signal is then dominated by yolk and reports on overall channel loading. Wide-isolation analysis revealed a systematic decrease in total signal over the developmental time course, consistent with active yolk utilization across embryogenesis (Figure S1b). We fit an exponential model (EM) to the wide-isolation-adjusted yolk median and used the smoothed curve as a per-channel correction.

Our standard Xenopus preparation removes the yolk, so for the turnover samples, we instead defined a curated set of proteins with no measurable degradation or synthesis to serve as the normalization anchor. Proteins in the EM-normalized dataset were clustered by *k*-means (*k* = 5; Figure S1c), and the flat cluster was retained. From these, we kept proteins whose maximum-to-minimum relative abundance across the time course was below 1.2.

We then validated that these candidates were neither degraded nor synthesized by measuring their isotopic envelopes at the final ^18^O timepoint by label-free acquisition. Each observed envelope was modeled as a two-state mixture, *E_obs_ =* (1*-f*) *E_start_ + fE_End label_*, where *f*is the fraction of newly synthesized protein (Figure S1d). *E_start_* was constructed from the peptide elemental composition and *E_End label_* broadened to the MS1 resolution, and was generated by convolving per-residue ^18^O

incorporation probabilities across the sequence, with the number of incorporable oxygens set by amino acid class (Supplementary Methods). Each observed envelope was then fit by least squares to extract *f*. As positive controls, newly synthesized proteins including CIRBP and HMGB3 gave high *f*with close agreement between observed and predicted heavy envelopes (Figure S1e, S1f), while candidate reference proteins including HIBADH and PC showed flat trajectories and no detectable incorporation (*f* ≈ 0; Figure S1g, S1h). This yielded a final set of 169 reference proteins used as the normalization anchor for all yolk-depleted Xenopus datasets (Table S1, Figure S1d).

The Drosophila samples were corrected by wide isolation as described above. Because yolk removal is not part of the Drosophila preparation, the turnover samples themselves retain yolk, so the EM correction was applied directly (Figure S10), without the separate reference-protein anchor required for the yolk-depleted Xenopus samples.

## Model selection

To classify which proteins underwent statistically significant degradation, we compared each protein’s mass-balance fit to a no-degradation null in which the degradation rate was fixed at zero, using BIC. BIC was computed as *n*ln (*RSS*/*n*)+*m* ln *n*, where is the number of time points and the number of free parameters in each fitted model.

ΔBIC is a model-selection statistic rather than a test statistic and does not itself yield a *p*-value. We assign statistical significance from a permutation null, where each protein’s observed ΔBIC is compared against an empirical distribution by permuting a dataset’s time points and refitting (10 permutations per dataset). The resulting null ΔBIC values define the distribution achievable by chance, from which we calculate the FDR at a given ΔBIC cutoff (Figure S3). Permuted null fits rarely favored the degradation model, so the ΔBIC threshold needed to control the false-discovery rate fell below zero and would have classified proteins as degrading without positive evidence. Therefore, we applied a fixed cutoff of ΔBIC > 2, the conventional threshold for positive evidence^113^. For the turnover experiments, this is more conservative than false-discovery control alone would require.

Because the cell-cycle data analysis let the oscillatory and exponential models fit noise, the null produced appreciable false positives (Figure S3b, S3c), so controlling the false-discovery rate at 5% required stringent, model-specific thresholds of ΔBIC > 17.6 for the cosine model and ΔBIC > 11.1 for the exponential model. Proteins exceeding these thresholds were classified as oscillating or as directionally increasing/decreasing respectively, and all others as flat.

## Absolute protein quantification

Absolute protein abundances were estimated from the label-free DIA runs using an intensity Based Absolute Quantification (iBAQ) approach anchored to known total protein mass. DIA-NN precursor quantities were filtered to a 1% protein-group q-value, and contaminants and zero-intensity precursors were removed. For each protein group, we summed precursor intensities across its peptides and divided by the median number of theoretically observable tryptic peptides across its constituent proteins to give an iBAQ intensity proportional to molar abundance. Protein groups were retained only when their constituent proteins agreed to within 20% in both theoretical peptide count and sequence length.

To place these relative molar values on an absolute scale, iBAQ intensities were converted to mass fractions by weighting each protein by its molecular weight, and the mass fractions were scaled to published total protein amounts for each embryo. Xenopus required a two-compartment anchor because the samples were yolk-depleted by spinout. Since the yolk was only partially removed, its remaining fraction is not known *a priori* and cannot be placed on the same scale as the rest of the proteome, and analyzing a non-depleted sample would let the abundant yolk proteins dominate the signal and distort every other concentration. We therefore anchored the yolk and non-yolk compartments separately, scaling the yolk proteins to the yolk protein amount (∼280 μg) and all remaining proteins to the non-yolk amount (∼45 μg)^94^. Drosophila was scaled to a single total protein amount (∼0.8 μg) approximating the protein content at the start of the experiment^114^. The resulting per-protein masses were divided by molecular weight to yield molar concentrations, using the embryo volume to convert mass per embryo into concentration (∼10 nL for Drosophila^115^; ∼1 μL for Xenopus^116^).

## Feature and functional analysis

Each protein from each respective organism was annotated for function, subcellular localization, domain content, and intrinsic disorder, and these annotations were related to the fitted degradation rates. The analysis was performed in R.

For Xenopus, Gene Ontology (GO) terms^117,118^ were assigned through human orthologs, taking each protein’s biological-process, molecular-function, and cellular-component terms from its mapped human gene. Drosophila utilized its own annotated GO terms. To test whether turnover was organized by function, proteins were ranked by degradation rate and assessed for GO enrichment using fgsea^119^

(fgseaMultilevel), restricting to terms with more than 20 annotated proteins, collapsing redundant terms (collapsePathways), and reporting Benjamini–Hochberg-adjusted *p*-values. Subcellular compartments were derived from the cellular-component terms, grouping proteins into major compartments (nucleus, cytoplasm, mitochondrion, endoplasmic reticulum, and others) through the GO offspring relationships in GO.db, and degradation rates were summarized by compartment.

Protein domains were annotated with InterProScan^120,121^, run separately on each proteome, and half-lives were grouped and ranked by domain signature. Intrinsic disorder was predicted with IUPred2A^122^. Proteins with a mean IUPred score of at least 0.5 were classified as disordered and the rest as ordered. The inverse relationship between abundance and turnover was visualized by plotting half-life against measured absolute protein abundance.

## Homeolog analysis

Because the two Xenopus S and L homeologs are highly similar in sequence, shared peptides cannot distinguish them, so for this comparison we restricted the data to confidently assignable proteins. We used only proteins in the unique (U) parsimony class and re-quantified turnover from unique peptides alone, requiring at least two unique peptides per protein to reduce noise. Degradation rates were then compared between paired S and L homeologs taken from Xenbase annotations^110^.

## Kinetic class assignment

Each protein was assigned to one of five kinetic classes by combining its degradation status with the trend in its total abundance. Degradation status was taken from the BIC classification described above. The abundance trend was measured from the control channels as the fold change in total protein abundance between the first and last control time points. To bound the range of no measurable change, we used the curated reference set of proteins with no measurable degradation or synthesis defined above for Xenopus. Proteins whose control fold change exceeded the 99th percentile of this reference set were classified as increasing, those below the 1st percentile as decreasing, and the remainder as unchanging. Crossing degradation status with abundance trend yielded five classes: no measurable degradation (not degrading, abundance not increasing), protein synthesis only (not degrading, increasing), net protein increase with degradation (degrading, increasing), balanced protein turnover (degrading, unchanging), and protein degradation only (degrading, decreasing).

In Drosophila, nearly the entire proteome degrades, so no population shows no measurable change in both channels from which to define the unchanging range internally. We therefore applied the abundance-trend cutoffs derived from the Xenopus reference set (control fold changes of 1.44 and 0.62) directly to Drosophila, keeping the classification equivalent between the two species.

## Cross-species comparison

Xenopus and Drosophila turnover were compared at single-copy orthologs identified with OrthoFinder^89^. To assign each ortholog a high-quality rate, we required both members of a pair to be quantified in both replicates of their respective experiments, the Xenopus 2-cell and Drosophila gastrulation time courses.

## Amino acid mass-balance modeling

To estimate the nucleic acid synthesis demand in Xenopus over our time course, we derived net biosynthesis requirements from the literature. The Xenopus embryo begins with roughly 12 pg of DNA, which expands to nearly 1 µg by late embryonic stages, as inferred from the DNA content of normal embryos^123,124^. Studies of anucleolate mutants unable to synthesize new rRNA show that the wild-type embryo synthesizes approximately 6 µg of net rRNA by the early swimming stages^124^. The embryo therefore requires roughly 7 µg of net major nucleic acid synthesis by NF35, using later-stage DNA and rRNA measurements as a conservative upper bound. We treat the net accumulation of minor RNA species as small relative to rRNA^125^, since their combined net expansion is subsumed by our upper-bound DNA and rRNA estimates, and finally, mitochondrial RNA abundance is static within our window^126^. Given an average elemental nitrogen fraction of 15.5% for mixed biological nucleotides, synthesizing 7 µg of net nucleic acid requires almost 1.1 µg of atomic nitrogen.

We repeated the same approximation for Drosophila, again drawing net biosynthesis requirements from the literature. We begin our experimental window at roughly 3 hours post-fertilization, a stage comprising approximately 6,000 diploid nuclei^127^. Using a genome size of 180 megabases^128^, the diploid genome contains 0.36 pg of DNA which corresponds to 2.2 ng of starting DNA for our experiment. By hatching, the embryo contains roughly 50,000 cells^129^. The Drosophila embryo also enters the endocycle, in which specific tissues (salivary gland, midgut, hindgut, Malpighian tubules, and dorsal cells) replicate their genomes an average of two additional times without cell division^130^. Based on anatomical proportions, we estimate these tissues make up roughly 10% of the embryo. By hatching, the embryo therefore contains approximately 23.9 ng of DNA, a net synthesis of 21.7 ng across our window. We did not include a separate RNA demand for the Drosophila embryo. Available measurements indicate that this interval is dominated by remodeling of maternally supplied RNA pools rather than large net accumulation of bulk RNA, with classical measurements of major RNA-class synthesis and more recent mRNA time-course data supporting this approximation^131,132^. Using the 16.8% elemental nitrogen mass fraction of the Drosophila genome, synthesizing 21.7 ng of net DNA requires approximately 3.6 ng of atomic nitrogen.

With the nucleic acid demand in hand, we assembled a mass balance of the free amino acid pool from quantities we measured directly. New protein synthesis was taken from the absolute protein abundances we quantified, using the control channels, to estimate how much new protein was produced across the time course. The amino acids liberated by turnover of the inherited proteome were estimated by applying the fitted degradation rates to the starting protein amounts, giving the quantity released from the beginning of the window.

To allocate the atomic nitrogen demand from nucleic acids, we first established an intermediate pool by summing the yolk-derived and proteome-derived amino acids and subtracting the demand of *de novo* protein synthesis. Assuming rapid transaminase-driven equilibration, each amino acid’s contribution was weighted by its molarity in this intermediate pool and its atomic nitrogen count. These proportional nitrogen demands were then converted back into amino acid molarities, yielding the physiological amino acid demands for nucleic acid synthesis.

## Uric acid quantification

Drosophila embryos were collected as above. Following heat fixation, 1,000 embryos were resuspended in 125 μL PBS. Samples were sonicated at 50% amplitude for ten cycles of 10 seconds on and 20 seconds off, with cooling on ice between cycles, followed by incubation at 90°C for 5 minutes.

Lysates were then centrifuged through 3 kDa filters (Amicon Ultra, UFC500396) at 14,000 x g to isolate the filtrate containing uric acid (approximately 1 hour). Uric acid concentration was measured using a uric acid assay kit (Sigma-Aldrich, MAK483) according to the manufacturer’s instructions.

## Resource availability Lead contact

Requests for further information and resources should be directed to and will be fulfilled by the lead contact, Martin Wühr.

## Materials availability

This study did not generate new unique reagents.

## Data and code availability

- The mass spectrometry proteomics data have been deposited to the ProteomeXchange consortium^133^ via the PRIDE^134^ partner repository with the dataset identifier PXD081127 (DDA) and PXD084487 (DIA) and will be publicly available as of the date of publication. During review, the datasets can be accessed at https://www.ebi.ac.uk/pride/login with the reviewer access tokens FQZwMc2nWysn (PXD081127) and nLQgg4gCKOoo (PXD084487).
- All original code is publicly available at https://github.com/wuhrlab/18O-protein-turnover-embryos and will be archived with a DOI at Zenodo as of the date of publication. The repository contains the analysis pipeline used to derive protein turnover rates from the deposited data: conversion of free amino acid isotopologue measurements into peptide light-synthesis probabilities, joint fitting of the control and ^18^O time courses to the mass-balance model to estimate degradation (k_d_) and synthesis (k_T_) rates, BIC-based model selection with the permutation null used to classify degrading proteins, the wide-isolation and reference-protein normalization procedures, and the amino acid mass-balance models. Peptide identification and reporter-ion quantification were performed with GFY, which is licensed from Harvard University and cannot be redistributed by the authors. All search parameters are specified in Methods.
- Any additional information required to reanalyze the data reported in this paper is available from the lead contact upon request.

## Author contributions

MWK conceived the use of ^18^O water-based metabolic labeling. ERC, EW, and MW designed the study. AM performed initial experiments and contributed to the conceptual development of the project. ERC and GB performed most experiments and data analysis. ANTJ and FCK contributed to developing the modeling. VP assisted with setting up experiments and data analysis. DI and MN performed the metabolomics. JDR, EW, and MW supervised the study and provided funding. ERC, EW, and MW wrote the manuscript with input from all authors.

## Acknowledgements

We thank Andrea Mariossi and Trudi Schüpbach for useful advice and discussions. We gratefully acknowledge support from the NSF Graduate Research Fellowship (ERC), Ruth L. Kirschstein NRSA F30 F30GM154411 (AM), Princeton University’s Summer Undergraduate Research Program (GB, VP), NIH R35GM128813 (MW), the Princeton Catalysis Initiative (MW), and the Eric and Wendy Schmidt Transformative Technology Fund (MW).

## Declaration of generative AI and AI-assisted technologies in the manuscript preparation process

During the preparation of this work, the authors used Claude (Anthropic) to assist with drafting and editing the text, literature searching, and cross-checking the manuscript against the analysis code. After using this tool, the authors reviewed and edited the content as needed and take full responsibility for the content of the published article.

## References

1. Aviles-Pagan, E.E., and Orr-Weaver, T.L. (2018). Activating embryonic development in Drosophila. Semin Cell Dev Biol 84, 100–110. 10.1016/j.semcdb.2018.02.019.

2. Stitzel, M.L., and Seydoux, G. (2007). Regulation of the oocyte-to-zygote transition. Science 316, 407–408. 10.1126/science.1138236.

3. Tadros, W., and Lipshitz, H.D. (2009). The maternal-to-zygotic transition: a play in two acts. Development 136, 3033–3042. 10.1242/dev.033183.

4. Vastenhouw, N.L., Cao, W.X., and Lipshitz, H.D. (2019). The maternal-to-zygotic transition revisited. Development 146. 10.1242/dev.161471.

5. Kojima, M.L., Hoppe, C., and Giraldez, A.J. (2025). The maternal-to-zygotic transition: reprogramming of the cytoplasm and nucleus. Nat Rev Genet 26, 245–267. 10.1038/s41576-024-00792-0.

6. Tadros, W., Goldman, A.L., Babak, T., Menzies, F., Vardy, L., Orr-Weaver, T., Hughes, T.R., Westwood, J.T., Smibert, C.A., and Lipshitz, H.D. (2007). SMAUG is a major regulator of maternal mRNA destabilization in Drosophila and its translation is activated by the PAN GU kinase. Dev Cell 12, 143–155. 10.1016/j.devcel.2006.10.005.

7. Chen, L., Dumelie, J.G., Li, X., Cheng, M.H., Yang, Z., Laver, J.D., Siddiqui, N.U., Westwood, J.T., Morris, Q., Lipshitz, H.D., and Smibert, C.A. (2014). Global regulation of mRNA translation and stability in the early Drosophila embryo by the Smaug RNA-binding protein. Genome Biol 15, R4. 10.1186/gb-2014-15-1-r4.

8. Laver, J.D., Li, X., Ray, D., Cook, K.B., Hahn, N.A., Nabeel-Shah, S., Kekis, M., Luo, H., Marsolais, A.J., Fung, K.Y., et al. (2015). Brain tumor is a sequence-specific RNA-binding protein that directs maternal mRNA clearance during the Drosophila maternal-to-zygotic transition. Genome Biol 16, 94. 10.1186/s13059-015-0659-4.

9. Bushati, N., Stark, A., Brennecke, J., and Cohen, S.M. (2008). Temporal reciprocity of miRNAs and their targets during the maternal-to-zygotic transition in Drosophila. Curr Biol 18, 501–506. 10.1016/j.cub.2008.02.081.

10. Lund, E., Liu, M., Hartley, R.S., Sheets, M.D., and Dahlberg, J.E. (2009). Deadenylation of maternal mRNAs mediated by miR-427 in Xenopus laevis embryos. RNA 15, 2351–2363. 10.1261/rna.1882009.

11. Peshkin, L., Wuhr, M., Pearl, E., Haas, W., Freeman, R.M., Jr., Gerhart, J.C., Klein, A.M., Horb, M., Gygi, S.P., and Kirschner, M.W. (2015). On the Relationship of Protein and mRNA Dynamics in Vertebrate Embryonic Development. Dev Cell 35, 383–394. 10.1016/j.devcel.2015.10.010.

12. Frese, A.N., Mariossi, A., Levine, M.S., and Wuhr, M. (2024). Quantitative proteome dynamics across embryogenesis in a model chordate. iScience 27, 109355. 10.1016/j.isci.2024.109355.

13. Subtelny, A.O., Eichhorn, S.W., Chen, G.R., Sive, H., and Bartel, D.P. (2014). Poly(A)-tail profiling reveals an embryonic switch in translational control. Nature 508, 66–71. 10.1038/nature13007.

14. Leesch, F., Lorenzo-Orts, L., Pribitzer, C., Grishkovskaya, I., Roehsner, J., Chugunova, A., Matzinger, M., Roitinger, E., Belacic, K., Kandolf, S., et al. (2023). A molecular network of conserved factors keeps ribosomes dormant in the egg. Nature 613, 712–720. 10.1038/s41586-022-05623-y.

15. Casas-Vila, N., Bluhm, A., Sayols, S., Dinges, N., Dejung, M., Altenhein, T., Kappei, D., Altenhein, B., Roignant, J.Y., and Butter, F. (2017). The developmental proteome of Drosophila melanogaster. Genome Res 27, 1273–1285. 10.1101/gr.213694.116.

16. da Silva Pescador, G., Baia Amaral, D., Varberg, J.M., Zhang, Y., Hao, Y., Florens, L., and Bazzini, A.A. (2024). Protein profiling of zebrafish embryos unmasks regulatory layers during early embryogenesis. Cell Rep 43, 114769. 10.1016/j.celrep.2024.114769.

17. Toralova, T., Kinterova, V., Chmelikova, E., and Kanka, J. (2020). The neglected part of early embryonic development: maternal protein degradation. Cell Mol Life Sci 77, 3177–3194. 10.1007/s00018-020-03482-2.

18. Cao, W.X., Kabelitz, S., Gupta, M., Yeung, E., Lin, S., Rammelt, C., Ihling, C., Pekovic, F., Low, T.C.H., Siddiqui, N.U., et al. (2020). Precise Temporal Regulation of Post-transcriptional Repressors Is Required for an Orderly Drosophila Maternal-to-Zygotic Transition. Cell Rep 31, 107783. 10.1016/j.celrep.2020.107783.

19. Cao, W.X., Karaiskakis, A., Lin, S., Angers, S., and Lipshitz, H.D. (2022). The F-box protein Bard (CG14317) targets the Smaug RNA-binding protein for destruction during the Drosophila maternal-to-zygotic transition. Genetics 220. 10.1093/genetics/iyab177.

20. Zavortink, M., Rutt, L.N., Dzitoyeva, S., Henriksen, J.C., Barrington, C., Bilodeau, D.Y., Wang, M., Chen, X.X.L., and Rissland, O.S. (2020). The E2 Marie Kondo and the CTLH E3 ligase clear deposited RNA binding proteins during the maternal-to-zygotic transition. Elife 9. 10.7554/eLife.53889.

21. Kronja, I., Whitfield, Z.J., Yuan, B., Dzeyk, K., Kirkpatrick, J., Krijgsveld, J., and Orr-Weaver, T.L. (2014). Quantitative proteomics reveals the dynamics of protein changes during Drosophila oocyte maturation and the oocyte-to-embryo transition. Proc Natl Acad Sci U S A 111, 16023–16028. 10.1073/pnas.1418657111.

22. Kronja, I., Yuan, B., Eichhorn, S.W., Dzeyk, K., Krijgsveld, J., Bartel, D.P., and Orr-Weaver, T.L. (2014). Widespread changes in the posttranscriptional landscape at the Drosophila oocyte-to-embryo transition. Cell Rep 7, 1495–1508. 10.1016/j.celrep.2014.05.002.

23. Presler, M., Van Itallie, E., Klein, A.M., Kunz, R., Coughlin, M.L., Peshkin, L., Gygi, S.P., Wuhr, M., and Kirschner, M.W. (2017). Proteomics of phosphorylation and protein dynamics during fertilization and meiotic exit in the Xenopus egg. Proc Natl Acad Sci U S A 114, E10838–E10847. 10.1073/pnas.1709207114.

24. Schoenheimer, R., and Rittenberg, D. (1935). Deuterium as an Indicator in the Study of Intermediary Metabolism. Science 82, 156–157. 10.1126/science.82.2120.156.

25. Schoenheimer, R. (1942). The Dynamic State of Body Constituents (Harvard University Press).

26. Claydon, A.J., and Beynon, R. (2012). Proteome dynamics: revisiting turnover with a global perspective. Mol Cell Proteomics 11, 1551–1565. 10.1074/mcp.O112.022186.

27. Ross, A.B., Langer, J.D., and Jovanovic, M. (2021). Proteome Turnover in the Spotlight: Approaches, Applications, and Perspectives. Mol Cell Proteomics 20, 100016. 10.1074/mcp.R120.002190.

28. Schwanhausser, B., Busse, D., Li, N., Dittmar, G., Schuchhardt, J., Wolf, J., Chen, W., and Selbach, M. (2011). Global quantification of mammalian gene expression control. Nature 473, 337–342. 10.1038/nature10098.

29. Mathieson, T., Franken, H., Kosinski, J., Kurzawa, N., Zinn, N., Sweetman, G., Poeckel, D., Ratnu, V.S., Schramm, M., Becher, I., et al. (2018). Systematic analysis of protein turnover in primary cells. Nat Commun 9, 689. 10.1038/s41467-018-03106-1.

30. Rolfs, Z., Frey, B.L., Shi, X., Kawai, Y., Smith, L.M., and Welham, N.V. (2021). An atlas of protein turnover rates in mouse tissues. Nat Commun 12, 6778. 10.1038/s41467-021-26842-3.

31. Harasimov, K., Gorry, R.L., Welp, L.M., Penir, S.M., Horokhovskyi, Y., Cheng, S., Takaoka, K., Stutzer, A., Frombach, A.S., Taylor Tavares, A.L., et al. (2024). The maintenance of oocytes in the mammalian ovary involves extreme protein longevity. Nat Cell Biol 26, 1124–1138. 10.1038/s41556-024-01442-7.

32. Gupta, M., Johnson, A.N.T., Cruz, E.R., Costa, E.J., Guest, R.L., Li, S.H., Hart, E.M., Nguyen, T., Stadlmeier, M., Bratton, B.P., et al. (2024). Global protein turnover quantification in Escherichia coli reveals cytoplasmic recycling under nitrogen limitation. Nat Commun 15, 5890. 10.1038/s41467-024-49920-8.

33. Rayon, T., Stamataki, D., Perez-Carrasco, R., Garcia-Perez, L., Barrington, C., Melchionda, M., Exelby, K., Lazaro, J., Tybulewicz, V.L.J., Fisher, E.M.C., and Briscoe, J. (2020). Species-specific pace of development is associated with differences in protein stability. Science 369, eaba7667. 10.1126/science.aba7667.

34. Matsuda, M., Hammarén, H.M., Lázaro, J., Savitski, M.M., and Ebisuya, M. (2026). Systematic differences in protein stability underlie species-specific developmental tempo. Dev Cell 61, 1855–1866. 10.1016/j.devcel.2026.07.012.

35. Nakanoh, S., Stamataki, D., Garcia-Perez, B., Azzi, C., Carr, H.L., Pokhilko, A., Yu, L., Doshi, L., Boezio, L.M., Melchionda, M., et al. (2026). Proteasome-dependent protein degradation shapes developmental tempo in mouse and human neural progenitors. Dev Cell 61, 1867–1882. 10.1016/j.devcel.2026.07.014.

36. Vastag, L., Jorgensen, P., Peshkin, L., Wei, R., Rabinowitz, J.D., and Kirschner, M.W. (2011). Remodeling of the metabolome during early frog development. PLoS One 6, e16881. 10.1371/journal.pone.0016881.

37. Sadygov, R.G. (2022). Protein turnover models for LC-MS data of heavy water metabolic labeling. Brief Bioinform 23. 10.1093/bib/bbab598.

38. Gross, P.R., and Spindel, W. (1960). Mitotic arrest by deuterium oxide. Science 131, 37–38. 10.1126/science.131.3392.37.

39. Scharf, S.R., Rowning, B., Wu, M., and Gerhart, J.C. (1989). Hyperdorsoanterior embryos from Xenopus eggs treated with D2O. Dev Biol 134, 175–188. 10.1016/0012-1606(89)90087-0.

40. Calve, S., Witten, A.J., Ocken, A.R., and Kinzer-Ursem, T.L. (2016). Incorporation of non-canonical amino acids into the developing murine proteome. Sci Rep 6, 32377. 10.1038/srep32377.

41. Saleh, A.M., Jacobson, K.R., Kinzer-Ursem, T.L., and Calve, S. (2019). Dynamics of Non-Canonical Amino Acid-Labeled Intra- and Extracellular Proteins in the Developing Mouse. Cell Mol Bioeng 12, 495–509. 10.1007/s12195-019-00592-1.

42. Bernlohr, R.W. (1972). 18Oxygen probes of protein turnover, amino acid transport, and protein synthesis in Bacillus licheniformis. J Biol Chem 247, 4893–4899. 10.1016/S0021-9258(19)44994-6.

43. Li, J., Cai, Z., Bomgarden, R.D., Pike, I., Kuhn, K., Rogers, J.C., Roberts, T.M., Gygi, S.P., and Paulo, J.A. (2021). TMTpro-18plex: The Expanded and Complete Set of TMTpro Reagents for Sample Multiplexing. J Proteome Res 20, 2964–2972. 10.1021/acs.jproteome.1c00168.

44. Erickson, B.K., Mintseris, J., Schweppe, D.K., Navarrete-Perea, J., Erickson, A.R., Nusinow, D.P., Paulo, J.A., and Gygi, S.P. (2019). Active Instrument Engagement Combined with a Real-Time Database Search for Improved Performance of Sample Multiplexing Workflows. J Proteome Res 18, 1299–1306. 10.1021/acs.jproteome.8b00899.

45. Schweppe, D.K., Eng, J.K., Yu, Q., Bailey, D., Rad, R., Navarrete-Perea, J., Huttlin, E.L., Erickson, B.K., Paulo, J.A., and Gygi, S.P. (2020). Full-Featured, Real-Time Database Searching Platform Enables Fast and Accurate Multiplexed Quantitative Proteomics. J Proteome Res 19, 2026–2034. 10.1021/acs.jproteome.9b00860.

46. Zahn, N., James-Zorn, C., Ponferrada, V.G., Adams, D.S., Grzymkowski, J., Buchholz, D.R., Nascone-Yoder, N.M., Horb, M., Moody, S.A., Vize, P.D., and Zorn, A.M. (2022). Normal Table of Xenopus development: a new graphical resource. Development 149. 10.1242/dev.200356.

47. Payne, S.H., and Loomis, W.F. (2006). Retention and loss of amino acid biosynthetic pathways based on analysis of whole-genome sequences. Eukaryot Cell 5, 272–276. 10.1128/EC.5.2.272-276.2006.

48. Crapse, J., Pappireddi, N., Gupta, M., Shvartsman, S.Y., Wieschaus, E., and Wuhr, M. (2021). Evaluating the Arrhenius equation for developmental processes. Mol Syst Biol 17, e9895. 10.15252/msb.20209895.

49. Funabiki, H., and Murray, A.W. (2000). The Xenopus chromokinesin Xkid is essential for metaphase chromosome alignment and must be degraded to allow anaphase chromosome movement. Cell 102, 411–424.

50. Castro, A., Vigneron, S., Bernis, C., Labbe, J.C., and Lorca, T. (2003). Xkid is degraded in a D-box, KEN-box, and A-box-independent pathway. Mol Cell Biol 23, 4126–4138. 10.1128/MCB.23.12.4126-4138.2003.

51. Salic, A., Waters, J.C., and Mitchison, T.J. (2004). Vertebrate shugoshin links sister centromere cohesion and kinetochore microtubule stability in mitosis. Cell 118, 567–578.

52. Fu, G., Hua, S., Ward, T., Ding, X., Yang, Y., Guo, Z., and Yao, X. (2007). D-box is required for the degradation of human Shugoshin and chromosome alignment. Biochem Biophys Res Commun 357, 672–678. 10.1016/j.bbrc.2007.03.204.

53. Karamysheva, Z., Diaz-Martinez, L.A., Crow, S.E., Li, B., and Yu, H. (2009). Multiple anaphase-promoting complex/cyclosome degrons mediate the degradation of human Sgo1. J Biol Chem 284, 1772–1780. 10.1074/jbc.M807083200.

54. Kitajima, T.S., Sakuno, T., Ishiguro, K., Iemura, S., Natsume, T., Kawashima, S.A., and Watanabe, Y. (2006). Shugoshin collaborates with protein phosphatase 2A to protect cohesin. Nature 441, 46–52. 10.1038/nature04663.

55. Liu, H., Rankin, S., and Yu, H. (2013). Phosphorylation-enabled binding of SGO1-PP2A to cohesin protects sororin and centromeric cohesion during mitosis. Nat Cell Biol 15, 40–49. 10.1038/ncb2637.

56. Rogers, C.D., Harafuji, N., Archer, T., Cunningham, D.D., and Casey, E.S. (2009). Xenopus Sox3 activates sox2 and geminin and indirectly represses Xvent2 expression to induce neural progenitor formation at the expense of non-neural ectodermal derivatives. Mech Dev 126, 42–55. 10.1016/j.mod.2008.10.005.

57. Hendrickson, C.L., Blitz, I.L., Hussein, A., Paraiso, K.D., Cho, J.S., Klymkowsky, M.W., Kofron, M.J., and Cho, K.W.Y. (2025). Foxi2 and Sox3 are master transcription regulators that control ectoderm germ layer specification in Xenopus. PLoS Biol 23, e3003476. 10.1371/journal.pbio.3003476.

58. Penzel, R., Oschwald, R., Chen, Y., Tacke, L., and Grunz, H. (1997). Characterization and early embryonic expression of a neural specific transcription factor xSOX3 in Xenopus laevis. Int J Dev Biol 41, 667–677.

59. Rogers, C.D., Archer, T.C., Cunningham, D.D., Grammer, T.C., and Casey, E.M. (2008). Sox3 expression is maintained by FGF signaling and restricted to the neural plate by Vent proteins in the Xenopus embryo. Dev Biol 313, 307–319. 10.1016/j.ydbio.2007.10.023.

60. Heasman, J., Ginsberg, D., Geiger, B., Goldstone, K., Pratt, T., Yoshida-Noro, C., and Wylie, C. (1994). A functional test for maternally inherited cadherin in Xenopus shows its importance in cell adhesion at the blastula stage. Development 120, 49–57. 10.1242/dev.120.1.49.

61. Kuroda, H., Inui, M., Sugimoto, K., Hayata, T., and Asashima, M. (2002). Axial protocadherin is a mediator of prenotochord cell sorting in Xenopus. Dev Biol 244, 267–277. 10.1006/dbio.2002.0589.

62. Hegazy, M., Perl, A.L., Svoboda, S.A., and Green, K.J. (2022). Desmosomal Cadherins in Health and Disease. Annu Rev Pathol 17, 47–72. 10.1146/annurev-pathol-042320-092912.

63. Nandadasa, S., Tao, Q., Menon, N.R., Heasman, J., and Wylie, C. (2009). N- and E-cadherins in Xenopus are specifically required in the neural and non-neural ectoderm, respectively, for F-actin assembly and morphogenetic movements. Development 136, 1327–1338. 10.1242/dev.031203.

64. Session, A.M., Uno, Y., Kwon, T., Chapman, J.A., Toyoda, A., Takahashi, S., Fukui, A., Hikosaka, A., Suzuki, A., Kondo, M., et al. (2016). Genome evolution in the allotetraploid frog Xenopus laevis. Nature 538, 336–343. 10.1038/nature19840.

65. Bass-Zubek, A.E., Hobbs, R.P., Amargo, E.V., Garcia, N.J., Hsieh, S.N., Chen, X., Wahl, J.K., 3rd, Denning, M.F., and Green, K.J. (2008). Plakophilin 2: a critical scaffold for PKC alpha that regulates intercellular junction assembly. J Cell Biol 181, 605–613. 10.1083/jcb.200712133.

66. Fishbain, S., Inobe, T., Israeli, E., Chavali, S., Yu, H., Kago, G., Babu, M.M., and Matouschek, A. (2015). Sequence composition of disordered regions fine-tunes protein half-life. Nat Struct Mol Biol 22, 214–221. 10.1038/nsmb.2958.

67. van der Lee, R., Lang, B., Kruse, K., Gsponer, J., Sanchez de Groot, N., Huynen, M.A., Matouschek, A., Fuxreiter, M., and Babu, M.M. (2014). Intrinsically disordered segments affect protein half-life in the cell and during evolution. Cell Rep 8, 1832–1844. 10.1016/j.celrep.2014.07.055.

68. Murray, A.W., and Kirschner, M.W. (1989). Cyclin synthesis drives the early embryonic cell cycle. Nature 339, 275–280.

69. Hartley, R.S., Rempel, R.E., and Maller, J.L. (1996). In vivo regulation of the early embryonic cell cycle in Xenopus. Dev Biol 173, 408–419. 10.1006/dbio.1996.0036.

70. McGarry, T.J., and Kirschner, M.W. (1998). Geminin, an inhibitor of DNA replication, is degraded during mitosis. Cell 93, 1043–1053.

71. Litscher, E.S., and Wassarman, P.M. (2014). Evolution, structure, and synthesis of vertebrate egg-coat proteins. Trends Dev Biol 8, 65–76.

72. Lang, T., Klasson, S., Larsson, E., Johansson, M.E., Hansson, G.C., and Samuelsson, T. (2016). Searching the Evolutionary Origin of Epithelial Mucus Protein Components-Mucins and FCGBP. Mol Biol Evol 33, 1921–1936. 10.1093/molbev/msw066.

73. Vandooren, J., and Itoh, Y. (2021). Alpha-2-Macroglobulin in Inflammation, Immunity and Infections. Front Immunol 12, 803244. 10.3389/fimmu.2021.803244.

74. Newport, J., and Kirschner, M. (1982). A major developmental transition in early Xenopus embryos: II. Control of the onset of transcription. Cell 30, 687–696. 10.1016/0092-8674(82)90273-2.

75. Tsai, T.Y., Theriot, J.A., and Ferrell, J.E., Jr. (2014). Changes in oscillatory dynamics in the cell cycle of early Xenopus laevis embryos. PLoS Biol 12, e1001788. 10.1371/journal.pbio.1001788.

76. Collart, C., Owens, N.D., Bhaw-Rosun, L., Cooper, B., De Domenico, E., Patrushev, I., Sesay, A.K., Smith, J.N., Smith, J.C., and Gilchrist, M.J. (2014). High-resolution analysis of gene activity during the Xenopus mid-blastula transition. Development 141, 1927–1939. 10.1242/dev.102012.

77. Newport, J., and Kirschner, M. (1982). A major developmental transition in early Xenopus embryos: I. characterization and timing of cellular changes at the midblastula stage. Cell 30, 675–686. 10.1016/0092-8674(82)90272-0.

78. Tsuyama, T., Tada, S., Watanabe, S., Seki, M., and Enomoto, T. (2005). Licensing for DNA replication requires a strict sequential assembly of Cdc6 and Cdt1 onto chromatin in Xenopus egg extracts. Nucleic Acids Res 33, 765–775. 10.1093/nar/gki226.

79. Arias, E.E., and Walter, J.C. (2005). Replication-dependent destruction of Cdt1 limits DNA replication to a single round per cell cycle in Xenopus egg extracts. Genes Dev 19, 114–126. 10.1101/gad.1255805.

80. Li, A., and Blow, J.J. (2005). Cdt1 downregulation by proteolysis and geminin inhibition prevents DNA re-replication in Xenopus. EMBO J 24, 395–404. 10.1038/sj.emboj.7600520.

81. Kisielewska, J., and Blow, J.J. (2012). Dynamic interactions of high Cdt1 and geminin levels regulate S phase in early Xenopus embryos. Development 139, 63–74. 10.1242/dev.068676.

82. Saxe, J.P., Chen, M., Zhao, H., and Lin, H. (2013). Tdrkh is essential for spermatogenesis and participates in primary piRNA biogenesis in the germline. EMBO J 32, 1869–1885. 10.1038/emboj.2013.121.

83. Hales, K.G., Korey, C.A., Larracuente, A.M., and Roberts, D.M. (2015). Genetics on the Fly: A Primer on the Drosophila Model System. Genetics 201, 815–842. 10.1534/genetics.115.183392.

84. Cox, R.T., Kirkpatrick, C., and Peifer, M. (1996). Armadillo is required for adherens junction assembly, cell polarity, and morphogenesis during Drosophila embryogenesis. J Cell Biol 134, 133–148. 10.1083/jcb.134.1.133.

85. Orsulic, S., and Peifer, M. (1996). An in vivo structure-function study of armadillo, the beta-catenin homologue, reveals both separate and overlapping regions of the protein required for cell adhesion and for wingless signaling. J Cell Biol 134, 1283–1300. 10.1083/jcb.134.5.1283.

86. Kiffin, R., Christian, C., Knecht, E., and Cuervo, A.M. (2004). Activation of chaperone-mediated autophagy during oxidative stress. Mol Biol Cell 15, 4829–4840. 10.1091/mbc.e04-06-0477.

87. Tsuchiya, Y., Yamaguchi, M., Chikuma, T., and Hojo, H. (2005). Degradation of glyceraldehyde-3-phosphate dehydrogenase triggered by 4-hydroxy-2-nonenal and 4-hydroxy-2-hexenal. Arch Biochem Biophys 438, 217–222. 10.1016/j.abb.2005.04.015.

88. Brown, J.B., Boley, N., Eisman, R., May, G.E., Stoiber, M.H., Duff, M.O., Booth, B.W., Wen, J., Park, S., Suzuki, A.M., et al. (2014). Diversity and dynamics of the Drosophila transcriptome. Nature 512, 393–399. 10.1038/nature12962.

89. Emms, D.M., and Kelly, S. (2019). OrthoFinder: phylogenetic orthology inference for comparative genomics. Genome Biol 20, 238. 10.1186/s13059-019-1832-y.

90. Carlisle, E., Yin, Z., Pisani, D., and Donoghue, P.C.J. (2024). Ediacaran origin and Ediacaran-Cambrian diversification of Metazoa. Sci Adv 10, eadp7161. 10.1126/sciadv.adp7161.

91. Kuhn, H., Sopko, R., Coughlin, M., Perrimon, N., and Mitchison, T. (2015). The Atg1-Tor pathway regulates yolk catabolism in Drosophila embryos. Development 142, 3869–3878. 10.1242/dev.125419.

92. Tennessen, J.M., Bertagnolli, N.M., Evans, J., Sieber, M.H., Cox, J., and Thummel, C.S. (2014). Coordinated metabolic transitions during Drosophila embryogenesis and the onset of aerobic glycolysis. G3 (Bethesda) 4, 839–850. 10.1534/g3.114.010652.

93. Song, Y., Park, J.O., Tanner, L., Nagano, Y., Rabinowitz, J.D., and Shvartsman, S.Y. (2019). Energy budget of Drosophila embryogenesis. Curr Biol 29, R566–R567. 10.1016/j.cub.2019.05.025.

94. Van Itallie, E., Sonnett, M., Kalocsay, M., Wuhr, M., Peshkin, L., and Kirschner, M.W. (2025). Transitions in the proteome and phospho-proteome during Xenopus laevis development. Dev Biol 525, 155–171. 10.1016/j.ydbio.2025.05.022.

95. Owens, N.D.L., Blitz, I.L., Lane, M.A., Patrushev, I., Overton, J.D., Gilchrist, M.J., Cho, K.W.Y., and Khokha, M.K. (2016). Measuring Absolute RNA Copy Numbers at High Temporal Resolution Reveals Transcriptome Kinetics in Development. Cell Rep 14, 632–647. 10.1016/j.celrep.2015.12.050.

96. Kilwein, M.D., Johnson, M.R., Thomalla, J.M., Mahowald, A.P., and Welte, M.A. (2023). Drosophila embryos spatially sort their nutrient stores to facilitate their utilization. Development 150. 10.1242/dev.201423.

97. Woodland, H.R., and Pestell, R.Q. (1972). Determination of the nucleoside triphosphate contents of eggs and oocytes of Xenopus laevis. Biochem J 127, 597–605. 10.1042/bj1270597.

98. Jorgensen, P., Steen, J.A., Steen, H., and Kirschner, M.W. (2009). The mechanism and pattern of yolk consumption provide insight into embryonic nutrition in Xenopus. Development 136, 1539–1548. 136/9/1539 [pii] 10.1242/dev.032425.

99. Rogers, K.W., Bläßle, A., Schier, A.F., and Müller, P. (2015). Measuring protein stability in living zebrafish embryos using fluorescence decay after photoconversion (FDAP). J Vis Exp, 52266. 10.3791/52266.

100. Miller, S.E., Wang, T.Y., Quan, B., Cammidge, T., Chou, T.F., Prober, D.A., and Tirrell, D.A. (2026). Time-resolved proteomic analysis in zebrafish using bioorthogonal noncanonical amino acid tagging. J Proteome Res 25, 2263. 10.1021/acs.jproteome.5c00845.

101. Pino, L.K., Baeza, J., Lauman, R., Schilling, B., and Garcia, B.A. (2021). Improved SILAC Quantification with Data-Independent Acquisition to Investigate Bortezomib-Induced Protein Degradation. J Proteome Res 20, 1918–1927. 10.1021/acs.jproteome.0c00938.

102. Sabatier, P., Lechner, M., Guzman, U.H., Beusch, C.M., Zeng, X., Wang, L., Izaguirre, F., Seth, A., Gritsenko, O., Rodin, S., et al. (2025). Global analysis of protein turnover dynamics in single cells. Cell 188, 2433–2450 e2421. 10.1016/j.cell.2025.03.002.

103. Sive, H.L., Grainger, R.M., and Harland, R.M. (2000). Early Development of Xenopus Laevis: A Laboratory Manual (Cold Spring Harbor Laboratory Press).

104. Nieuwkoop, P.D., and Faber, J. (1994). Normal Table of Xenopus laevis (Daudin): A Systematical and Chronological Survey of the Development from the Fertilized Egg till the End of Metamorphosis, 1 Edition (Garland Science). 10.1201/9781003064565.

105. Gupta, M., Sonnett, M., Ryazanova, L., Presler, M., and Wuhr, M. (2018). Quantitative Proteomics of Xenopus Embryos I, Sample Preparation. Methods Mol Biol 1865, 175–194. 10.1007/978-1-4939-8784-9_13.

106. Hughes, C.S., Moggridge, S., Muller, T., Sorensen, P.H., Morin, G.B., and Krijgsveld, J. (2019). Single-pot, solid-phase-enhanced sample preparation for proteomics experiments. Nat Protoc 14, 68–85. 10.1038/s41596-018-0082-x.

107. Edwards, A., and Haas, W. (2016). Multiplexed Quantitative Proteomics for High-Throughput Comprehensive Proteome Comparisons of Human Cell Lines. Methods Mol Biol 1394, 1–13. 10.1007/978-1-4939-3341-9_1.

108. Munger, J., Bennett, B.D., Parikh, A., Feng, X.J., McArdle, J., Rabitz, H.A., Shenk, T., and Rabinowitz, J.D. (2008). Systems-level metabolic flux profiling identifies fatty acid synthesis as a target for antiviral therapy. Nat Biotechnol 26, 1179–1186. 10.1038/nbt.1500.

109. Mitulovic, G., Stingl, C., Steinmacher, I., Hudecz, O., Hutchins, J.R., Peters, J.M., and Mechtler, K. (2009). Preventing carryover of peptides and proteins in nano LC-MS separations. Anal Chem 81, 5955–5960. 10.1021/ac900696m.

110. Fisher, M., James-Zorn, C., Ponferrada, V., Bell, A.J., Sundararaj, N., Segerdell, E., Chaturvedi, P., Bayyari, N., Chu, S., Pells, T., et al. (2023). Xenbase: key features and resources of the Xenopus model organism knowledgebase. Genetics 224. 10.1093/genetics/iyad018.

111. Elias, J.E., and Gygi, S.P. (2007). Target-decoy search strategy for increased confidence in large-scale protein identifications by mass spectrometry. Nat Methods 4, 207–214. nmeth1019 [pii] 10.1038/nmeth1019.

112. Savitski, M.M., Wilhelm, M., Hahne, H., Kuster, B., and Bantscheff, M. (2015). A Scalable Approach for Protein False Discovery Rate Estimation in Large Proteomic Data Sets. Mol Cell Proteomics 14, 2394–2404. 10.1074/mcp.M114.046995.

113. Kass, R.E., and Raftery, A.E. (1995). Bayes Factors. Journal of the American Statistical Association 90, 773–795. 10.1080/01621459.1995.10476572.

114. Medina, M., and Vallejo, C.G. (1989). The contents of proteins, carbohydrates, lipids and DNA during the embryogenesis of Drosophila. Int J Dev Biol 33, 403–405.

115. Miles, C.M., Lott, S.E., Hendriks, C.L., Ludwig, M.Z., Manu, Williams, C.L., and Kreitman, M. (2011). Artificial selection on egg size perturbs early pattern formation in Drosophila melanogaster. Evolution 65, 33–42. 10.1111/j.1558-5646.2010.01088.x.

116. Mitchison, T.J., Ishihara, K., Nguyen, P., and Wuhr, M. (2015). Size Scaling of Microtubule Assemblies in Early Xenopus Embryos. Cold Spring Harb Perspect Biol 7, a019182. 10.1101/cshperspect.a019182.

117. Gene Ontology, C. (2026). The Gene Ontology knowledgebase in 2026. Nucleic Acids Res 54, D1779–D1792. 10.1093/nar/gkaf1292.

118. Ashburner, M., Ball, C.A., Blake, J.A., Botstein, D., Butler, H., Cherry, J.M., Davis, A.P., Dolinski, K., Dwight, S.S., Eppig, J.T., et al. (2000). Gene ontology: tool for the unification of biology. The Gene Ontology Consortium. Nat Genet 25, 25–29. 10.1038/75556.

119. Korotkevich, G., Sukhov, V., Budin, N., Shpak, B., Artyomov, M.N., and Sergushichev, A. (2021). Fast gene set enrichment analysis. bioRxiv, 060012. 10.1101/060012.

120. Blum, M., Andreeva, A., Florentino, L.C., Chuguransky, S.R., Grego, T., Hobbs, E., Pinto, B.L., Orr, A., Paysan-Lafosse, T., Ponamareva, I., et al. (2025). InterPro: the protein sequence classification resource in 2025. Nucleic Acids Res 53, D444–D456. 10.1093/nar/gkae1082.

121. Jones, P., Binns, D., Chang, H.Y., Fraser, M., Li, W., McAnulla, C., McWilliam, H., Maslen, J., Mitchell, A., Nuka, G., et al. (2014). InterProScan 5: genome-scale protein function classification. Bioinformatics 30, 1236–1240. 10.1093/bioinformatics/btu031.

122. Meszaros, B., Erdos, G., and Dosztanyi, Z. (2018). IUPred2A: context-dependent prediction of protein disorder as a function of redox state and protein binding. Nucleic Acids Res 46, W329–W337. 10.1093/nar/gky384.

123. Gurdon, J.B., and Wickens, M.P. (1983). The use of Xenopus oocytes for the expression of cloned genes. Methods Enzymol 101, 370–386. 10.1016/0076-6879(83)01028-9.

124. Brown, D.D., and Gurdon, J.B. (1964). Absence of Ribosomal Rna Synthesis in the Anucleolate Mutant of Xenopus Laevis. Proc Natl Acad Sci U S A 51, 139–146. 10.1073/pnas.51.1.139.

125. Brown, D.D., and Littna, E. (1964). Rna Synthesis during the Development of Xenopus Laevis, the South African Clawed Toad. J Mol Biol 8, 669–687. 10.1016/s0022-2836(64)80116-9.

126. Chase, J.W., and Dawid, I.B. (1972). Biogenesis of mitochondria during Xenopus laevis development. Dev Biol 27, 504–518. 10.1016/0012-1606(72)90189-3.

127. Zalokar, M., and Erk, I. (1976). Division and migration of nuclei during early embryogenesis of Drosophila melanogaster. Journal de Microscopie et de Biologie Cellulaire 25, 97–106.

128. Adams, M.D., Celniker, S.E., Holt, R.A., Evans, C.A., Gocayne, J.D., Amanatides, P.G., Scherer, S.E., Li, P.W., Hoskins, R.A., Galle, R.F., et al. (2000). The genome sequence of Drosophila melanogaster. Science 287, 2185–2195. 10.1126/science.287.5461.2185.

129. Salvador-Martinez, I., Grillo, M., Averof, M., and Telford, M.J. (2019). Is it possible to reconstruct an accurate cell lineage using CRISPR recorders? Elife 8. 10.7554/eLife.40292.

130. Smith, A.V., and Orr-Weaver, T.L. (1991). The regulation of the cell cycle during Drosophila embryogenesis: the transition to polyteny. Development 112, 997–1008. 10.1242/dev.112.4.997.

131. Anderson, K.V., and Lengyel, J.A. (1979). Rates of synthesis of major classes of RNA in Drosophila embryos. Dev Biol 70, 217–231. 10.1016/0012-1606(79)90018-6.

132. Becker, K., Bluhm, A., Casas-Vila, N., Dinges, N., Dejung, M., Sayols, S., Kreutz, C., Roignant, J.Y., Butter, F., and Legewie, S. (2018). Quantifying post-transcriptional regulation in the development of Drosophila melanogaster. Nat Commun 9, 4970. 10.1038/s41467-018-07455-9.

133. Deutsch, E.W., Bandeira, N., Perez-Riverol, Y., Sharma, V., Carver, J.J., Mendoza, L., Kundu, D.J., Wang, S., Bandla, C., Kamatchinathan, S., et al. (2023). The ProteomeXchange consortium at 10 years: 2023 update. Nucleic Acids Res 51, D1539–D1548. 10.1093/nar/gkac1040.

134. Perez-Riverol, Y., Bandla, C., Kundu, D.J., Kamatchinathan, S., Bai, J., Hewapathirana, S., John, N.S., Prakash, A., Walzer, M., Wang, S., and Vizcaino, J.A. (2025). The PRIDE database at 20 years: 2025 update. Nucleic Acids Res 53, D543–D553. 10.1093/nar/gkae1011.

