## Supplementary information for "Proteome-wide quantification of protein turnover in frog and fly embryos reveals divergent strategies of maternal inheritance"

**Table of contents**

- Figure S1. Wide isolation normalization and identification of stable proteins for frog turnover quantification.
- Figure S2. Labeling dynamics of individual free amino acids during frog 2-cell ^18^O experiment.
- Figure S3. Significance thresholds and model selection across experiments.
- Figure S4. Coordinated turnover of homeolog pairs.
- Figure S5. Cell-cycle-coupled turnover is a small fraction of proteome turnover.
- Figure S6. Labeling dynamics of individual free amino acids during frog gastrulation ^18^O experiment.
- Figure S7. Reproducibility and precision limits of the frog gastrulation 18O experiment
- Figure S8. Labeling dynamics of individual free amino acids during fly blastoderm ^18^O experiment.
- Figure S9. Peptide light-synthesis probability for the fly ^18^O time course.
- Figure S10. Normalization of the fly ^18^O turnover dataset.
- Figure S11. Uric acid accumulation during fly embryogenesis.
- Supplementary Methods
- Description of Additional Supplementary Files

**Supplementary Figures**

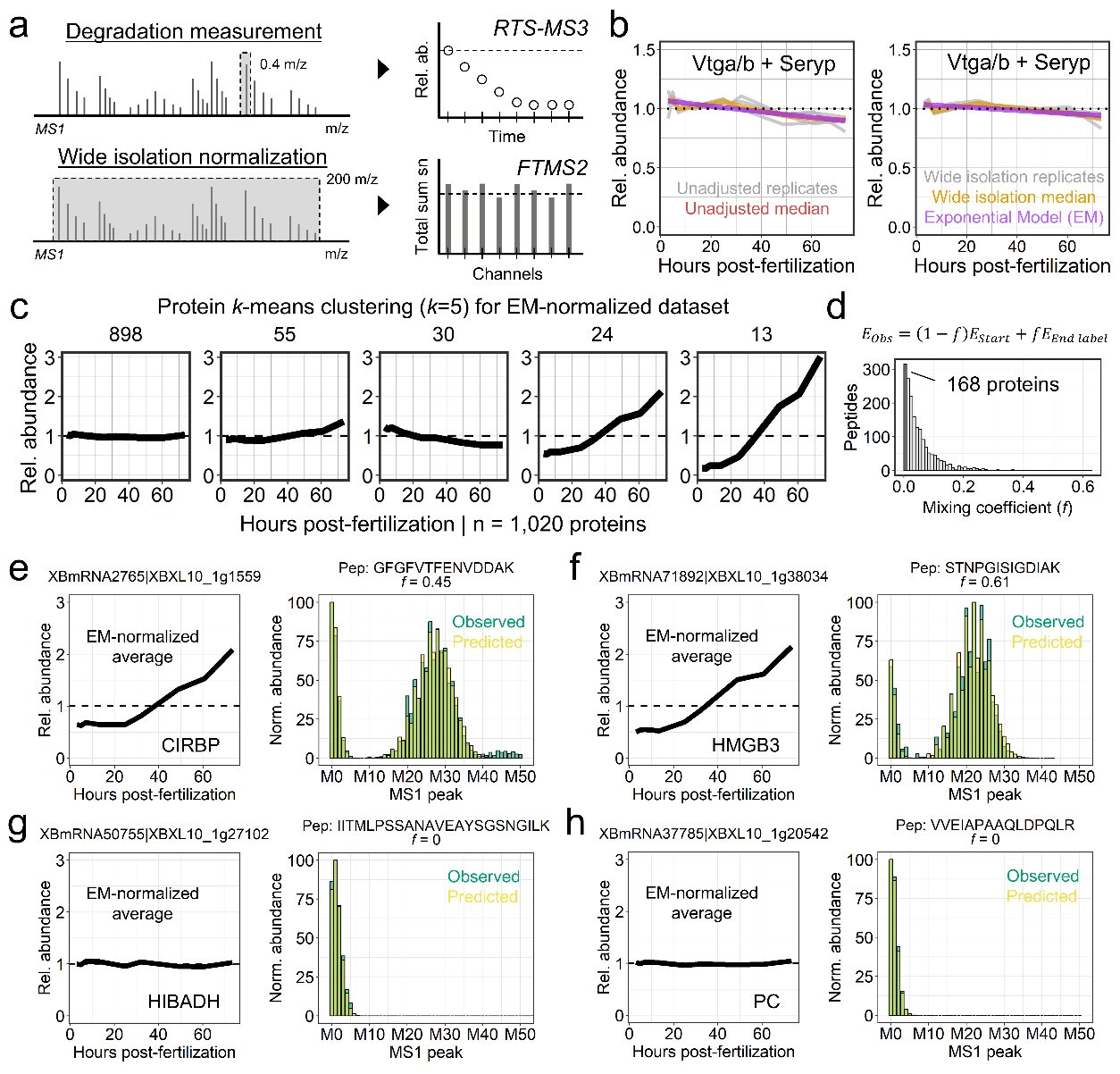

**Supplemental Figure 1.** **Wide isolation normalization and identification of stable proteins for frog turnover quantification.**

**(a)** Strategy for wide isolation normalization. Degradation measurements isolate a narrow precursor window (0.4 m/z) to quantify peptide isotopic decay over time. To assess channel balancing across multiplexed samples, we performed wide isolation MS1 scans (window of 200 m/z), capturing all nearby precursors and enabling quantification of total reporter ion signal across channels. **(b)** Estimating global abundance changes from yolk proteins. In standard *X. laevis* proteomics workflows, abundant yolk proteins are removed by centrifugation to increase proteomic depth. To obtain an unperturbed population of stable proteins, embryos were instead processed without yolk removal, producing a dataset dominated by stable yolk proteins. Wide isolation normalization reveals a systematic decrease in yolk abundance over developmental time. An exponential model (EM) was fitted to this decay and used as a global correction factor for the dataset. **(c)** Protein *k*-means clustering of the EM-normalized dataset. Following correction, proteins exhibiting less than 20% relative abundance change across the time course were selected as a candidate set of stable proteins. **(d)** Modeling isotopic labeling at the final ^18^O labeling time point. To determine whether candidate stable proteins incorporated heavy isotopes through new synthesis or active degradation, label-free acquisition was used to measure MS1 isotopic shifts at the final ^18^O timepoint. Theoretical steady-state isotopic envelopes were generated by convolving the labeling probabilities of each amino acid sequence (see Methods). Observed peptide envelopes ($E_{Obs}$) were then fitted as mixtures of unlabeled ($E_{start}$) and fully labeled ($E_{end label}$) states to extract a mixing coefficient ($f$), representing the fraction of heavy isotope incorporation. Stable candidate proteins with minimal incorporation ($f$ < 0.01) were finalized as the reference stable protein set (n=169) used to normalize all subsequent yolk-depleted datasets. **(e-h)** Representative protein abundance trajectories (left) and final-timepoint isotopic envelopes (right). **(e-f)** Examples of newly synthesized proteins (CIRBP, HMGB3) increasing in abundance. These proteins exhibit substantial heavy isotope incorporation, indicated by high mixing coefficients ($f$). The close alignment between the observed (green) and predicted (yellow) peaks provides confidence that the lack of heavy isotopologues observed in our stable protein set represents true absence of labeling rather than a failure of detection. **(g-h)** Examples of proteins from the reference stable set (HIBADH, PC). These exhibit flat abundance profiles in the EM-normalized data and no detectable heavy isotope incorporation, confirming their stability and validating their use for downstream dataset normalization.

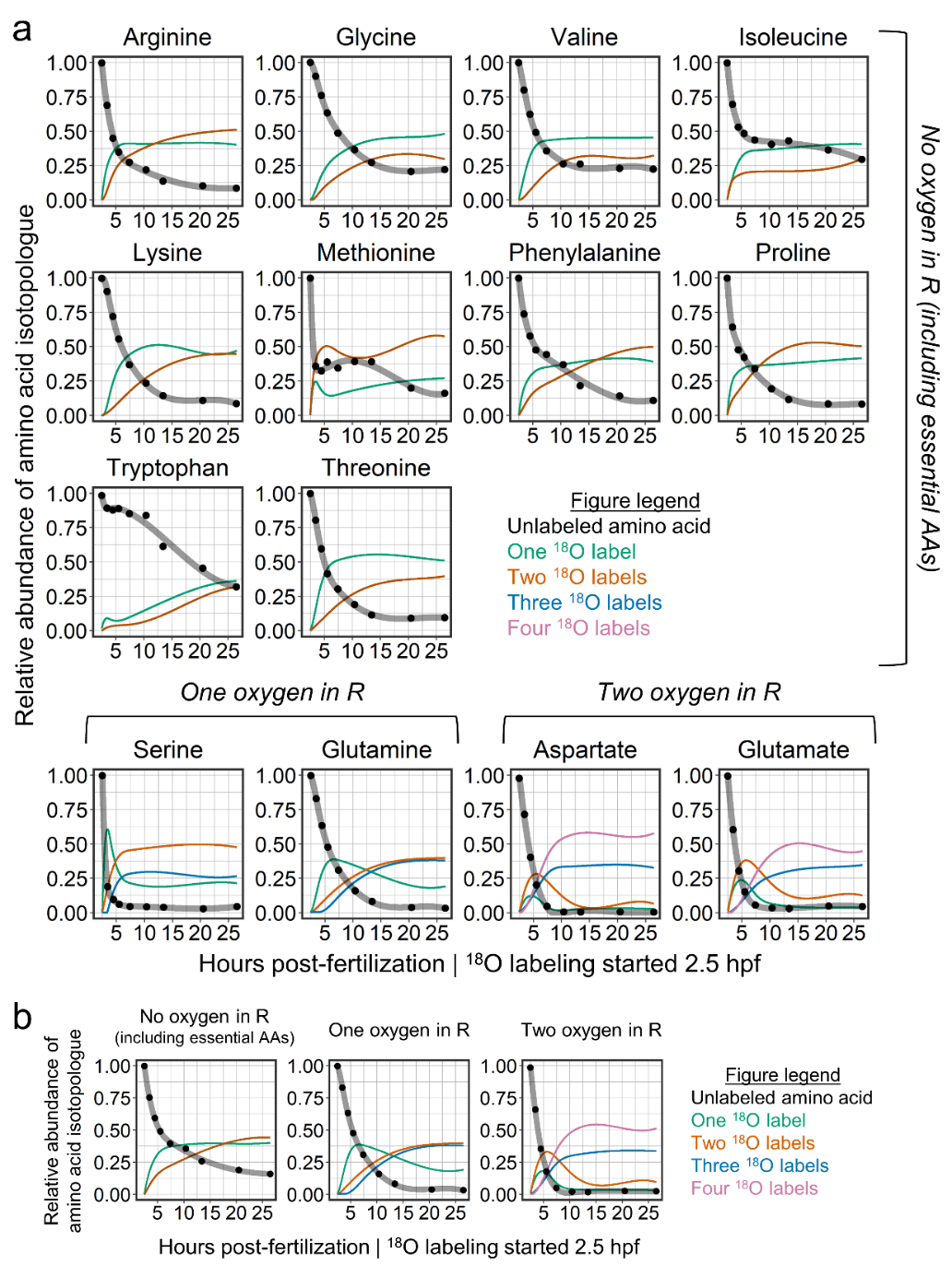

**Supplemental Figure 2. Labeling dynamics of individual free amino acids during frog 2-cell ^18^O experiment.**

**(a)** Relative abundances of unlabeled and ^18^O-labeled isotopologues were measured for individual amino acids following transfer of Xenopus embryos to H_2_^18^O at 2.5 hours post-fertilization. Points represent measured isotopologue fractions and lines indicate a cubic spline regression used to model precursor labeling dynamics. Amino acids are grouped according to the number of oxygen atoms present in their side chains (R groups), which determines the maximum number of ^18^O atoms that can be incorporated during labeling. Colored curves represent isotopologues containing increasing numbers of ^18^O atoms. These measurements are used to calculate the time-dependent probability of synthesizing light peptides used for downstream protein turnover modeling. **(b)** For amino acids not directly measured, precursor labeling dynamics were imputed using the median trajectory of their respective oxygen-count group.

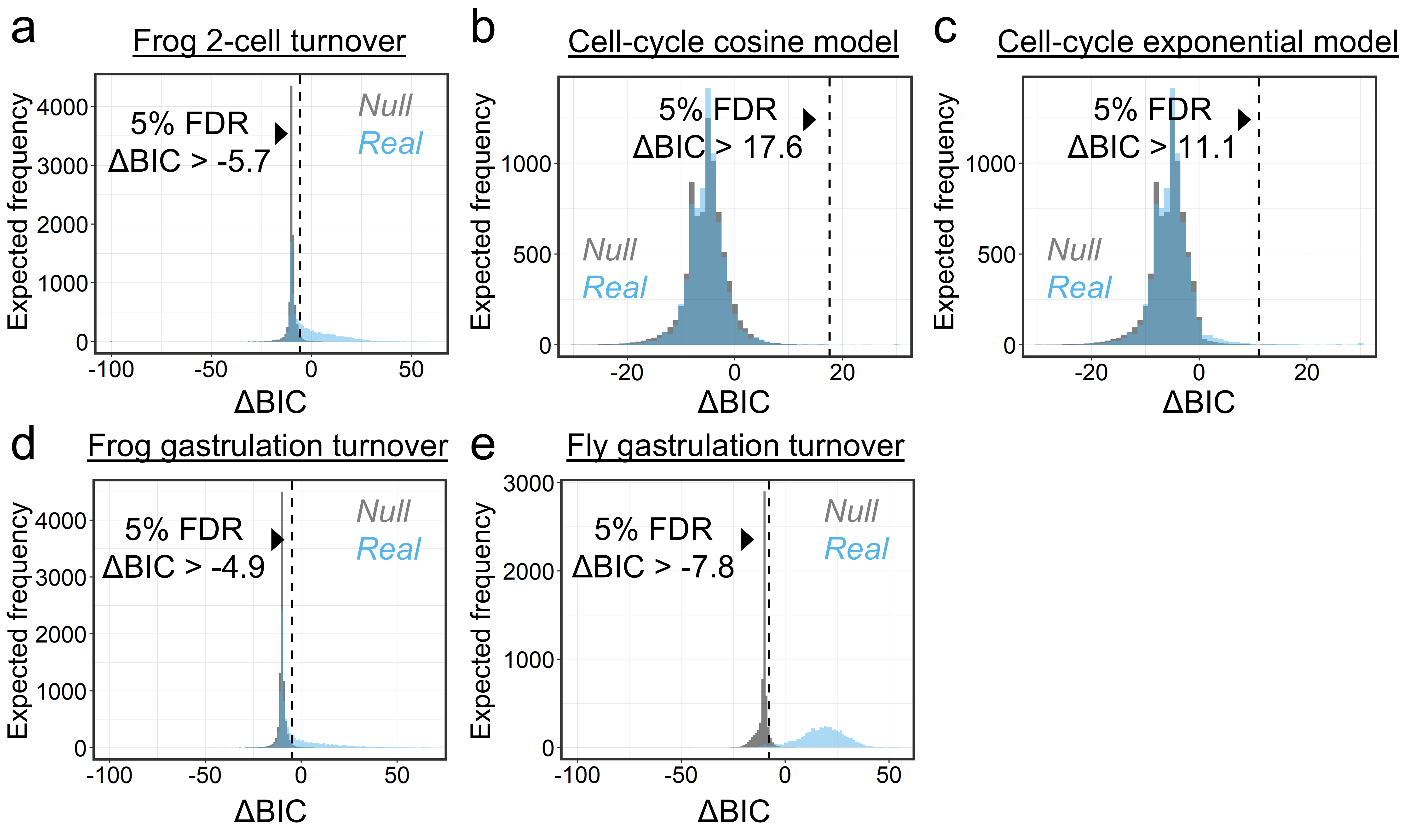

**Supplemental Figure 3. Significance thresholds and model selection across experiments.**

**(a)** Empirical null calibration of degradation classification for the Xenopus 2-cell ^18^O experiment. Fits were compared to a no-degradation null by Bayesian Information Criterion (BIC). Null distributions were generated by randomizing time points. Because randomized fits almost never favored degradation, the ΔBIC threshold that would control the false-discovery rate fell below zero, which would classify proteins as degrading without positive evidence. We therefore applied the conventional ΔBIC > 2 as a more stringent, positive-evidence threshold for cutoffs that fell below zero. **(b, c)** Model selection for the Xenopus egg activation time course (Figure S5). Each protein was tested against a cosine model for cell-cycle oscillation **(b)** and an exponential model for directional change **(c)** against an empirical null. **(d)** Null calibration for the Xenopus gastrulation experiment, as in (a). **(e)** Null calibration for the *D. melanogaster* experiment, as in **(a)**.

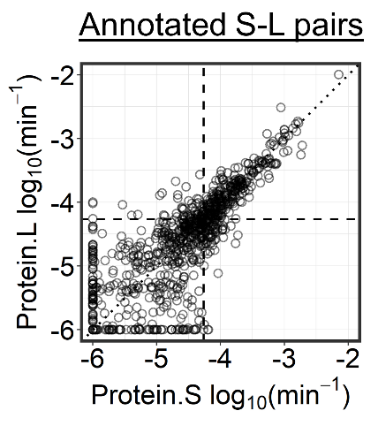

**Supplementary Figure 4. Coordinated turnover of homeolog pairs.**

*X. laevis* is an allotetraploid formed by the fusion of two frog genomes ~17 million years ago, and most genes are retained as short (S) and long (L) homeolog pairs. Degradation rate constants (k_d_) for annotated S-L pairs are plotted against each other, with the dotted line marking y = x. Dashed lines mark the precision limit (212 hours) of the 2-cell experiment on which these data are based. Rates are very similar within a pair, indicating that turnover is set largely by intrinsic protein features shared between homeologs rather than by regulatory differences between the copies.

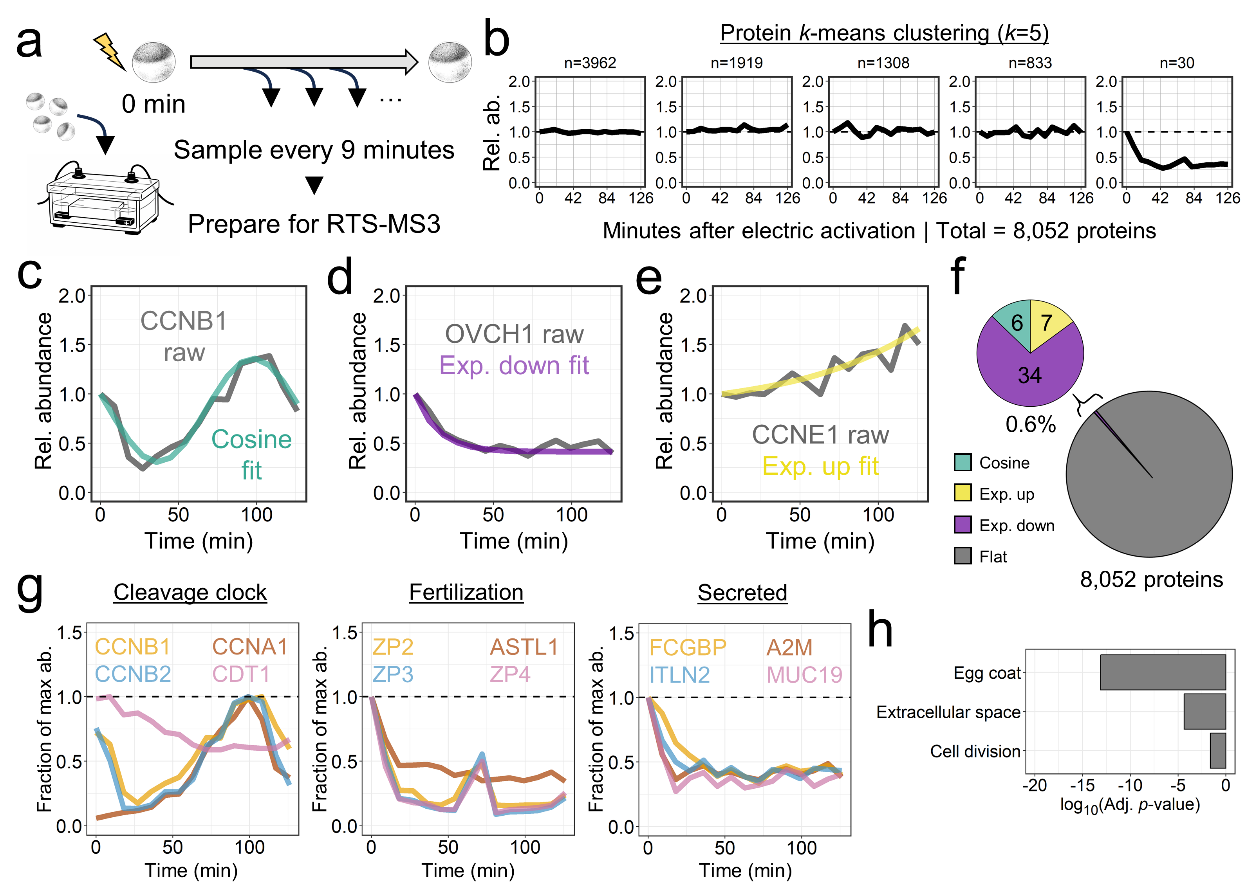

**Supplemental Figure 5. Cell-cycle-coupled turnover is a small fraction of proteome turnover.**

**(a)** Experimental design. Unfertilized Xenopus eggs were electrically activated and sampled every 9 minutes over a two-hour period at 16°C, spanning the first cell cycle. Samples were quantified using a multiplexed RTS-MS3 pipeline. **(b)** Global abundance stability. Protein *k*-means clustering (*k* = 5) of the 8,052 quantified proteins shows that most of the proteome has no measurable change across the first cell cycle. **(c–e)** Kinetic modeling of dynamic subpopulations. While the bulk proteome is flat, specific subsets exhibit highly dynamic trajectories. All proteins were tested against two possible models: a cosine-based model to capture oscillatory proteins and an exponential model to capture net protein loss or accumulation. **(c)** Known oscillatory cell-cycle drivers (e.g., CCNB1) with the cosine model. A representative cell-cycle period was first empirically determined from known oscillatory proteins, allowing a two-parameter fit for amplitude and phase (green cosine fit). **(d, e)** Proteins exhibiting directional abundance changes following activation were fit with a three-parameter exponential model, capturing active clearance (**d**; e.g., OVCH1, purple) or rapid accumulation (**e**; e.g., CCNE1, yellow). **(f)** Global kinetic classification. Statistical model selection reveals that only 47 proteins (~0.6%) undergo dynamic change: 6 oscillating, 7 accumulating, 34 declining. **(g)** Functional trajectories of dynamic networks. Decreasing and oscillating proteins segregate into distinct biological behaviors: the cleavage clock (canonical cell-cycle regulators); egg-activation response proteins (structural ZP proteins and their paired proteases); and other actively secreted factors (e.g., MUC19, A2M). The rapid loss of abundance in the latter two groups reflects their established ejection or secretion into the extracellular space following activation, rather than strict intracellular degradation. **(h)** Functional enrichment of significant proteins. Gene Ontology (GO) enrichment of the dynamically changing subpopulation highlights known pathways related to cell division and extracellular structures (egg coat, extracellular space).

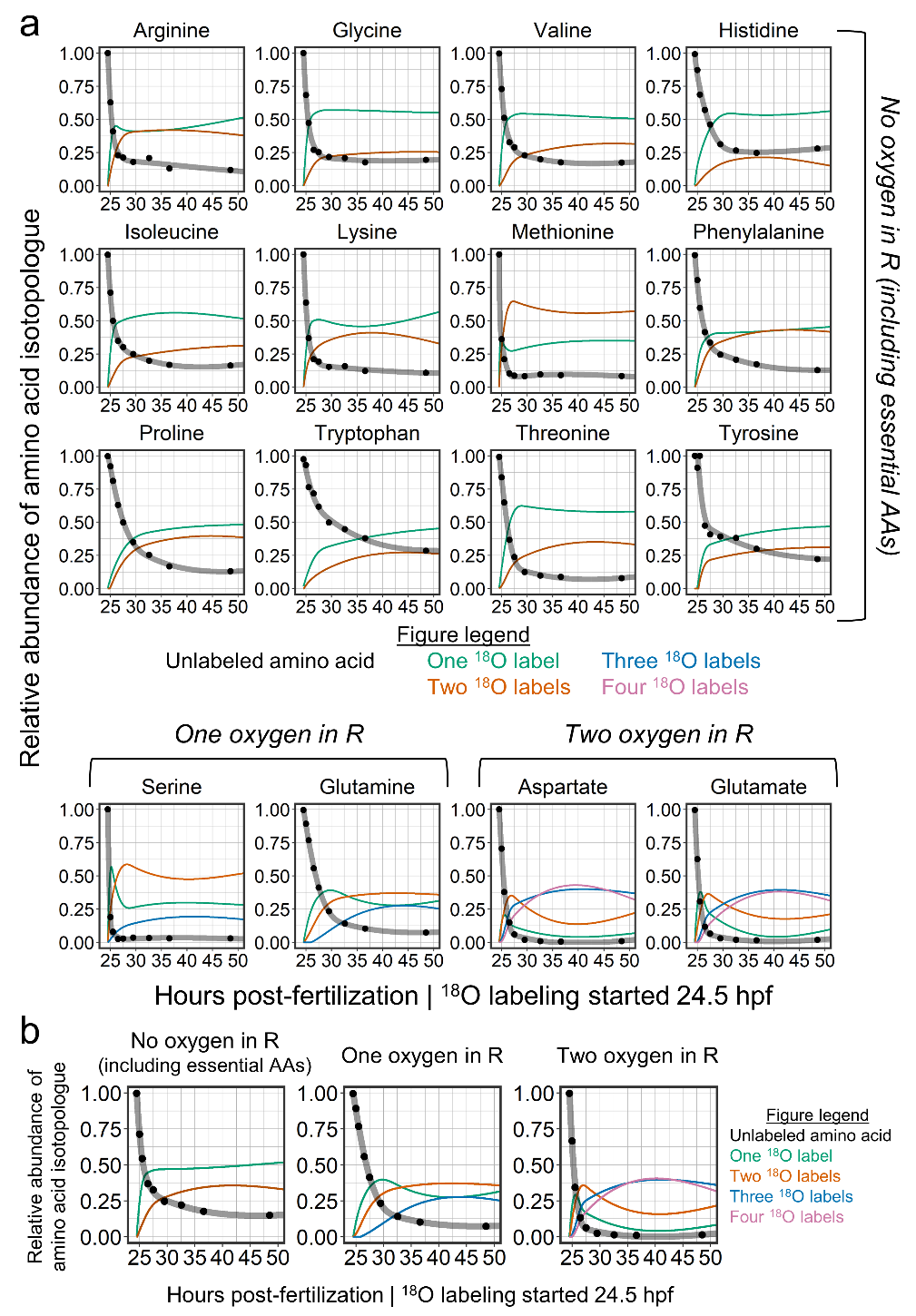

**Supplemental Figure 6. Labeling dynamics of individual free amino acids during frog gastrulation ^18^O experiment.**

**(a)** Relative abundances of unlabeled and ^18^O-labeled isotopologues were measured for individual amino acids following transfer of Xenopus embryos to H_2_^18^O at 24.5 hours post-fertilization. Points represent measured isotopologue fractions, and lines indicate a cubic spline regression used to model precursor labeling dynamics. Amino acids are grouped according to the number of oxygen atoms present in their side chains (R groups), which determines the maximum number of ^18^O atoms that can be incorporated during labeling. Colored curves represent isotopologues containing increasing numbers of ^18^O atoms. These measurements are used to calculate the time-dependent probability of synthesizing light peptides used for downstream protein turnover modeling. **(b)** For amino acids not directly measured, precursor labeling dynamics were imputed using the median trajectory of their respective oxygen-count group.

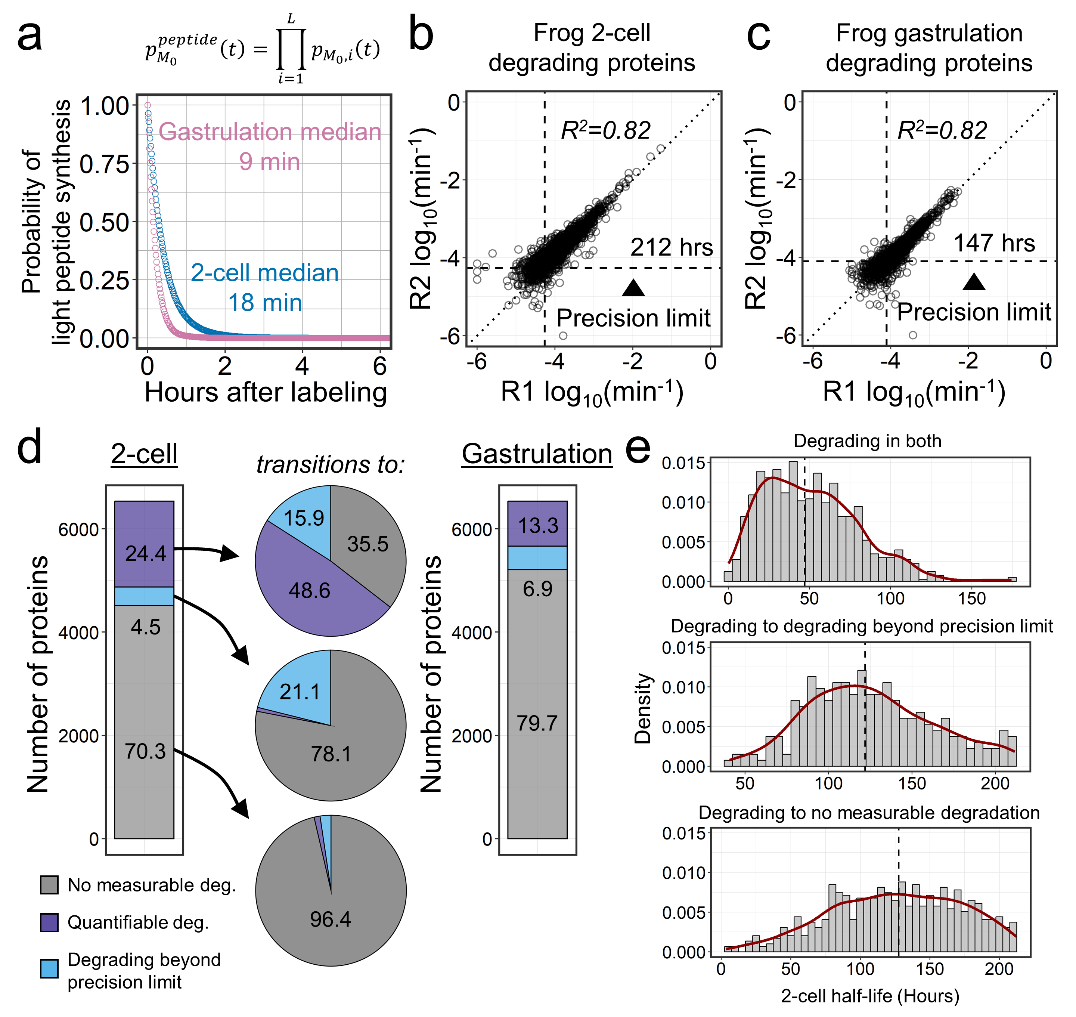

**Supplemental Figure 7. Reproducibility and precision limits of the frog gastrulation ^18^O experiment.**

**(a)** Amino acid pool equilibration. The probability of synthesizing light peptides is plotted over time following isotopic labeling. Amino acids equilibrate roughly twice as fast during gastrulation (median 9 minutes) compared to the 2-cell stage (median 18 minutes).

**(b-c)** Inter-replicate agreement of fitted degradation rate constants. Because both ^18^O courses end at the same developmental stage, the gastrulation window is shorter and resolves a correspondingly shorter maximum half-life (147 hours at gastrulation versus 212 hours at the 2-cell stage). Among the degrading proteins, the gastrulation replicates agree as well as at the 2-cell stage (R^2^ = 0.82; **b**, 2-cell; **c**, gastrulation).

**(d)** Kinetic classifications and precision limits. A flow diagram tracking protein classification from the 2-cell stage into gastrulation. While a significant portion of the quantifiable proteome appears to shift toward no measurable degradation, this is largely driven by the shorter gastrulation experimental window, which proportionally reduces the maximum quantifiable half-life. Consequently, many proteins shifting to "no measurable degradation" or "beyond precision" reflect a systematic loss of analytical resolution for slow-degrading proteins rather than a true biological slowdown of turnover.

**(e)** Initial half-life predicts late-stage classification. Density plots mapping the initial 2-cell half-lives of proteins, grouped by their subsequent kinetic classification during gastrulation. Proteins that maintain quantifiable degradation across both stages (top) are restricted to those with the shortest initial half-lives. Conversely, proteins that transition to "beyond precision" (middle) or "no measurable degradation" (bottom) originate from populations with significantly longer 2-cell half-lives. This rightward shift implies that the apparent global slowdown is primarily driven by experimental boundary limits, suggesting that the bulk proteome neither systematically accelerates nor decelerates during gastrulation.

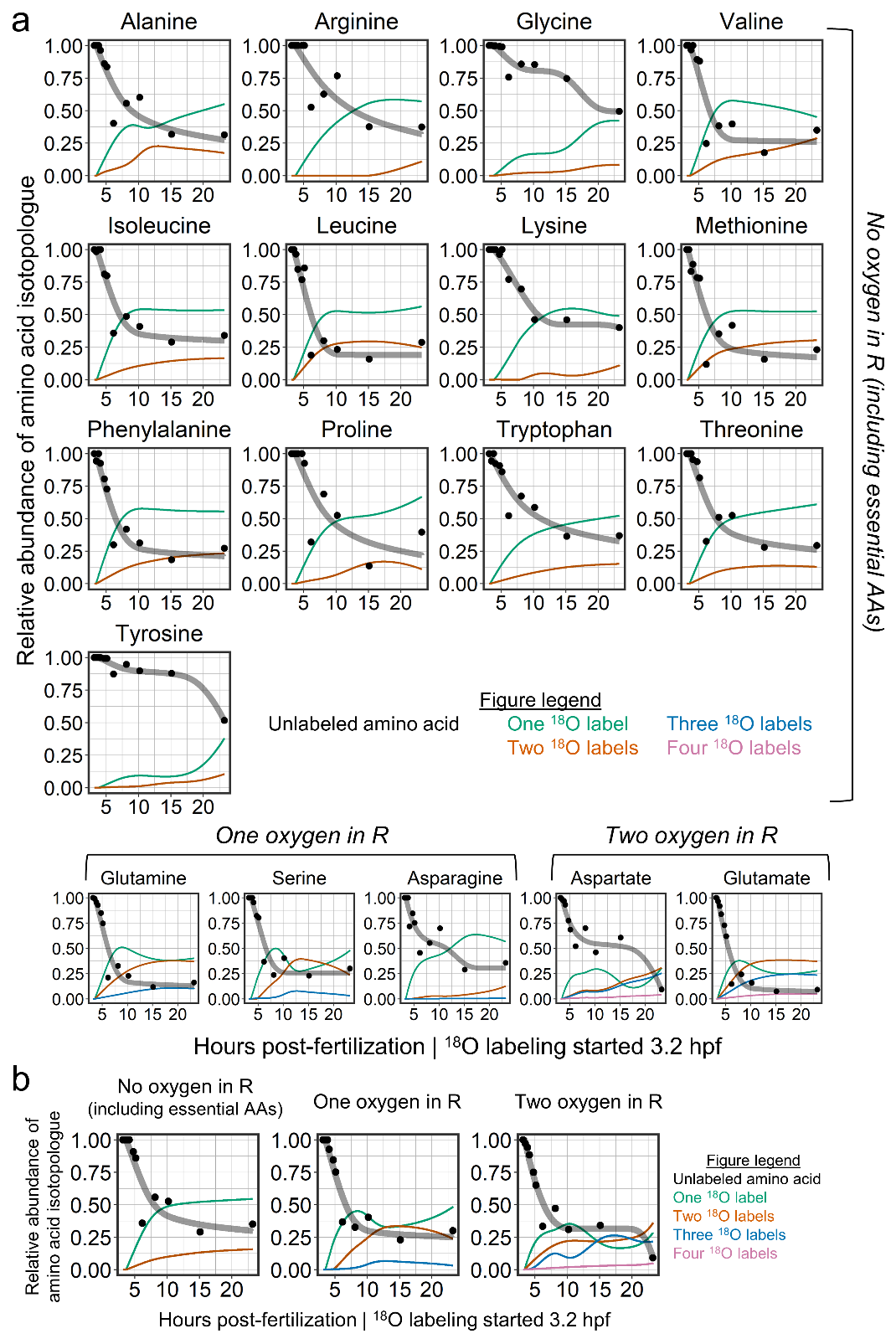

**Supplemental Figure 8. Labeling dynamics of individual free amino acids during fly blastoderm ^18^O experiment.**

**(a)** Relative abundances of unlabeled and ^18^O-labeled isotopologues were measured for individual amino acids following transfer of dechorionated Drosophila embryos to an ^18^O-soaked paper towel at 3.1 hours post-fertilization. Points represent measured isotopologue fractions, and lines indicate the fitted labeling curves used to model precursor labeling dynamics. Amino acids are grouped according to the number of oxygen atoms present in their side chains (R groups), which determines the maximum number of ^18^O atoms that can be incorporated during labeling. Colored curves represent isotopologues containing increasing numbers of ^18^O atoms. These measurements are used to calculate the time-dependent probability of synthesizing light peptides used for downstream protein turnover modeling. **(b)** For amino acids not directly measured, precursor labeling dynamics were imputed using the median trajectory of their respective oxygen-count group.

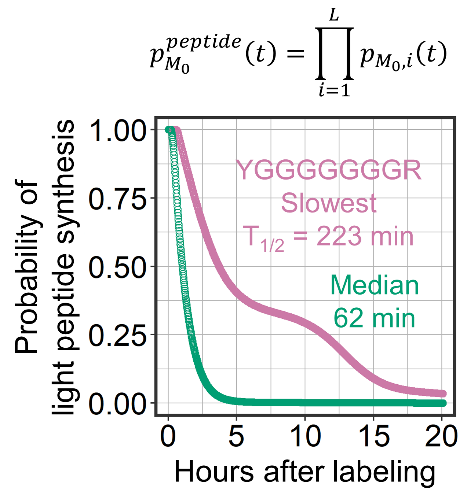

**Supplemental Figure 9. Peptide light-synthesis probability for the fly ^18^O time course.**

From the measured free amino acid labeling curves (Figure S8), we calculated the time-dependent probability of newly synthesized peptides that are fully unlabeled (light) for each detected sequence. The median tryptic peptide halves within 62 minutes of label exposure; the slowest-labeled peptide observed halves within 223 minutes.

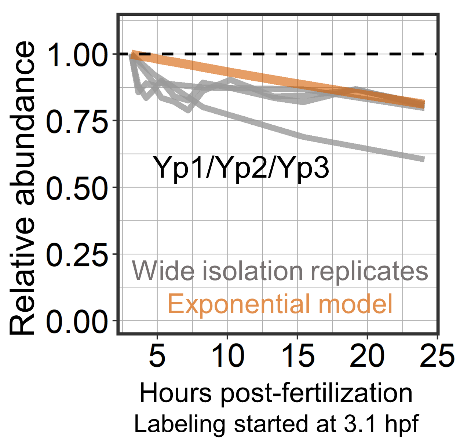

**Supplemental Figure 10. Normalization of the fly ^18^O turnover dataset.**

Wide-isolation scans acquired alongside the narrow-isolation degradation scans provide a loading control independent of protein identity, the same strategy used for Xenopus (Figure S1). Grey lines show median yolk protein abundance (Yp1/Yp2/Yp3) across the four time courses (two control, two ¹⁸O), with an exponential decay model fitted in orange. Because Drosophila preparation did not include the centrifugation step to remove yolk that was used for Xenopus, no stable reference set was generated, so a correction factor derived from this exponential yolk-decay model was applied directly to the turnover dataset.

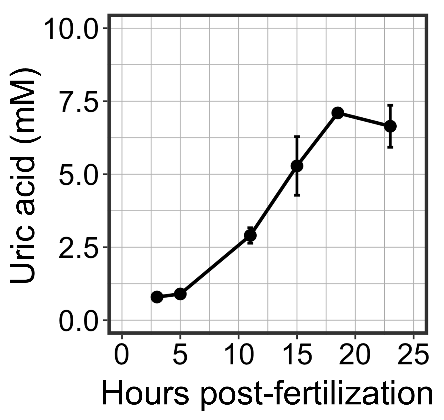

**Supplemental Figure 11. Uric acid accumulation during fly embryogenesis.**

Intra-embryonic uric acid concentration in Drosophila as a function of time after fertilization, measured by enzymatic assay (Methods). Lysate measurements were corrected to intra-embryonic concentrations using the counted number of embryos per sample and the corresponding embryo volume. Levels rise roughly 8-fold, from ~0.8 mM at ~3 hours post-fertilization to a plateau of ~6.6 mM by ~18 hours. Points are means and error bars are SEM across three biological replicates.

**Supplementary Methods**

**Conversion of free amino acid labeling to peptide light synthesis probability**

The kinetic model for protein turnover requires, as an input, the time-dependent probability that a newly synthesized peptide is fully unlabeled (the *light synthesis probability*). This quantity cannot be measured directly. Instead, metabolomics provides the isotopologue distribution of the free amino acid pool, i.e. the fraction of each free amino acid present as its M_0_, M_2_, … , M_n_ isotopologue following the switch to ^18^O-labeled water. Here, we derive the relationship between the free amino acid M_0_ fraction and the peptide light synthesis probability used in the turnover model.

Notation

For a given amino acid, we define two independent labeling probabilities:

- $p$ : the probability that a backbone carboxyl oxygen is a ^16^O atom
- $r$ : the probability that a given side-chain oxygen is a ^16^O atom (defined per side-chain oxygen, so that an amino acid with two labelable side-chain oxygens contributes a factor of r^2^)

and their complements:

- $q = 1 - p$ : the probability that a backbone carboxyl oxygen is a ^18^O atom
- $s=1-r$ : the probability that the side chain carries a ^18^O atom

Each amino acid carries at least two backbone carboxyl oxygens. During ^18^O labeling, each oxygen position is independently light with probability $p$. The measured quantity from metabolomics, $a_{n}$, is the fraction of the free amino acid in each $M_{n}$ isotopologue.

Consider an amino acid with no labelable side chain oxygen. Here, the $M_{0}$ isotopologue requires both backbone oxygens to be light. Because the two oxygens are independent,

$$a_{0}={rp}^{2}=\left( 1 \right)p^{2}=p^{2}$$

More generally, $a_{0}$ is the $M_{0}$ fraction of the free amino acid as reported by metabolomics, and we wish to relate it to the probability that the amino acid contributes only light atoms when incorporated into a peptide.

Light synthesis probability for a single amino acid

We require the probability that, when an amino acid is incorporated into a peptide, it contributes no ^18^O atoms. However, isotopic labeling is measured stoichiometrically via metabolomics, without positional resolution, so position-specific labeling probabilities cannot be assigned from the data. We therefore adopt the simplifying assumption that all labelable positions on an amino acid label at comparable rates ($r=p$). Under this assumption, the light synthesis probability depends only on the number of labelable oxygen positions an amino acid carries.

We further account for side-chain oxygens that lack an active labeling route. Whether a side-chain oxygen acquires a ^18^O label depends on the amino acid's metabolic origin in this system rather than on its chemical group alone. Serine, synthesized *de novo*, labels its hydroxyl oxygen, whereas the chemically similar hydroxyls of threonine and tyrosine do not (Figure S2; Figure S6; Figure S8). Threonine is essential and therefore has no biosynthetic route to label its side chain. Tyrosine is synthesized by hydroxylation of phenylalanine, which is essential and therefore unlabeled in this system; tyrosine consequently has no route to label its side chain and behaves as an amino acid with no labelable side-chain oxygen. For these essential amino acids, we treat the side-chain oxygen as non-labelable and assign the corresponding lower oxygen count. This yields three cases.

Case 1: No labelable side-chain oxygen

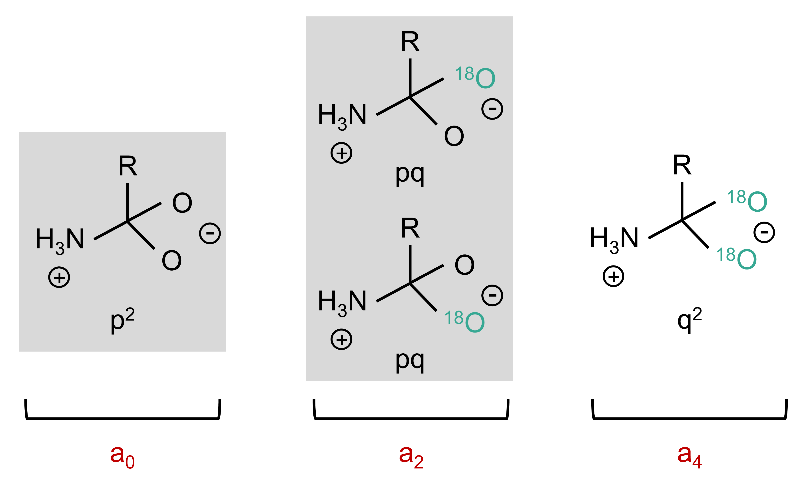

The only labelable positions are the two backbone carboxyl oxygens. Upon peptide bond formation, one carboxyl oxygen is lost, so the residue retains a single backbone oxygen in the peptide. The retained oxygen is light with certainty when both backbone oxygens were light (the M_0_ case), but only with probability one-half when exactly one was light (the M_2_ case), since either oxygen is equally likely to be the one lost. The probability that the incorporated residue is light is therefore the sum of these contributions (highlighted in grey), weighted by the chance that the retained oxygen is the light one:

$$p_{M_{o}}=p_{M_{o}|a_{o}}+\frac{1}{2}p_{M_{o}|a_{2}}$$

Considering each individual isotopologues contribution

$$p_{M_{o}|a_{o}}=p^{2}$$

$$p_{M_{o}|a_{2}}=2pq$$

$$p_{M_{o}}=p^{2}+pq$$

Expression in terms of $p$

$$p_{M_{o}}=p^{2}+p(1-p)$$

$$p_{M_{o}}=p$$

Since the measured $M_{0}$ fraction is $a_{0}=p^{2}$, we have:

$$p_{M_{o}}=\sqrt{a_{o}}=a_{0}^{1/2}$$

Case 2: One labelable side-chain oxygen

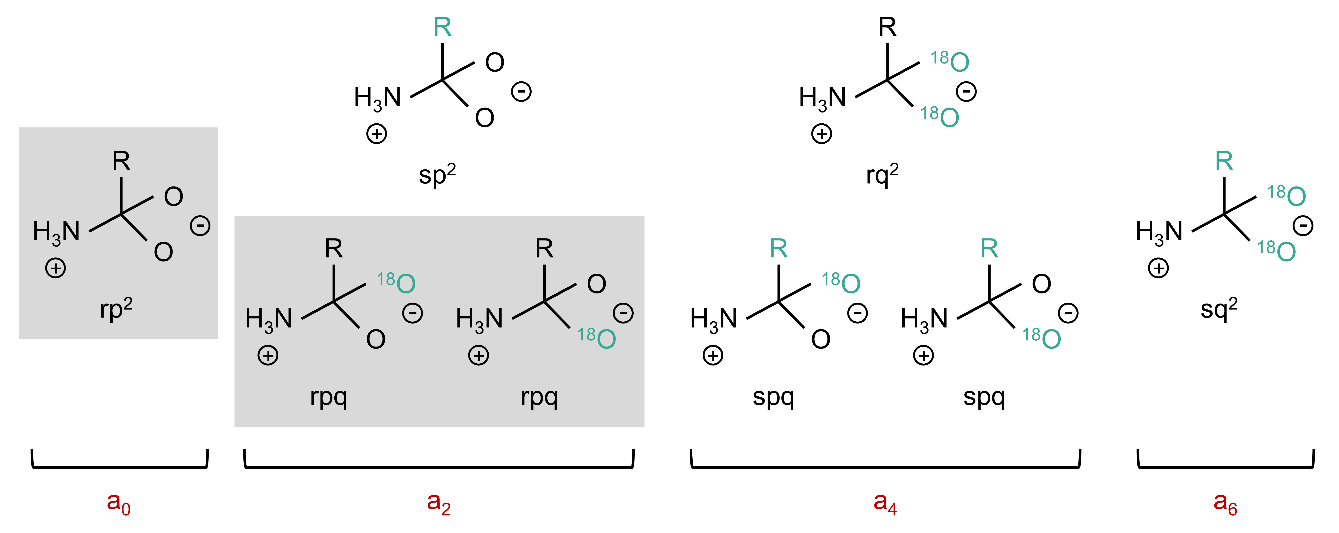

With one additional labelable oxygen on the side chain, the residue carries two relevant positions into the peptide: the single retained backbone oxygen plus the side-chain oxygen, both of which must be light (highlighted in grey). As in Case 1, the one-half factor accounts for which backbone oxygen is retained after peptide bond formation. The probability that the incorporated residue is light is

$$p_{M_{o}}=p_{M_{o}|a_{o}}+\frac{1}{2}p_{M_{o}|a_{2}}$$

Considering each individual isotopologues contribution

$$p_{M_{o}|a_{o}}={rp}^{2}$$

$$p_{M_{o}|a_{2}}=2rpq$$

$$p_{M_{o}}={rp}^{2}+rpq$$

Expression in terms of $p$

$$p_{M_{o}}={rp}^{2}+rp(1-p)$$

$$p_{M_{o}}=rp$$

Under the assumption that the side chain and backbone oxygen label at the same rate ($r=p$), the measured $M_{0}$ fraction is $a_{0}={rp}^{2}=p^{3}$, we have:

$$p_{M_{o}}=p^{2}=a_{0}^{2/3}$$

Case 3: Two labelable side-chain oxygens

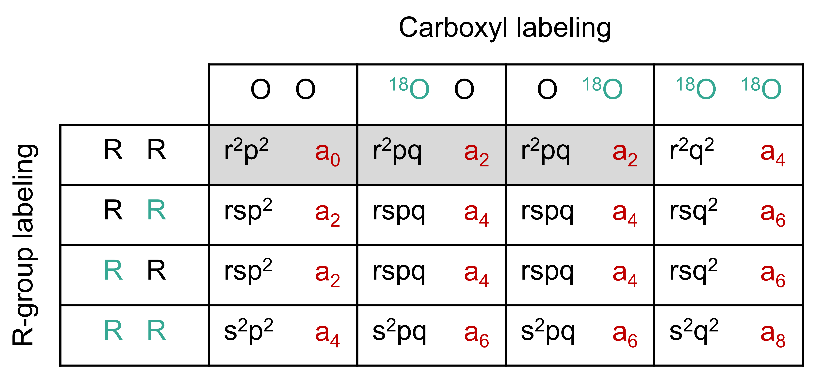

With two side-chain oxygens, the residue carries three relevant positions into the peptide: the retained backbone oxygen plus the two side-chain oxygens, all of which must be light (highlighted in grey). As before, the one-half factor accounts for which backbone oxygen is retained after peptide bond formation. The light-synthesis probability is

$$p_{M_{o}}=p_{M_{o}|a_{o}}+\frac{1}{2}p_{M_{o}|a_{2}}$$

Considering each individual isotopologues contribution

$$p_{M_{o}|a_{o}}={r^{2}p}^{2}$$

$$p_{M_{o}|a_{2}}=2r^{2}pq$$

$$p_{M_{o}}={r^{2}p}^{2}+r^{2}pq$$

Expression in terms of $p$

$$p_{M_{o}}={r^{2}p}^{2}+r^{2}p(1-p)$$

$$p_{M_{o}}=r^{2}p$$

Under $r=p$, the measured $M_{0}$ fraction is $a_{0}={r^{2}p}^{2}=p^{4}$, giving:

$$p_{M_{o}}=p^{3}=a_{0}^{3/4}$$

General relationship

The three cases follow a single pattern. For an amino acid with *n* labelable oxygen positions relevant to peptide incorporation,

$$P_{M_{0}}=a_{0}^{n/(n+1)}$$

with *n* = 1, 2, 3 corresponding to Cases 1, 2, and 3 respectively. This expression converts the directly measured free amino acid $M_{0}$ fraction ($a_{0}$) into the per-residue light-synthesis probability required by the model, using only the oxygen content of each amino acid.

From single residues to whole peptides

A peptide contributes to its own M₀ peak only if every constituent residue is light. Because residues label independently, the peptide light-synthesis probability is the product of the per-residue probabilities across all residues in the peptide:

$$p_{M_{0}}^{peptide}(t)=\prod_{i=1}^{L} p_{M_{0},i}(t)$$

where the product runs over the *L* residues of the peptide and $p_{M_{0}.i}(t)$ is the time-dependent light-synthesis probability of residue *i*, computed from its measured free amino acid M₀ fraction via the appropriate power law above.

Fitting the precursor function

The peptide light-synthesis probability computed above is evaluated at each metabolomics time point. Because no labeling has occurred at the moment of the switch to ^18^O-water, a newly synthesized peptide is fully light with probability one at t = 0. The start is therefore fixed to 1 and the curve is fit to a single-exponential decay,

$$p_{M_{0}}^{peptide}(t)=e^{-k_{2}t}$$

where *k₂* is the effective precursor decay rate. The fitted function serves as the precursor labeling input $p_{L}\left( t \right)$ in the protein turnover model below.

Notes on implementation

A subset of amino acids were not directly quantified in the free amino acid metabolomics. For each such amino acid, we assigned it to the appropriate oxygen-count case based on its side-chain chemistry and metabolic labeling route, then substituted the median labeling trajectory of the measured amino acids within that same case grouping.

**Derivation of the protein turnover model**

We model the fully unlabeled ($M_{0}$) signal of each peptide following the switch to ^18^O-labeled water. Two processes change this signal over time. Existing protein, whether light or heavy, is removed by degradation at a first-order rate $k_{d}$. Newly synthesized protein is added at an effective rate $k_{T}$, but a newly made peptide is unlabeled only with the precursor light synthesis probability $p_{L}\left( t \right)$, which decays as the free amino acid pool incorporates ^18^O. The contribution of synthesis to the unlabeled signal is therefore $k_{T}p_{L}\left( t \right)$. These processes give the governing equation

$$\frac{dM_{0}}{dt}=k_{T}p_{L}\left( t \right)-k_{d}M_{0}$$

We fit two channels that share $k_{d}$ and $k_{T}$ and differ only in their precursor function. In the heavy (^18^O-labeled) channel the precursor labels over time, so the light synthesis probability decays; because no labeling has occurred at the moment of the switch, the probability is one at $t=0$, giving a single-exponential precursor

$$p_{L}\left( t \right)=e^{-k_{2}t}$$

where $k_{2}$ is the effective precursor decay rate, determined per peptide from the free amino acid labeling fit and treated here as a fixed input rather than a free parameter. In the unlabeled control channel, no ^18^O-water is present, so newly synthesized protein is always light and $p_{L}\left( t \right)=1$. The precursor labeling is determined empirically from the free amino acid metabolomics, so it enters the synthesis term as a measured input and no precursor flux model is required.

General solution

The governing equation is a linear first-order ordinary differential equation. Writing it in standard form and multiplying by the integrating factor $e^{k_{d}t}$,

$$\frac{dM_{0}}{dt}+k_{d}M_{0}=k_{T}p_{L}\left( t \right)$$

$$\frac{d}{dt}\left( M_{0}e^{k_{d}t} \right)=k_{T}p_{L}\left( t \right)e^{k_{d}t}$$

Integrating from 0 to $t$ and rearranging gives the general solution

$$M_{0}\left( t \right)=M_{0}\left( 0 \right)e^{-k_{d}t}+k_{T}e^{-k_{d}t}\int_{0}^{t} p_{L}\left( \tau\right)e^{k_{d}\tau}d\tau$$

The first term is the decay of the initial pool; the second is the accumulated contribution of synthesis. We now evaluate this for each channel.

Heavy channel

Substituting the heavy-channel precursor $p_{L}\left( \tau\right)=e^{-k_{2}\tau}$ and evaluating the integral,

$$\int_{0}^{t} e^{-k_{2}\tau}e^{k_{d}\tau}d\tau=\frac{e^{\left( k_{d}-k_{2} \right)t}-1}{k_{d}-k_{2}}$$

Multiplying by $k_{T}e^{-k_{d}t}$ and writing $m_{0,h}$ for the initial heavy-channel value $M_{0}\left( 0 \right)$ yields the heavy-channel model

$$M_{0}\left( t \right)=m_{0,h}e^{-k_{d}t}+\frac{k_{T}}{k_{d}-k_{2}}\left( e^{-k_{2}t}-e^{-k_{d}t} \right)$$

Control channel

For the control channel the precursor is constant, $p_{L}\left( \tau\right)=1$. The integral evaluates to $\frac{e^{k_{d}t}-1}{k_{d}}$, and multiplying by $k_{T}e^{-k_{d}t}$ gives, with $m_{0,n}$ the initial control-channel value,

$$M_{0}\left( t \right)=\frac{k_{T}}{k_{d}}+\left( m_{0,n}-\frac{k_{T}}{k_{d}} \right)e^{-k_{d}t}$$

As $t$ increases the control-channel signal approaches the steady-state value $\frac{k_{T}}{k_{d}}$, the ratio of synthesis to degradation. The control channel does not depend on the precursor decay rate, since its precursor is constant.

Fitted parameters and limiting behavior

For each peptide the two channels are fit jointly for four parameters: the initial channel values $m_{0,h}$ and $m_{0,n}$, the degradation rate $k_{d}$, and the effective synthesis rate $k_{T}$. The precursor decay rate $k_{2}$ is fixed per peptide as described above, and the precursor initial value is fixed to one. The degradation rate $k_{d}$ is the quantity of biological interest; $k_{T}$ is an anchor parameter that scales synthesis and is not interpreted directly, as it convolves transcription, translation, and abundance.

Because newly synthesized peptide is unlabeled only while the precursor light-synthesis probability remains appreciable, light-channel timepoints were retained up to and including the first timepoint at which this probability, $e^{-k_{2}t}$, had fallen to 0.1 or below. Beyond this point, newly synthesized peptide is essentially fully labeled, and synthesis contributes negligibly to the monoisotopic signal, which instead reflects decay of the pre-existing pool. These later timepoints were therefore excluded from the fit, as they do not inform $k_{T}$ and can bias the estimate. A minimum of two light-channel timepoints was retained per peptide.

The solution satisfies the expected boundary and limiting conditions. At $t=0$ both channels return their initial values $m_{0,h}$ and $m_{0,n}$. As $t$ increases the control channel approaches $\frac{k_{T}}{k_{d}}$ while the heavy channel approaches zero as the initial unlabeled pool is replaced by labeled protein. When $k_{T}=0$ the heavy channel reduces to pure first-order decay, $m_{0,h}e^{-k_{d}t}$.

**Cell-cycle dynamics in the activated egg**

Across the 15-time-point activation series (0 to 126 min in 9-minute steps), we first set the cell-cycle period by fitting a cosine model with a free period to the CCNB1.S/L peptides:

$$y\left( t \right)=A(cos( \frac{2\pi t}{T}+\varphi)+1)+(1-A)$$

where $A$is the amplitude, $\varphi$ the phase, and $T$ the period. The fitted period from these peptides was T ≈ 122 min, which we fixed for all subsequent fits.

Each protein was then compared against three models. The null was a flat model of constant abundance with no free parameters. Cell-cycle oscillation was described by the cosine model above with T fixed and amplitude and phase free. Directional changes were described by an exponential model:

$$y\left( t \right)=P_{0}e^{kt}+C$$

where $k$ > 0 corresponds to accumulation and $k$ < 0 to loss. We included a nonzero offset C because decreasing proteins did not fully deplete within the time course, and complete depletion was not expected. C therefore lets the model approach a residual level rather than forcing decay to zero.

For each protein, we took the better-fitting dynamic model, the lower BIC of the cosine and exponential, and used it for model selection.

**Description of Additional Supplementary Files**

**File:** Supplementary_Tables_Merge.xlsx

**Description:** Contains Supplementary Tables S1-7.

- **Table S1.** Proteins used to normalize the frog turnover data.
- **Table S2.** Protein turnover during frog development starting at the 2-cell stage.
- **Table S3.** Frog cell-cycle protein dynamics.
- **Table S4.** Protein turnover during frog gastrulation.
- **Table S5.** Protein turnover during fly gastrulation.
- **Table S6.** Absolute protein abundances in frog.
- **Table S7.** Absolute protein abundances in fly.
